# Conversion of natural cytokine receptors into orthogonal synthetic biosensors

**DOI:** 10.1101/2024.03.23.586421

**Authors:** Hailey I. Edelstein, Amparo Cosio, Max L. Ezekiel, William K. Corcoran, Aaron H. Morris, Joshua N. Leonard

## Abstract

Synthetic receptors enable bioengineers to build cell-based therapies that perform therapeutic functions in a targeted or conditional fashion to enhance specificity and efficacy. Although many synthetic receptors exist, it remains challenging to generate new receptors that sense soluble cues and relay that detection through orthogonal mechanisms independent of native pathways. Towards this goal, we investigated co-opting natural cytokine receptor ectodomains into Modular Extracellular Sensor Architecture receptors (yielding natural ectodomain, NatE MESA receptors). We generated multiple high-performing, orthogonal synthetic cytokine receptors, identified design principles and constraints, and propose guidance for extending this approach to other natural receptors. We demonstrate utility of NatE MESA by engineering T cells to sense an immunosuppressive cue and respond with customized transcriptional output to support CAR T-cell activity. Finally, we multiplex NatE MESA to logically evaluate multiple cues associated with the tumor microenvironment. These technologies and learnings will enable engineering cellular functions for new applications.

## INTRODUCTION

Engineered cell therapies have emerged as a powerful technology for targeted treatment of complex diseases by leveraging the intrinsic properties of living mammalian cells^1^. These advances are supported by the broadly enabling technology of synthetic receptors, which enable bioengineers to direct engineered cells to sense and respond to specific surface-displayed or soluble environmental cues.^2^ Extension of engineered cell therapies to address unmet therapeutic needs will be driven by, or limited by, the availability of suitable synthetic receptors for addressing application-specific needs.

There exist many classes of synthetic receptors which differ in mechanism and capabilities^2^. One strategy for building such receptors is to generate chimeras in which extracellular domains that sense a target of interest are fused to intracellular domains which redirect ligand binding-induced signaling into natural intracellular signaling pathways of interest. Key examples of this approach include chimeric antigen receptors (CARs)^3–5^, synthetic cytokine receptors (SyCyRs)^6^, and the generalized extracellular molecule sensor (GEMS)^7^. These receptors can be directed to regulate custom transcriptional output using engineered promoters that are responsive to cognate endogenous transcription factors. A potential limitation of this approach, for some applications, is the possibility of crosstalk with native signaling and regulation^2^. An alternative strategy is to design synthetic receptors that signal in a manner that is orthogonal to (i.e., independent of) endogenous signaling. Key examples of this approach include synthetic Notch and synthetic intramembrane proteolysis receptors (synNotch^8,9^, SNIPR^10^) and the modular extracellular sensor architecture (MESA^11,12^). Because orthogonal receptors are self-contained (i.e., they embody the required ligand recognition and signal transduction mechanisms), they are particularly attractive for deployment across different cell types^13^. Orthogonal receptor systems may also be multiplexed to implement information processing, such as conditioning output gene expression on the presence of some cues and absence of others^14,15^. These properties collectively make orthogonal synthetic receptors useful for implementing diverse engineered cellular functions.

Despite collective efforts to engineer synthetic receptors in a modular fashion, targeting new extracellular ligands remains laborious and challenging. First, modification of receptor amino acid sequences may adversely affect surface expression in ways that are difficult to predict^16,17^. Receptor localization can be influenced by post-translational modifications that are protein domain-specific^16^. Design choices that boost surface expression, such as choice of transmembrane domain and linkers, are not always optimal for receptor performance^17^. Second, rationally designing receptor signaling mechanisms that couple ligand binding to subsequent signaling is non-trivial with existing tools, and it still requires extensive protein engineering expertise. Choice of ligand binding domain and the resulting ligand binding complex size and affinity can strongly influence or even preclude signaling via some mechanisms^7,17–19^.

To address these challenges, we investigated a strategy for coopting natural cytokine receptors to build orthogonal, self-contained synthetic receptors. We selected the modular extracellular sensor architecture (MESA), a class of orthogonal, self-contained synthetic receptors that can sense soluble ligands^11,12,20,21^. MESA receptors signal via ligand binding-induced dimerization, which drives intracellular reconstitution of a split protease and *trans*-cleavage to release a transcription factor to drive target gene expression. MESA receptors have incorporated chemically induced dimerization (CID) domains, antibody single chain variable fragments (scFvs), and nanobodies as ligand-binding ectodomains to sense target ligands including small molecules and proteins. We hypothesized that we could employ natural receptor ectodomains (ECDs) as novel MESA ligand-recognition domains. Other synthetic receptors have incorporated natural receptor ligand binding domains using aforementioned chimeric designs, but this approach has generally not been extended to engineer orthogonal synthetic receptors^22–24^. One study implemented a mechanism similar to MESA to employ receptor tyrosine kinase ECDs to achieve autocrine, internal ligand sensing, suggesting that this general approach could be feasible for sensing exogenous ligands^25^. Since there exist multiple options for tuning MESA performance guided by prior knowledge, including split protease reconstitution propensity^21^ and transcription factor properties^26,27^, MESA is well-suited for exploring general strategies for integrating natural receptor ectodomains into synthetic receptors.

In this study, we converted natural human cytokine receptors into orthogonal biosensors by pairing natural receptor ECDs (NatE) with MESA intracellular mechanisms to yield NatE MESA receptors. We investigated four natural receptor candidates and identified ligand-inducible NatE MESA receptor designs for three of the four, although performance varied substantially across receptors. We then synthesized learnings to propose design principles for extending this approach to co-opt other natural receptors. Finally, we demonstrate the utility of our high-performing NatE MESA receptors by employing them to build therapeutically motivated sense-and-respond programs in T cells and logic gates that integrate multiple cues.

## MATERIALS AND METHODS

### General DNA assembly

Plasmid cloning was performed primarily using standard PCR and restriction enzyme cloning with Phusion DNA Polymerase (NEB #M0530L), restriction enzymes (NEB), T4 DNA Ligase (NEB #M0202L), Antarctic Phosphatase (NEB #M0289L) and T4 Polynucleotide Kinase (NEB M0201L). Golden gate assembly and Gibson assembly were also utilized. Plasmids with pcDNA-based backbones (including transcription unit positioning vectors, TUPVs) were transformed into chemically competent TOP10 *Escherichia coli* (Thermo Fisher #C404010), and cells were grown at 37 °C. Plasmids with poly-transfection, transposon, lentiviral, integration vector backbones were transformed into chemically competent NEB Stable *Escherichia coli* (NEB #C3040H), and cells were grown at 30 °C. A complete list of plasmids used in this study can be found in **Supplementary Data 1** and plasmid sequences can be found in **Supplementary Data 2**.

### Source vectors for DNA assembly

Genes encoding each ligand were sourced from: VEGFA165 (pVax1-hVEGF165, which was a gift from Loree Heller, Addgene plasmid #74466)^28^, IL-10 (pHR_Gal4UAS_humanIL-10_T2A_PDL1_PGK_mCherry, which was a gift from Wendell Lim, Addgene plasmid #85430)^9^, TNF (pLI_TNF, which was a gift from Veit Hornung, Addgene plasmid #171179)^29^, and TGF-β1 (TGFB1_pLX307, which was a gift from William Hahn and Sefi Rosenbluh, Addgene plasmid #98377)^30^. DsRed-Express2 was obtained by site directed mutagenesis of pDsRed2-N1, which was a gift from David Schaffer (University of California, Berkeley). The mMoClo BxB1 recombinase expression vector was a gift from Ron Weiss^31^. The red crt operon used in mMoClo destination vectors was sourced from the landing pad destination vector plasmid, which was also a gift from Ron Weiss^31^. The cHS4 insulator was sourced from PhiC31-Neo-ins-5xTetO-pEF-H2B-Citrin-ins, which was a gift from Michael Elowitz (Addgene plasmid #78099)^32^. The CAG promoter was sourced from pR26R CAG/GFP Asc, which was a gift from Ralf Kuehn (Addgene plasmid #74285)^33^. The human EF1α promoter was sourced from pLVX-Tet3G (Clontech/Takara). Barcodes used for the TUPVs were designed by the Elledge lab^34^. BlastR was sourced from lenti dCAS-VP64_Blast, which was a gift from Feng Zhang (Addgene plasmid #61425)^35^. PuroR and HygroR were sourced from pGIPZ (Open Biosystems). Receptor components including TEVp and split TEVp mutants, the murine CD28 TMD, and linkers were sourced from plasmids published in our previous work^17,21^. Genes encoding natural transcription factors were sourced from: cJun (pCLXSN-c-JUN, which was a gift from Jin Chen, Addgene plasmid #102758)^36^ and BATF (pFUW-TetO-BATF, which was a gift from Filipe Pereira, Addgene plasmid #178451)^37^. The NF-κB inducible promoter was sourced from pLNEE^38^. PiggyBac transposon inverted terminal repeats were sourced from pPB_Muc1_mOxGFP_dCT_BlpI which was a gift from Matthew J. Paszek^39^. Sleeping Beauty transposon inverted terminal repeats were sourced from pT4/HB, which was a gift from Wolfgang Uckert, Addgene plasmid #108352)^40^. The gene encoding hyPBase, a hyperactive from of the PiggyBac transposase for high efficiency insertion, was sourced from pCMV-hyPBase, which was a gift from Matthew J. Paszek^41^. The gene encoding SB100X, a hyperactive form of the Sleeping Beauty transposase for high efficiency insertion, was sourced from pCMV(CAT)T7-SB100, which was a gift from Zsuzsanna Izsvak, Addgene plasmid #34879^42^.

### Cloning MESA receptors

In most cases, MESA receptors were first cloned into poly-transfection backbones, which are modified versions of pcDNA to confer high expression in HEK293FT cells and include a secondary transcriptional unit encoding constitutive expression of a fluorescent protein^43^. These plasmid backbones are derived from a modified version of the pcDNA3.1/Hygro(+) Mammalian Expression Vector (Thermo Fisher #V87020), which was modified by our laboratory in previous work (pPD005, Addgene #138749)^26^ and further modified here to change the antibiotic resistance marker from ampicillin to kanamycin and to add the additional expression cassette containing a CMV promoter, fluorescent protein gene, and polyadenylation sequence. Receptors with the CTEVp domain were cloned into a poly-transfection backbone with a constitutively expressed mTagBFP2 gene, while receptors with the NTEVp domain were cloned into a poly-transfection backbone with a constitutively expressed mNeonGreen gene. In general, restriction sites were chosen to facilitate modular swapping of parts via restriction enzyme cloning. Selected receptor constructs were moved into TUPV backbones by restriction enzyme-based cloning to facilitate transposon vector assembly via mammalian modular cloning (mMoClo)^31^. In some cases, receptors were cloned directly into TUPV backbones.

### Cloning ligands for co-expression and secretion

Ligands were cloned into a pcDNA backbone to confer high expression in HEK293FT cells (Addgene #138749)^26^. PCR was used to amplify the coding region of each ligand and append a secretion signal sequence if necessary and restriction enzyme-based cloning was used to insert the products into pcDNA. Selected ligands were moved into a second generation pGIPZ lentiviral vector by restriction enzyme-based cloning to enable generation of stable ligand-secreting HEK293FT and SKOV3 cell lines.

### Cloning reporters

Golden Gate assembly was used to construct all synthetic TF-responsive reporters in a TUPV backbone to facilitate assembly into transposon or landing pad integration vectors^44^. The specific TUPV backbone used for golden gate reporter assembly (pGGB022) includes a pair of BsaI restriction sites upstream of a YB_TATA minimal promoter and a DsRedExpress-2 reporter gene^45^. Promoter inserts containing TF binding sites were synthesized as 15–100 bp oligonucleotides (some promoters were long enough to require multiple inserts) by Integrated DNA Technologies or Life Technologies (Thermo Fisher). The coding and reverse strands were synthesized separately and designed to anneal, resulting in dsDNA with a 4 nt sticky end overhang on each side. The coding and reverse oligonucleotides were mixed (6 µL H_2_O, 1 µL T4 Ligase Buffer, 1 µL T4 PNK (10 U/µL; NEB), 1 µL of each 100 µM oligonucleotide) and phosphorylated at 37 °C for 1 h. They were then denatured at 95 °C for 5 min and cooled slowly to room temperature (here, approximately 22 °C) to allow for annealing. The mix was then diluted 500-fold to make a 20 nM stock and included in the Golden Gate reaction. Golden Gate reaction mixtures comprise 1 µL T4 ligase buffer, 1 µL 10× BSA (1 mg/mL), 0.5 µL BsaI-HF (20 U/µL; NEB), 0.5 µL T4 Ligase (400 U/µL; NEB), 10 fmol of vector, 1 µL of each insert (diluted to 20 nM), and water to a total volume of 10 µL. The reaction was incubated at 37 °C for 1 h, 55 °C for 15 min, and 80 °C for 20 min, and then cooled to room temperature. Then, 3 µL of the reaction was immediately transformed into 50 µL of chemically competent TOP10 *E. coli*.

Natural TF-responsive promoters were constructed using restriction enzyme-based assembly. Promoters were synthesized using the same oligo annealing approach described for synthetic TF-responsive promoters. These inserts were diluted 1:500 to generate a 20 nM stock. The backbone used was a TUPV plasmid containing a different array of binding sites, a YB_TATA minimal promoter, and a DsRedExpress-2 reporter gene. The backbone was cut upstream of the binding site array using BglII (NEB) and in the YB_TATA minimal promoter using SpeI-HF (NEB). The natural TF binding site array inserts were designed to match these overhangs and replace the part of the YB_TATA minimal promoter that was removed when linearizing the backbone. The ligation reactions comprise 2 µL T4 ligase buffer, 1 µL T4 Ligase (400 U/µL; NEB), 1 µL backbone (typically around 50 ng), 3 µL of each insert (diluted to 20 nM), and water to a total volume of 20 µL. The specific binding sites used for each natural TF are as follows: cJun (reporter 1, AP-1 binding site TGAGTCA; reporter 2, cJun dimer binding site TTACCTCA)^46,47^, BATF (BATF/IRF binding site GAAATGAGTCA)^48^, Tbet (palindromic Tbet binding site AATTTCACACCTAGGTGTGAAATT)^49^, NFAT (NFAT binding site ACGCCTTCTGTATGAAACAGTTTTTCCTCC)^50^.

### Cloning transposon vectors and landing pad integration vectors

Both transposon vectors (for genomic integration of receptors and sometimes reporters) and landing pad integration vectors (for genomic integration of reporters) were assembled through a BbsI-mediated Golden Gate reaction based on the previously published mMoClo system^31^. Each 20 μL reaction comprised 2 µL 10× T4 ligase buffer, 2 µL 10× BSA (1 mg/mL stock), 0.8 µL BbsI-HF (NEB), 0.8 µL T4 DNA Ligase (400 U/µL stock), 20 fmol integration vector backbone, and 40 fmol of each transcription unit and linker plasmid to be inserted. The backbone for piggyBac transposon vectors was pHIE426, for sleeping beauty transposon vectors was pHIE430, and for landing pad integration vectors was pPD1178. Transposon vector assemblies typically included transcriptional units encoding receptors, constitutive fluorescent markers, antibiotic selection markers, and sometimes a fluorescent reporter regulated by a synthetic TF-responsive promoter. Landing pad integration vector assemblies typically included transcriptional unit(s) encoding fluorescent reporters regulated by synthetic TF-responsive promoters, constitutive fluorescent markers, and antibiotic selection markers. The Golden Gate reaction was incubated at 37 °C for 15 min, then subjected to 55 iterations of thermocycling (37 °C for 5 min, 16 °C for 3 min, repeat), followed by 37 °C for 15 min, 50 °C for 5 min, 80 °C for 10 min to terminate the reactions; then the mixture was cooled to room temperature and placed on ice prior to immediate transformation into NEB Stable *E. coli*. Because all three backbones used here contain the crt operon, colonies that were transformed with unmodified backbone appeared red after 24 h of growth and were discarded. Additional information on landing pad integration vector design is depicted in **Supplementary Figure 1.**

### Plasmid preparation

In most cases, TOP10 or NEB Stable *E. coli* were grown overnight in 100 mL of LB with the appropriate selective antibiotic. The following morning, cells were pelleted at 3000*g* for 10 min and then resuspended in 4 mL of a solution of 25 mM Tris pH 8.0, 10 mM EDTA, and 15% sucrose. Cells were lysed for 15 min by addition of 8 mL of a solution of 0.2 M NaOH and 1% SDS, followed by neutralization with 5 mL of 3 M sodium acetate (pH 5.2). Precipitate was pelleted by centrifugation at 9000*g* for 20 min. Supernatant was decanted and treated with 3 µL of RNAse A (Thermo Fisher) for 1 h at 37 °C. 5 mL of phenol chloroform was added, and the solution was mixed and then centrifuged at 7500*g* for 20 min. The aqueous layer was removed and subjected to another round of phenol chloroform extraction with 7 mL of phenol chloroform. The aqueous layer was then subjected to an isopropanol precipitation (41% final volume isopropanol, 10 min at room temperature, 9000*g* for 20 min), and the pellet was briefly dried and resuspended in 420 µL of water. The DNA mixture was incubated on ice for at least 12 h in a solution of 6.5% PEG 20,000 and 0.4 M NaCl (1 mL final volume). DNA was precipitated with centrifugation at maximum speed for 20 min.

The pellet was washed once with ethanol, dried for several h at 37 °C, and resuspended for several h in TE buffer (10 mM Tris, 1 mM EDTA, pH 8.0). DNA purity and concentration were confirmed using a Nanodrop 2000 (Thermo Fisher). In some cases, TOP10 or NEB Stable *E. coli* were grown overnight in 50-100 mL of LB with the appropriate selective antibiotic and DNA was prepped using a ZymoPURE II Plasmid Midiprep Kit (Zymo #D4201) by following the manufacturer’s instructions. In a given experiment, all receptor variants being compared were prepped using the same method. For some transposon vectors and reporter landing pad integration vectors, 10 mL of NEB Stable *E. coli* were grown overnight in LB with the appropriate selective antibiotic and DNA was prepped using a ZymoPURE Plasmid Miniprep Kit (Zymo #D4210) by following the manufacturer’s instructions.

### Cell culture

The HEK293FT cell line was purchased from Thermo Fisher/Life Technologies (RRID: CVCL_6911) and was not further authenticated. The HEK293FT-LP cell line was a gift from Ron Weiss and was authenticated by flow cytometric analysis of EYFP expression, which was shown to be homogenous and stable over time—a pattern which is consistent with the original description of this cell line^31^. HEK293FT and HEK293FT-LP cells were cultured in DMEM (Gibco 31600-091) with 10% FBS (Gibco 16140-071), 6 mM L-glutamine (2 mM from Gibco 31600-091 and 4 mM from additional Gibco 25030-081), penicillin (100 U/μL), and streptomycin (100 μg/mL) (Gibco 15140122), in a 37 °C incubator with 5% CO_2_. HEK293FT and HEK293FT-LP cells were subcultured at a 1:5 to 1:10 ratio every 2–3 d using Trypsin-EDTA (Gibco 25300-054). All HEK293FT cell lines generated by engineering either the HEK293FT or HEK293FT-LP parent cell lines were cultured in the same way. The Jurkat T cell line (ATCC TIB-152) was purchased from ATCC and was not further authenticated. Jurkat T cells were cultured in Rosewell Park Memorial Institute Medium (RPMI 1640, Gibco 31800-105) supplemented with 10% FBS (Gibco 16140-071), 4 mM L-glutamine (Gibco 25030-081), penicillin (100 U/μL), and streptomycin (100 μg/mL) (Gibco 15140122). Jurkat T cells were subcultured at a 1:5 or 1:10 ratio every 2-3 d. Cells were maintained at 37 °C with 5% CO_2_. All Jurkat cell lines generated by engineering the parent cell line were cultured in the same way. The HEK293FT, HEK293FT-LP, and Jurkat cell lines tested negative for mycoplasma with the MycoAlert Mycoplasma Detection Kit (Lonza Cat #LT07-318).

### Transient co-transfection of HEK293FTs

Transient co-transfection of HEK293FT cells was conducted using the calcium phosphate method. Cells were plated at a minimum density of 1.0 × 10^5^ cells per well in a 24-well plate in 0.5 ml DMEM, supplemented as described above. For western blot and surface staining experiments, cells were plated at a minimum density of 2.0 × 10^5^ cells per well in a 12-well plate in 1 ml DMEM. After about 24 h, by which time the cells had adhered to the plate and grown to about 50% confluency in the well, the cells were transfected. Plasmids (up to 500 ng DNA for 24-well plates and up to 1,000 ng DNA for 12-well plates) were mixed in H_2_O, and 2 M CaCl_2_ was added to a final concentration of 0.3 M CaCl_2_. This mixture was added dropwise to an equal-volume solution of 2× HEPES-buffered saline (280 mM NaCl, 0.05 M HEPES, 1.5 mM Na_2_HPO_4_) and gently pipetted up and down four times. After 4 min, the solution was mixed vigorously by pipetting ten times. Next, 100 µl of this mixture was added dropwise to the plated cells in 24-well plates (or 200 µl to 12-well plates), and the plates were gently swirled. All receptor plasmid masses were calculated by normalizing to a copy number of 4.22 × 10^9^ for 24-well plates and 8.44×10^9^ for 12-well plates, approximately 35-40 ng per 24-well and 70-80 ng per 12-well. These numbers were determined empirically from prior experiments involving rapamycin-sensing receptors^17^ and the plasmid masses scale with well size. For conditions that received co-expressed ligands, 20 ng of ligand-expressing plasmid was included per 24-well. The total mass of all samples in an experiment was held constant by supplementing with empty vector filler DNA (pHIE298). A complete set of plasmid masses used in this study can be found in **Supplementary Data 3**. The next morning, the medium was aspirated and replaced with fresh medium.

### Transient poly-transfection of HEK293FTs

Transient poly-transfection of HEK293FT cells was conducted using the calcium phosphate method. Cells were plated at a minimum density of 5.05 × 10^5^ cells per well in a 6-well plate in 2 ml DMEM, supplemented as described above. After about 24 h, by which time the cells had adhered to the plate and grown to about 75% confluency in the well, the cells were transfected. Complexes for the two receptor-encoding plasmids being transfected were complexed and added to cells separately. Plasmids (up to 1 µg DNA per mix for 2 mixes per well) were mixed in H_2_O, and 2 M CaCl_2_ was added to a final concentration of 0.3 M CaCl_2_. This mixture was added dropwise to an equal-volume solution of 2× HEPES-buffered saline (280 mM NaCl, 0.05 M HEPES, 1.5 mM Na_2_HPO_4_) and gently pipetted up and down four times. After 4 min, the solutions were mixed vigorously by pipetting ten times. Next, 200 µl of each of each mixture (to make 400 uL total mixture) were added dropwise to the plated cells sequentially, and the plates were gently swirled. All receptor plasmid masses were calculated by normalizing to a copy number of 16.88×10^9^ for 6-well plates, approximately 140-160 ng per 6-well. These numbers were determined empirically from prior experiments involving rapamycin-sensing receptors^17^ and the plasmid masses scale with well size. The total mass of all samples in an experiment was held constant by supplementing with empty vector filler DNA (pHIE298). A complete set of plasmid masses used in this study can be found in **Supplementary Data 3**. The next morning, the medium was aspirated and replaced with fresh medium. An overview of data collection and analysis strategy for poly-transfected samples can be found in **Supplementary Figure 2**.

### Signaling assays with co-expressed ligand

Cells were transfected as described in the section on transient co-transfection of HEK293FTs. The morning after transfection, medium was aspirated and replaced with fresh medium. Typically, at 36–48 h post-transfection and at least 24 h post-media change, cells were harvested. Cells were harvested for flow cytometry by washing with PBS pH 7.4 and using 0.05% Trypsin-EDTA (Thermo Fisher Scientific #25300120) for 5 min followed by quenching with medium. Cell suspensions were pipetted and added to 1 mL of FACS buffer (PBS pH 7.4, 2–5 mM EDTA, 0.1% BSA). Cells were spun at 150×g for 5 min, supernatant was decanted, and fresh FACS buffer was added.

### Signaling assays with exogenous, recombinant ligand

For experiments with transfected receptors, HEK293FT cells were transfected using lipofectamine LTX based on the manufacturer’s recommendations and no co-expressed ligand was included. Briefly, cells were plated in 12-well plates at a density of 2.0 × 10^5^ cells per well in complete DMEM the left to adhere. After 24 h, cells were transfected with 100 µl of mix containing 8.44 × 10^9^ receptor plasmid copies and supplemented with filler DNA (pHIE298) to reach a total mass of 1,000 ng. A complete set of plasmid masses used in this study can be found in **Supplementary Data 3**. The morning after transfection (14 h after transfection), medium was aspirated and replaced with fresh medium with or without recombinant ligand, typically 100 ng/mL VEGF (Biolegend #583706), 100 or 250 ng/mL IL-10 (Biolegend #573206), or 250 ng/mL TNF (ACROBiosystems, ActiveMax #TNA-H4211). Details on ligand stock concentrations can be found in **Supplementary Table 1**. The next day (38 h after transfection), cells were passaged at a 1:2 subculture ratio and treated again with or without ligand. The next day (62 h after transfection), cells were harvested for flow cytometry by washing with PBS pH 7.4 and using 0.05% Trypsin-EDTA (Thermo Fisher Scientific #25300120) for 5 min followed by quenching with medium. Cell suspensions were pipetted and added to 1 mL of FACS buffer (PBS pH 7.4, 2–5 mM EDTA, 0.1% BSA). Cells were spun at 150 × g for 5 min, supernatant was decanted, and fresh FACS buffer was added.

For experiments with stably expressed receptors, HEK293FT cells were plated in 24-well plates at a density of 0.75 × 10^5^ cells per well. The next morning, medium was aspirated and replaced with fresh medium with or without recombinant ligand, typically 100 ng/mL VEGF (Biolegend #583706) or 250 ng/mL IL-10 (Biolegend #573206). Cells were left alone for 48 h before harvesting for flow. Cells were harvested for flow cytometry by washing with PBS pH 7.4 and using 0.05% Trypsin-EDTA (Thermo Fisher Scientific #25300120) for 5 min followed by quenching with medium. Cell suspensions were pipetted and added to 1 mL of FACS buffer (PBS pH 7.4, 2–5 mM EDTA, 0.1% BSA). Cells were spun at 150 × g for 5 min, supernatant was decanted, and fresh FACS buffer was added. For experiments with stably expressed receptors in Jurkats, cells were plated in 0.25 mL/well in 48-well plates or in 0.5 mL in 24-well plates at a density of 2 × 10^5^ cells/mL. At the time of plating, recombinant ligand was added to reach the desired final concentration, typically 100 or 250 ng/mL IL-10 (Biolegend #573206). Cells were left alone for 48 h before harvesting for flow or passaging at a split ratio of 1:5 and re-dosed with fresh ligand for another 48 h before harvesting. Cells were harvested for flow cytometry by pipetting up and down gently and adding to 1 mL of FACS buffer (PBS pH 7.4, 2–5 mM EDTA, 0.1% BSA). Cells were spun at 125 × g for 5 min, supernatant was decanted, and fresh FACS buffer was added.

### Time course microscopy signaling assays with exogenous ligand

For time course microscopy experiments, cells were plated at a density of 0.25 × 10^5^ cells per well in 48-well plates. The next morning, IL-10 was diluted in serum free (incomplete) DMEM to a concentration of 25 ng/µL and 5 µL of this mix was added to the side of each well to make the final concentration 250 ng/µL. For untreated wells, 5 µL of serum free DMEM was added. Plates were immediately placed in the microscope (Keyence BZ-X800E), well positions were chosen using the brightfield channel, and the time course was started. Plates were held in a stage top incubator (TOKAI HIT, INU-KIW-F1) to maintain humidity, a temperature of 37 °C, and CO_2_ concentration of 5%. Images were taken every 2 hrs. for 48 hrs. A BZ-X GFP filter (Ex 470/40 nm, Em 525/50 nm, dichroic 495 nm) was used to measure mNeonGreen fluorescence and a BZ-X Texas Red filter (Ex 560/40 nm, Em 630/75 nm, dichroic 585 nm) was used to measure DsRed-Express2 reporter fluorescence. The following exposure times were used: 1/5 sec for mNeonGreen, 1/40 sec for DsRed-Express2, 1/7500 sec for brightfield using 25% transmitted light with oblique illumination. Images were acquired using BZ Series Application software v01.01.00.17 and PlanApo 4× objective with a numerical aperture of 0.2. Images were not modified or corrected in any way. Images were processed using custom software in MATLAB to extract the fluorescence in the green channel per pixel, discard non-green pixels, and extract the fluorescence in the red channel for green pixels only. We then calculate the average dsRedExpress2 fluorescence across the selected pixels for each field of view at each timepoint.

### CAR-mediated NFAT signaling assays

To assess CAR expression and activity in Jurkat cells, Jurkat cell lines modified with various CAR-expressing circuits were co-cultured with target cancer cells and/or IL-10. Target cells used here were SKOV3 cells, which express the CAR T1E28z target antigen HER2^51^. Either unmodified SKOV3 cells or IL-10 secreting SKOV3 cells (cells transduced with lentiviral vector pHIE730 encoding constitutive secretion of human IL-10 and puromycin resistance and selected with 1 ug/mL puromycin) were plated at a density of 3 × 10^4^ cells per well of a 48-well plate in 300 μL of complete DMEM and left to adhere overnight. About 20 h later, 100 μL media was removed from each well to leave about 150 μL of remaining media per well. 150 μL of complete RPMI containing Jurkat cell lines at a density of 2 × 10^5^ cells/mL was added such that the final Jurkat cell concentration in each well was 1 × 10^5^ cells/mL. The Jurkat cell suspensions also contained either 500 ng/mL IL-10 (such that the final concentration per well was 250 ng/mL IL-10) or an equivalent volume of vehicle, water. Conditions without SKOV3 cells were prepared my adding 150 μL complete DMEM to empty 48-wells and then supplementing with the same Jurkat +/- IL-10 suspensions added to SKOV3-containing wells. The cells were incubated at 37 °C for approximately 48h before harvesting for flow. To harvest Jurkat cells and minimize harvested SKOV3 cells, the full volume of media in each well was pipetted gently 4-5 times and added to 1 mL of FACS buffer containing 3 uM DAPI (PBS pH 7.4, 2–5 mM EDTA, 0.1% BSA, 3 uM DAPI (Thermo Scientific #62247)). Cells were spun at 125 × g for 5 min, supernatant was decanted, and fresh FACS buffer was added before flow analysis. Generally, the larger size of SKOV3 cells compared to Jurkats enabled distinction of the two cell types by FSC vs. SSC. For engineered Jurkat cell lines that express a constitutive mNeonGreen fluorescent protein, only mNeonGreen+ engineered Jurkat cells were evaluated, adding an additional gating layer to exclude any SKOV3 cells harvested.

### Immunohistochemistry (surface staining)

For surface staining to quantify MESA receptor surface expression, HEK293FT cells were plated at 2 × 10^5^ cells per well in 1 mL DMEM in 12-well plates 24 h before transfection and transfected as described above, using 200 μL transfection reagent per well. The receptor plasmid copy number for individual receptors matches the copy number used in functional assays, scaled up to 12-well format (8.44 × 10^9^ receptor plasmid copies). At 36–48 h after transfection, cells were harvested with 500 µl FACS buffer and spun at 150 × g at 4 °C for 5 min. Supernatant was decanted, and 50 µL fresh FACS buffer and 10 µL human IgG (Human IgG Isotype Control, ThermoFisher Scientific #02-7102, RRID: AB_2532958, stock concentration 1 mg/mL) was added. Cells were incubated in this mixture at 4 °C for 5 min. Next, 5 µL FLAG tag antibody (Anti-DDDDK-APC, Abcam ab72569, RRID: AB_1310127) was added at a concentration of 0.5 µg per sample and cells incubated at 4 °C for 30 min. Following incubation, 1 mL of FACS buffer was added, cells were spun at 150 × g at 4 °C for 5 min, and supernatant was decanted. This wash step was repeated two more times to total three washes. After decanting supernatant in the final wash, 1–3 drops of FACS buffer were added.

For surface staining to quantify CAR expression, HEK293FTs that were genomically engineered to contain the circuit components for CAR expression were plated at 2 × 10^5^ cells per well in 1 mL DMEM and treated with or without 250 ng/mL recombinant IL-10 for 48 h before harvesting. Cells were harvested as described for MESA receptor surface staining. For HEK293FTs transfected with circuit components for CAR expression, cell plating and harvest is identical to that described for MESA receptor surface staining. For Jurkats that were genomically engineered to contain the circuit components for CAR expression, cells were plated at 2 × 10^5^ cells/mL in 1 mL in 12-well plates with or without 250 ng/mL recombinant IL-10 and incubated for 48 h. Jurkats were harvested by collecting the full volume remaining in each well, diluting in 1 mL FACS buffer, and spinning down at 150 × g at 4 °C for 5 min. Supernatant was decanted, and 50 µL fresh FACS buffer and 10 µL human IgG (Human IgG Isotype Control, ThermoFisher Scientific #02-7102, RRID: AB_2532958, stock concentration 1 mg/mL) was added. Cells were incubated in this mixture at 4 °C for 5 min. Next, 2.5 µL anti-hEGF tag antibody (Anti-hEGF-biotin, R&D Systems BAF236) was added at a concentration of 0.5 µg per sample and cells incubated at 4 °C for 20 min. Following incubation, 1 mL of FACS buffer was added, cells were spun at 150 × g at 4 °C for 5 min, and supernatant was decanted. Next, 2.5 µL Strep-APC (Abcam ab134362) was added at a concentration of 0.5 µg per sample and cells incubated at 4 °C for 20 min. Following incubation, 1 mL of FACS buffer was added, cells were spun at 150 × g at 4 °C for 5 min, and supernatant was decanted. This wash step was repeated two more times to total three washes. After decanting supernatant in the final wash, 1–3 drops of FACS buffer were added. For Jurkats, FACS buffer contained 3 µM DAPI for viability staining.

### Analytical flow cytometry

Flow cytometry was run on a BD LSR Fortessa Special Order Research Product (Robert H. Lurie Cancer Center Flow Cytometry Core). Lasers and filter sets used for data acquisition are listed in **Supplementary Table 2**. Approximately 3,000–10,000 single, transfected cells were analyzed per sample in transfection experiments. Transfected cells were identified using a separate, single transfection control fluorescent protein or multiple fluorescent proteins encoded on receptor plasmids (e.g., mNeonGreen and mTagBFP2). In cases where the transfected cells are landing pad engineered cells, a constitutive miRFP720 gene is included in landing pad cargo so cells are first identified by miRFP720 expression before setting transfection gate(s). In cases where the cells were engineered with transposons, a constitutive mNeonGreen gene is included in the transposon so engineered cells were identified by mNeonGreen fluorescence.

Samples were analyzed using FlowJo v10 software (FlowJo, LLC). Fluorescence data were compensated for spectral bleed-through. As shown in **Supplementary Figure 3a**, the HEK293FT cell population was identified by SSC-A versus FSC-A gating, and singlets were identified by FSC-A versus FSC-H gating. To distinguish transfected from non-transfected cells, a control sample of cells was generated by transfecting cells with a mass of pcDNA (empty vector) equivalent to the mass of DNA used in other samples in the experiment. For the single-cell subpopulation of the pcDNA-only sample, a gate was made to identify cells that were positive for the constitutive fluorescent protein(s) used as a transfection control in other samples, such that the gate included no more than 1% of the non-fluorescent cells (**Supplementary Figure 3b**). A similar approach was used to identify genomically modified miRFP720 or mNeonGreen expressing HEK293FTs. The Jurkat cell population was identified by SSC-A versus FSC-A gating, and singlets were identified by FSC-A versus FSC-H gating. Live cells were identified by inclusion of a DAPI viability stain such that DAPI+ (dead) cells were excluded from analysis. Genomically engineered Jurkats were identified by mNeonGreen expression by drawing a gate on un-modified Jurkats such that the gate included no more than 1% of the non-fluorescent cells.

### Generation of stable receptor cell lines

To generate HEK293FT cell lines, from exponentially growing cells, 5 × 10^4^ cells were plated per well (0.5 mL medium) in 24-well format, and cells were cultured for 24 h to allow cells to attach and spread. HyPBase (hyperactive PiggyBac transposase, pHIE425) or SB100X (hyperactive Sleeping Beauty transposase, pHIE429) encoded on pcDNA-based plasmids were co-transfected with their respective transposon vector by lipofection with Lipofectamine LTX with PLUS Reagent (ThermoFisher 15338100). 100 ng of transposase expression vector was mixed with 500 ng of reporter-containing integration vector, 0.6 μL of PLUS reagent, and enough OptiMEM (ThermoFisher/Gibco 31985062) to bring the mix volume up to 25 μL. In a separate tube, 2 μL of LTX reagent was mixed with 23 μL of OptiMEM. The DNA/PLUS reagent mix was added to the LTX mix, pipetted up and down four times, and then incubated at room temperature for 5 min. 50 μL of this transfection mix was added dropwise to each well of cells and mixed by gentle swirling. Cells were cultured until the well was ready to split (typically 3 d), without any media changes.

To begin selection of HEK293FT cells that successfully integrated the transposon vector, cells were harvested from the 24-well plate when confluent by trypsinizing and transferring to a single well of a 12-well plate in 1 mL of medium supplemented with 1 μg/mL puromycin (Invivogen ant-pr) or 200 μg/mL hygromycin (Millipore, #400053) depending on the vector. Cells were trypsinized daily (typically 3 d) until cell death was no longer evident. Cells were cultured in medium supplemented with puromycin through exponential expansion until reaching a confluent 10 cm dish, upon which cells were frozen. Selective pressure was maintained when culturing these cells but not included during experiments.

For HEK293FT sorting experiments, cells were harvested by trypsinizing, resuspended at approximately 10^7^ cells per mL in pre-sort medium (DMEM with 10% FBS, 25 mM HEPES (Sigma H3375), and 100 µg/mL gentamycin (Amresco 0304)), and held on ice until sorting was performed. Cells were sorted using a BD FACS Aria 4-laser Special Order Research Product (Robert H. Lurie Cancer Center Flow Cytometry Core). Details on laser and channel configurations can be found in **Supplementary Table 3.** The first sorting strategy was as follows: single cells were first gated to exclude all mNeonGreen negative cells (as mNeonGreen is a constitutive marker in the transposon and successfully engineered, non-silenced cells should express this protein). Then, the mNeonGreen positive population was broken into octiles that each contained 12.5% of the mNeonGreen positive population and the top (brightest mNeonGreen) four octiles were sorted. The second sorting strategy required an additional gate for exclusion of reporter (DsRed-Express2) positive cells to isolate low background cells and the same octile sorting strategy was applied to this mNeonGreen+/DsRed-Express2-population. The top four mNeonGreen octiles were sorted. 50,000 cells were collected in post-sort medium (DMEM with 20% FBS, 25 mM HEPES, and 100 μg/mL gentamycin), and cells were held on ice until they could be centrifuged at 150 × g for 5 min, resuspended in 0.5 mL complete medium supplemented with 100 µg/mL gentamycin, and plated in one well of a 24-well plate. Cells were maintained in gentamycin for 7 d after sorting during expansion before banking. Cells were thawed for use in experiments in this study.

To generate Jurkat cell lines, from exponentially growing cells, 5 × 10^4^ cells were plated per well (0.5 mL medium) in 24-well format at the time of transfection. HyPBase (hyperactive PiggyBac transposase, pHIE425) was co-transfected with a transposon vector by lipofection with Lipofectamine LTX with PLUS Reagent (ThermoFisher 15338100). 100 ng of transposase expression vector was mixed with 500 ng of reporter-containing integration vector, 0.6 μL of PLUS reagent, and enough OptiMEM (ThermoFisher/Gibco 31985062) to bring the mix volume up to 25 μL. In a separate tube, 2 μL of LTX reagent was mixed with 23 μL of OptiMEM. The DNA/PLUS reagent mix was added to the LTX mix, pipetted up and down four times, and then incubated at room temperature for 30-60 min. 50 μL of this transfection mix was added dropwise to each well of cells and mixed by gentle swirling. Cells were cultured until the well was ready to split (typically 3 d), without any media changes.

To begin selection of Jurkat cells that successfully integrated the transposon vector, cells were harvested from the 24-well plate and transferred to a single well of a 12-well plate in 1 mL of fresh medium. After 24 h, the entire contents of the 12-well was transferred to a 6-well and supplemented with 0.2 μg/mL puromycin (Invivogen ant-pr). Cells were analyzed daily and spun at 125 × g for 5 min at 4 °C approximately once per week to fully refresh media. Every 3 days, 1 mL puro-containing medium was added. Once cells were exponentially growing, they were transferred up to 10 cm dishes in medium supplemented with puromycin, passaged once without puromycin, and frozen. Selective pressure was maintained when culturing these cells but not included during experiments.

To sort, Jurkat cells were harvested, spun down at 125 × g for 5 min at 4 °C before resuspending at approximately 10^7^ cells/mL in pre-sort medium (RPMI with 10% FBS, 25 mM HEPES (Sigma H3375), and 100 µg/mL gentamycin (Amresco 0304)), and held on ice until sorting was performed. Cells were sorted using a BD FACS Aria 4-laser Special Order Research Product (Robert H. Lurie Cancer Center Flow Cytometry Core). Details on laser and channel configurations can be found in **Supplementary Table 3.** The sorting strategy was as follows: single cells were first gated to exclude all mNeonGreen negative cells (as mNeonGreen is a constitutive marker in the transposon and successfully engineered, non-silenced cells should express this protein). Then, the mNeonGreen positive population was broken into octiles that each contained 12.5% of the mNeonGreen positive population and the top (brightest mNeonGreen) octile was sorted. 50,000 cells were collected in post-sort medium (DMEM with 20% FBS, 25 mM HEPES, and 100 μg/mL gentamycin), and cells were held on ice until they could be centrifuged at 125 × g for 5 min at 4 °C, resuspended in 0.5 mL complete medium supplemented with 100 µg/mL gentamycin, and plated in one well of a 24-well plate. Cells were maintained in gentamycin for 7 d after sorting during expansion before banking. Cells were thawed for use in experiments in this study.

### Generation of stable reporter cell lines

From exponentially growing HEK293LP cells, which were a gift from Ron Weiss, 5 × 10^4^ cells were plated per well (0.5 mL medium) in 24-well format, and cells were cultured for 24 h to allow cells to attach and spread^31^. More information on integration vector design can be found in **Supplementary Figure 1a**. Bxb1 recombinase was co-transfected with the integration vector by lipofection with Lipofectamine LTX with PLUS Reagent (ThermoFisher 15338100). 300 ng of BxB1 expression vector was mixed with 300 ng of reporter-containing integration vector, 0.5 μL of PLUS reagent, and enough OptiMEM (ThermoFisher/Gibco 31985062) to bring the mix volume up to 25 μL. In a separate tube, 1.9 μL of LTX reagent was mixed with 23.1 μL of OptiMEM. The DNA/PLUS reagent mix was added to the LTX mix, pipetted up and down four times, and then incubated at room temperature for 5 min. 50 μL of this transfection mix was added dropwise to each well of cells and mixed by gentle swirling. Cells were cultured until the well was ready to split (typically 3 d), without any media changes.

To begin selection of cells that successfully integrated the reporter integration vector, cells were harvested from the 24-well plate when confluent by trypsinizing and transferring to a single well of a 6-well plate in 2 mL of medium supplemented with 1 μg/mL puromycin (Invivogen ant-pr). Cells were trypsinized daily (typically 3 d) until cell death was no longer evident. Cells were cultured in medium supplemented with puromycin until the 6-well was confluent and cells were exponentially growing. Cells were then selected with 6 μg/mL blasticidin (Alfa Aesar/ThermoFisher J61883) for 7 d. Cells were cultured in both puromycin and blasticidin to maintain selective pressure until flow sorting (**Supplementary Figure 1b**).

To sort, cells were harvested by trypsinizing, resuspended at approximately 10^7^ cells per mL in pre-sort medium (DMEM with 10% FBS, 25 mM HEPES (Sigma H3375), and 100 µg/mL gentamycin (Amresco 0304)), and held on ice until sorting was performed. Cells were sorted using a BD FACS Aria 4-laser Special Order Research Product (Robert H. Lurie Cancer Center Flow Cytometry Core). The sorting strategy was as follows: single cells were first gated to exclude all EYFP positive cells (as EYFP positive cells still have an intact landing pad locus, suggesting a mis-integration event occurred) and to include only miRFP720+ cells. EYFP expression was measured using the FITC channel and miRFP720 expression was measured using a modified APC-Cy7 channel (details on channel configurations can be found in **Supplementary Table 3**). Then a gate was drawn on miRFP720 expression to capture the 88th to 98th percentile of miRFP720-expressing cells (the top 2% were excluded to exclude cells suspected to possess two or more integrated copies of the cargo vector). 50,000 cells were collected in post-sort medium (DMEM with 20% FBS, 25 mM HEPES, and 100 μg/mL gentamycin), and cells were held on ice until they could be centrifuged at 150 × g for 5 min, resuspended in 0.5 mL complete medium supplemented with 100 µg/mL gentamycin, and plated in one well of a 24-well plate. Cells were maintained in gentamycin for 7 d after sorting during expansion before banking. Cells were thawed for use in experiments in this study. Additional information is depicted in **Supplementary Figure 1c**.

### Quantification of reporter output

MESA signaling was quantified by measuring the expression of a downstream fluorescent reporter protein, DsRed-Express2, regulated by a synTF-inducible promoter. The mean fluorescence intensity (MFI) for each relevant channel (as defined in **Supplementary Table 2**) of the single cell transfected population (transfection control marker+) or the single cell transposon modified population (mNeonGreen+) was calculated and exported for further analysis. To calculate reporter expression, MFI in the PE-Texas Red channel was averaged across three biological replicates. The MFI was converted to Molecules of Equivalent PE-Texas Red (MEPTRs); as shown in **Supplementary Figure 3c**, to determine conversion factors for MFI to MEPTRs, UltraRainbow Calibration Particles (Spherotech #URCP-100-2H) were run with each flow cytometry experiment. These reagents contain nine subpopulations of beads, each with a known number of various fluorophores. The total bead population was identified by FSC-A vs. SSC-A gating, and bead subpopulations were identified through two fluorescent channels. MEPTR values corresponding to each subpopulation were supplied by the manufacturer. A calibration curve was generated for the experimentally determined MFI vs. the manufacturer supplied MEPTRs, and a linear regression was performed with the constraint that 0 MFI equals 0 MEPTRs. The slope from the regression was used as the conversion factor, and error was propagated. Fold differences were calculated by dividing reporter expression with ligand treatment by the reporter expression without ligand treatment. Standard error was propagated through all calculations.

### Western blotting

For western blotting to detect MESA receptor expression, HEK293FT cells were plated at 2 × 10^5^ cells per well in 1 mL DMEM in 12-well plates 24 h before transfection and transfected as above, using 200 μL transfection reagent per well (the reaction scales with the volume of medium). At 36–48 h after transfection, cells were lysed with 250 μL RIPA (150 mM NaCl, 50 mM Tris-HCl pH 8.0, 1% Triton X-100, 0.5% sodium deoxycholate and 0.1% sodium dodecyl sulfate) with protease inhibitor cocktail (Pierce/Thermo Fisher #A32953) and incubated on ice for 30 min. Lysate was cleared by centrifugation at 14 000 × g for 20 min at 4 °C, and supernatant was harvested. A BCA assay was performed to determine protein concentration, and after a 10 min incubation in Laemmli buffer (final concentration 60 mM Tris-HCl pH 6.8, 10% glycerol, 2% sodium dodecyl sulfate, 100 mM dithiothreitol and 0.01% bromophenol blue) at 70 °C, protein (10– 25 μg, the maximum amount of protein that could be loaded to keep protein mass constant across all samples on a gel) was loaded onto a 4–15% Mini-PROTEAN TGX Precast Protein Gel (Bio-Rad) and run at 50 V for 10 min followed by 100 V for at least 1 h. Wet transfer was performed onto an Immuno-Blot PVDF membrane (Bio-Rad) for 45 min at 100 V. Ponceau-S staining was used to confirm protein transfer.

For detection of MESA receptors using the N-terminal 3xFLAG epitope tag, membranes were blocked for 30 min with 3% milk in Tris-buffered saline pH 8.0 (TBS pH 8.0: 50 mM Tris, 138 mM NaCl, 2.7 mM KCl, HCl to pH 8.0), washed once with TBS pH 8.0 for 5 min and incubated for overnight at 4 °C in primary antibody (Mouse-anti-FLAG M2 [Sigma #F1804, RRID: AB_262044] [http://antibodyregistry.org/AB_262044]), diluted 1:1000 in 3% milk in TBS pH 8.0. Primary antibody solution was decanted, and the membrane was washed once with TBS pH 8.0 and twice with TBS pH 8.0 with 0.05% Tween, for 5 min each. Secondary antibody (HRP-anti-Mouse [CST 7076, RRID: AB_330924] [http://antibodyregistry.org/AB_330924]), diluted 1:3000 in 5% milk in TBST pH 7.6 (TBST pH 7.6: 50 mM Tris, 150 mM NaCl, HCl to pH 7.6, 0.1% Tween), was applied for 1 h at room temperature, and the membrane was washed three times for 5 min each time with TBST pH 7.6. After primary and secondary staining and washing, the membrane was incubated with Clarity Western ECL Substrate (Bio-Rad) for 5 min, and then imaged using an Azure c280 (Azure Biosystems). Images were cropped with Illustrator CC (Adobe). No other image processing was employed.

For detection of membrane-bound IL-15 (mbIL-15) using the C-terminal myc epitope tag, membranes were blocked for 30 min with 5% milk in Tris-buffered saline pH 7.6 with 0.1% Tween (TBST pH 7.6: 50 mM Tris, 150 mM NaCl, HCl to pH 8.0, 0.1% Tween-20), washed once with TBST pH 7.6 for 5 min and incubated overnight at 4 °C in primary antibody (Mouse-anti-myc [Abcam #ab32, RRID: AB_303599] [https://antibodyregistry.org/AB_303599]) diluted 1:1000 in 5% milk in TBST pH 7.6. Primary antibody solution was decanted, and the membrane was washed three times with TBST pH 7.6 for 5 min each. Secondary antibody (HRP-anti-Mouse [CST 7076, RRID: AB_330924] [http://antibodyregistry.org/AB_330924]), diluted 1:3000 in 5% milk in TBST pH 7.6, was applied for 1 h at room temperature, and the membrane was washed three times for 5 min each time with TBST pH 7.6. After primary and secondary staining and washing, the membrane was incubated with Clarity Western ECL Substrate (Bio-Rad) for 5 min, and then imaged using an Azure c280 (Azure Biosystems). Images were cropped with Illustrator CC (Adobe). No other image processing was employed.

### Statistical analyses

ANOVA tests and Tukey’s HSD tests were performed using RStudio. Tukey’s HSD tests were performed with *α* = 0.05. Pairwise comparisons were made using a two-tailed Welch’s *t*-test, which is a version of Student’s *t*-test in which the variance between samples is treated as not necessarily equal. Two-tailed Welch’s *t*-tests were performed in GraphPad and Excel. To decrease the false discovery rate, the Benjamini-Hochberg procedure was applied to each set of tests per figure panel; in all tests, after the Benjamini-Hochberg procedure, the null hypothesis was rejected for *p*-values < 0.05. The outcomes for each statistical test are provided in the figure captions and in more detail in **Supplementary Notes 1-3**.

## RESULTS

### Development of synthetic biosensors using natural human cytokine receptors

To investigate strategies for engineering natural ectodomain (NatE) MESA receptors, we defined a limited design space focused on evaluating the feasibility of natural receptor conversion. We first selected a panel of native human cytokine receptors that represent a variety of receptor classes and signaling mechanisms and that—if successful—would yield biosensors useful for detecting soluble factors of relevance to immune-related processes. These receptors included human VEGFR1 and VEGFR2 (for sensing vascular endothelial growth factor, VEGF)^52,53^, IL-10Rα and IL-10Rβ (for sensing interleukin 10, IL-10)^54–57^, TNFR1 and TNFR2 (for sensing tumor necrosis factor, TNF)^58,59^, and TGF-βR1 and TGF-βR2 (for sensing transforming growth factor β, TGF-β)^60,61^ (UniProt entries listed in **Supplementary Table 4**). VEGF, IL-10, and TGF-β are upregulated in the tumor microenvironment, and TNF and IL-10 play immunoregulatory roles, generally driving inflammatory and immunosuppressive states, respectively. Additional background for each of these receptors is provided in subsequent sections and in **Supplementary Notes 4 and 5** (for TNFR and TGF-βR, respectively). These receptors include hetero-associative (different receptor chains associate) and homo-associative (like receptor chains associate) ligand binding-induced signaling mechanisms involving two or more receptor chains, making them promising candidates for coupling to MESA receptors’ split protease reconstitution and *trans*-cleavage signaling mechanism (hypothesized mechanisms for each receptor are summarized in **Supplementary Figure 4**). We next chose a previously validated set of MESA intracellular signaling domains, including the split tobacco etch virus protease (TEVp) mutant 75S/190K and a minimal zinc finger domain-based synthetic transcription factor (synTF)^17,21,26^. We included designs that retain the native transmembrane (TMD) and juxtamembrane (JMD) domains for each selected natural receptor because those domains influence surface expression, ligand-independent receptor chain association, and ligand-induced signaling events^62^. We also evaluated designs that replace the native TMD and JMD with domains validated in prior MESA receptors, including a truncated murine cluster of differentiation 28 (CD28) TMD and a glycine-serine-rich intracellular linker^21^. Finally, we evaluated variants with either the native signal sequence for each selected human receptor or alternative human signal sequences which we hypothesized could improve trafficking to the cell surface if necessary. These include two signal sequences widely used on chimeric antigen receptors (CARs) and synNotch receptors which are derived from the human cluster of differentiation 8a (CD8a) T cell receptor and the human immunoglobulin variable heavy chain (IgG VH)^8,10,63–65^. The exploration of design space described here (and summarized in **Supplementary Figure 4**) provides a starting point for evaluating general prospects for natural receptor conversion.

### Conversion of VEGFR

We first investigated the creation of biosensors for detecting VEGF. VEGF plays a key role in neovascularization, and its dysregulated activity is related to solid tumor growth and development^66,67^. VEGF is upregulated in many tumors and has been harnessed as a tumor microenvironment (TME) marker and therapeutic target^68^. VEGFR1 and VEGFR2 belong to the receptor tyrosine kinase (RTK) family and are the main receptors involved in VEGF signaling and angiogenesis^52,53^. These receptors can homo- or heterodimerize^69,70^ in the absence of VEGF, but they do not signal without VEGF^71^. All VEGF isoforms exist mainly as homodimers^72^. Ligand binding to the receptor’s extracellular domain induces a conformational change, which leads to the activation and autophosphorylation of the intracellular kinase domains, activating downstream signaling cascades (**Figure 1a**)^73^. Given previous studies identifying MESA receptor signaling requirements^17^ and demonstrating the feasibility of employing VEGFR domains in a similar receptor mechanism^25^, we hypothesized that this ligand-induced conformational change could be compatible with the requirements for inducing MESA intracellular signaling.

**Figure 1.**
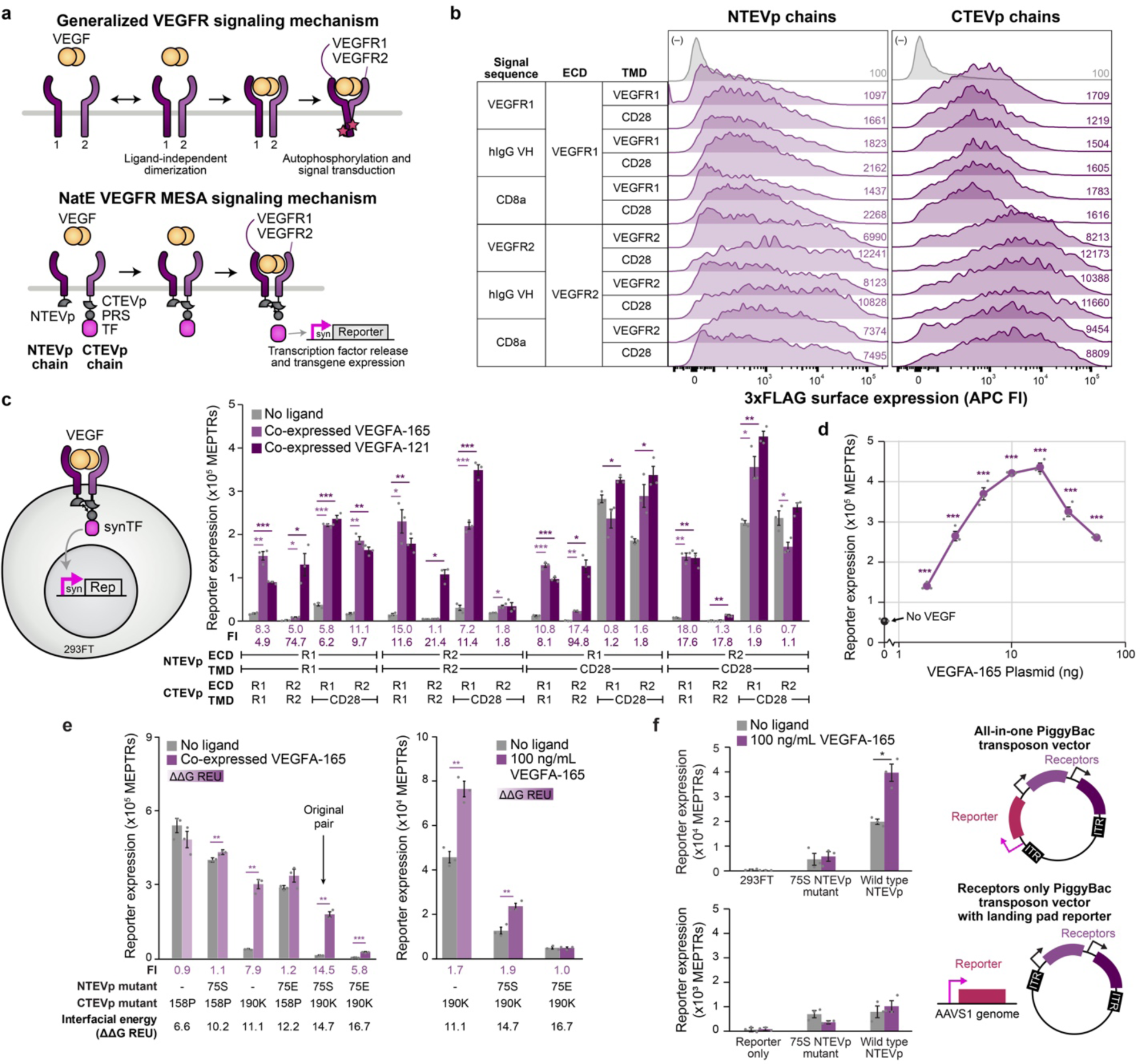
Conversion of VEGFR into a VEGF NatE MESA receptor. **(a)** Schematic of generalized VEGFR signaling mechanism highlighting receptor interactions (top), and schematic of the proposed VEGF NatE MESA signaling mechanism (bottom). **(b)** Surface expression of each chain (transfected HEK293FTs, anti-3xFLAG stain). Histograms show data for transfected (fluorescent) cells and the gray histograms in each column are transfection controls (no receptor). All receptors express well on the cell surface, with VEGFR2 constructs showing higher expression than VEGFR1 constructs. Mean fluorescence intensities are listed. **(c)** Functional evaluation of VEGF NatE MESA receptors (transfected HEK293FTs) in response to autocrine human VEGF (two isoforms). Many receptor configurations exhibit ligand-inducible signaling, with fold induction (FI: induced signal divided by background signal) shown below the plot (two-tailed Welch’s t-test results indicated above each bar pairing for *p* < 0.05). Some receptors signal more in response to one VEGF isoform over another. All receptors include the native signal sequence that matches the ECD. **(d)** Modulating VEGFA-165 expression by varying co-transfected plasmid masses yields dose-dependent receptor signaling. All plasmid masses used confer ligand-induced signaling compared to the no-ligand case (single-factor ANOVA, *p* < 0.001). **(e)** Functional evaluation of VEGF NatE MESA receptors incorporating different split TEVp mutants in response to co-expressed (left) and recombinant (right) human VEGF. Only receptors including the 190K CTEVp mutant show inducible signaling with both ligand formats, with choice of NTEVp mutant modulating overall signal magnitude (two-tailed Welch’s t-test, ** *p* < 0.01, *** *p* < 0.001). **(f)** Functional evaluation VEGF NatE MESA receptors expressed stably from the genome. HEK293FTs were modified with PiggyBac transposons encoding receptors and a reporter in the transposon (top) or encoding just receptors and introduced into a HEK293FT cell line containing a reporter integrated in the AAVS1 locus (bottom). Two receptor configurations were investigated: A CTEVp (190K) chain with a VEGFR1 signal sequence, ECD, and TMD paired with a NTEVp (75S or WT) chain with a VEGFR2 signal sequence, ECD, and TMD. Both transposons constitutively express mNeonGreen and puromycin resistance to facilitate selection and fluorescence-based identification of engineered cells (not shown here but included in schematics in **Supplementary Figure 6h**). Data shown include all mNeonGreen+ cells. Results for two-tailed Welch’s t-test shown for *p* < 0.05. For all bar graphs, bars represent the mean of three biological replicates and error bars depict standard error of the mean (S.E.M.). Abbreviations: VEGFR, vascular endothelial growth factor receptor; PRS, protease recognition sequence; TF, transcription factor; ECD, ectodomain; TMD, transmembrane domain; TEVp, Tobacco Etch Virus protease; NTEVp, N-terminal component of split, mutant TEVp; CTEVp, C-terminal component of split, mutant TEVp; APC FI, allophycocyanin fluorescence intensity; MEPTRs, molecules of equivalent PE-TexasRed; Rep, reporter; FI, fold induction; R1, VEGFR1; R2, VEGFR2; REU, Rosetta energy units (dimensionless).

To investigate conversion of VEGF receptors to NatE MESA receptors, we first evaluated the key property of surface expression. We evaluated VEGFR-based receptor chains with a range of signal sequence, ECD, TMD, and signaling domain choices (**Supplementary Figure 5**). In general, all of the receptor variants tested expressed well on the cell surface (**Figure 1b**). To verify that full-length receptors were expressed, we evaluated whole cell expression by western blot, and all receptors were expressed at their expected size considering post-translational modifications (**Supplementary Figure 6a**). VEGFR2-based receptors exhibit higher surface and whole-cell expression levels than did VEGFR1-based receptors. Signal sequence and TMD choice did not meaningfully affect receptor trafficking to the cell surface (**Figure 1b**).

These observations suggest that ECD choice is the primary determinant of receptor surface expression. Since no one signal sequence was better than the rest (and signal sequence choice would not impact signaling because it is cleaved in the receptor trafficking process), only receptors with natural signal sequences were carried forward.

We next evaluated whether candidate VEGF NatE MESA receptors were functional. First, to quickly identify receptor configurations that are incapable of signaling, we employed an approach in which the ligand is co-expressed (in an autocrine fashion) with the receptor chains (the fluorescent reporter is genomically integrated) (**Figure 1c**). This strategy maximizes the chance for receptor-ligand contacts both in the secretory pathway inside the cell and at the cell surface; if a receptor did not signal in this assay, we deemed it non-functional. We elected to test both the VEGFA165 isoform, which is most abundant and can be bound to the extracellular matrix, and the VEGFA121 isoform, which is only found in soluble form^72^. We observed multiple receptors that exhibited ligand-inducible signaling, including both homo- and heterodimeric receptor configurations (**Figure 1c**, **Supplementary Figure 6b**). Receptors in which both chains contain a CD28 TMD produced high ligand-independent signal (i.e., background), resulting in low fold inductions. This is consistent with previous observations with MESA receptors and CARs, as the CD28 TMD has a high propensity to aggregate^17,74–76^. Receptors in which the CTEVp chain (which includes the C-terminal fragment of TEV protease) contains a VEGFR2 ECD and TMD exhibited signaling only when co-expressed with VEGFA121. We speculate that this could be attributable to differences in binding geometry across the various ECDs and VEGFA variants, which could affect the interaction of the intracellular receptor domains.

We also verified our assumption that changes in signal sequence would not affect receptor signaling competency, although some variations in reporter output magnitude may be attributable to variations in receptor expression level (**Supplementary Figure 6a,c**). We decided to focus subsequent experiments on detection of VEGFA165, since it is the most abundant variant and most relevant to sensing features of the TME^77–80^. Using the most promising receptor configuration we identified (VEGFR2 NTEVp, VEGFR1 CTEVp), we observed that this receptor signals in a manner that increases with dose of ligand-encoding plasmid (**Figure 1d**). We speculate that decline of receptor output at the highest VEGFA165-encoding plasmid doses may be attributed to resource competition and/or saturation of ligand-binding sites with excess ligand (reducing receptor chain dimerization). Overall, these findings support the fundamental signaling competency of VEGF NatE MESA receptors.

We next evaluated whether VEGF NatE MESA can detect exogenous (external) recombinant VEGF. Some inducibility was observed, but the magnitude of the response to recombinant ligand was modest compared to that induced by co-expressed ligand (**Supplementary Figure 6d**). We hypothesized that the intracellular receptor components (i.e., the split TEVp parts) could provide tuning handles for improving performance. We constructed new VEGF NatE MESA receptors containing alternative split TEVp mutants that varied in split protease reconstitution propensity (interfacial energies) compared to the mutants used in the base case^21^. Since our goal was increasing receptor output, we focused primarily on mutants with lower interfacial energies than our base case (i.e., to reconstitute split TEVp more readily). We first verified that the new split TEVp variants did not ablate surface expression (**Supplementary Figure 6e**). We then tested receptor performance in response to either co-expressed or exogenous VEGF and identified a new promising receptor configuration including a wildtype NTEVp domain and 190K CTEVp domain (**Figure 1e, Supplementary Figure 6f**). In general, decreasing the interfacial energy between split TEVp components to increase reconstitution propensity increased signaling (both background and ligand-induced), but this change did not substantially impact fold induction. Interestingly, choice of NTEVp mutant modulated the magnitude of reporter expression, while choice of CTEVp mutant determined the inducibility. While it is possible that additional tuning could further improve performance, these receptors already conferred substantial ligand-inducibility and were carried forward for further development.

Expressing receptors by transient means is suitable for prototyping, but stable expression of biosensors (e.g., from a genomically integrated cassette) is desirable for many applications, and thus we next investigated how such deployment impacts receptor performance. To this end, we stably integrated VEGF NatE MESA receptor expression cassettes into the genome of HEK293FT cells using PiggyBac transposons, since they can accommodate large genetic cargo. We evaluated two strategies for integrating receptor expression cassettes and fluorescent reporters: in the first strategy, the whole system (inducible reporter, NTEVp receptor chain, and CTEVp receptor chain) was contained in a single transposon, and in the second strategy, only the receptor expression cassettes were delivered by transposons into previously engineered reporter-containing cells (**Supplementary Figure 6g**). Stable expression of receptors from PiggyBac transposons enabled sensing of exogenous ligand (**Figure 1f**), exceeding the performance previously observed when receptors were expressed by transient transfection (**Supplementary Figure 6d**). Interestingly, the receptor that includes the wild type NTEVp domain, which exhibited higher background and induced signal when expressed transiently, exhibited the best performance when expressed stably. The “all-in-one” transposon integration strategy yielded better overall performance (magnitude of induced and background signal) than did the receptor-only transposon with a pre-integrated reporter, for reasons that are not clear, but which suggest that the site and state of the genomic reporter integrations impact performance.

Because silencing is a potential challenge facing genomically integrated transgenes^81^, we next investigated whether silencing played a role in our experiments. When the high performing cell line was subcultured and retested after multiple passages over three weeks, we observed a reduction in receptor signaling and fold inductions as well as a reduction in the fraction of cells with an active reporter (**Supplementary Figure 6h**). To test the hypothesis that silencing impacted accessibility of the integrated transposons, we treated the cells with sodium butyrate (NaB), an HDAC inhibitor that is known to help re-open chromatin^82^ and is validated for use with gene circuits in mouse and human cell lines^83^. NaB treatment led to dose- and time-dependent reporter expression increase (**Supplementary Figure 6i**), as well as dose-dependent receptor surface expression increase at longer incubation times (**Supplementary Figure 6j**). Moreover, the initial fold inductions observed prior to extended passage could be recovered by NaB treatment (**Supplementary Figure 6k**). Altogether, these observations support the conclusion that transgene silencing affects apparent receptor performance, and this effect is reversible. While addressing transgene silencing is certainly an important opportunity for improving synthetic systems in general, such challenges do not preclude use of high performing biosensors even given current capabilities.

### Conversion of IL-10R

We next investigated the creation of biosensors for detecting IL-10. IL-10 is an immunoregulatory cytokine, and its dysregulation is implicated in many diseases. Overexpression of IL-10 can suppress immune responses to pathogens or other threats while down-regulation of IL-10 can mediate chronic inflammation^84,85^. IL-10 is also considered a pleiotropic molecule, with both immunosuppressive and immunostimulatory actions in different contexts^86^. Additionally, many types of cancer are associated with high local concentrations of IL-10 in the TME, including melanoma, ovarian cancer, and several lymphomas^87^. Targeting IL-10 with a cell-based therapy may permit targeting immunostimulative therapies to the tumor specifically. IL-10 is a homodimeric ligand that signals through IL-10Rα and IL-10Rβ, which are class II cytokine receptors^84^. Signaling occurs when a pair of high-affinity IL-10Rα receptor chains bind to the homodimeric ligand, eliciting a conformational change that permits two low-affinity IL-10Rβ chains to bind and form a full heterotetrameric complex which signals through the JAK/STAT signaling pathway (**Figure 2a**)^54,57,88,89^. Signaling is retained when using monomeric IL-10 mutants, suggesting that some signaling can be induced via a single α and β chain complex^57,90^. While IL-10Rα/β complexes may pre-assemble in the absence of ligand to some extent^55^, addition of ligand clearly induces an increase in homo- and hetero-association of the two receptor chains^90^. Given these mechanistic features, we hypothesized that IL-10R ECDs could be compatible with the MESA mechanism.

**Figure 2.**
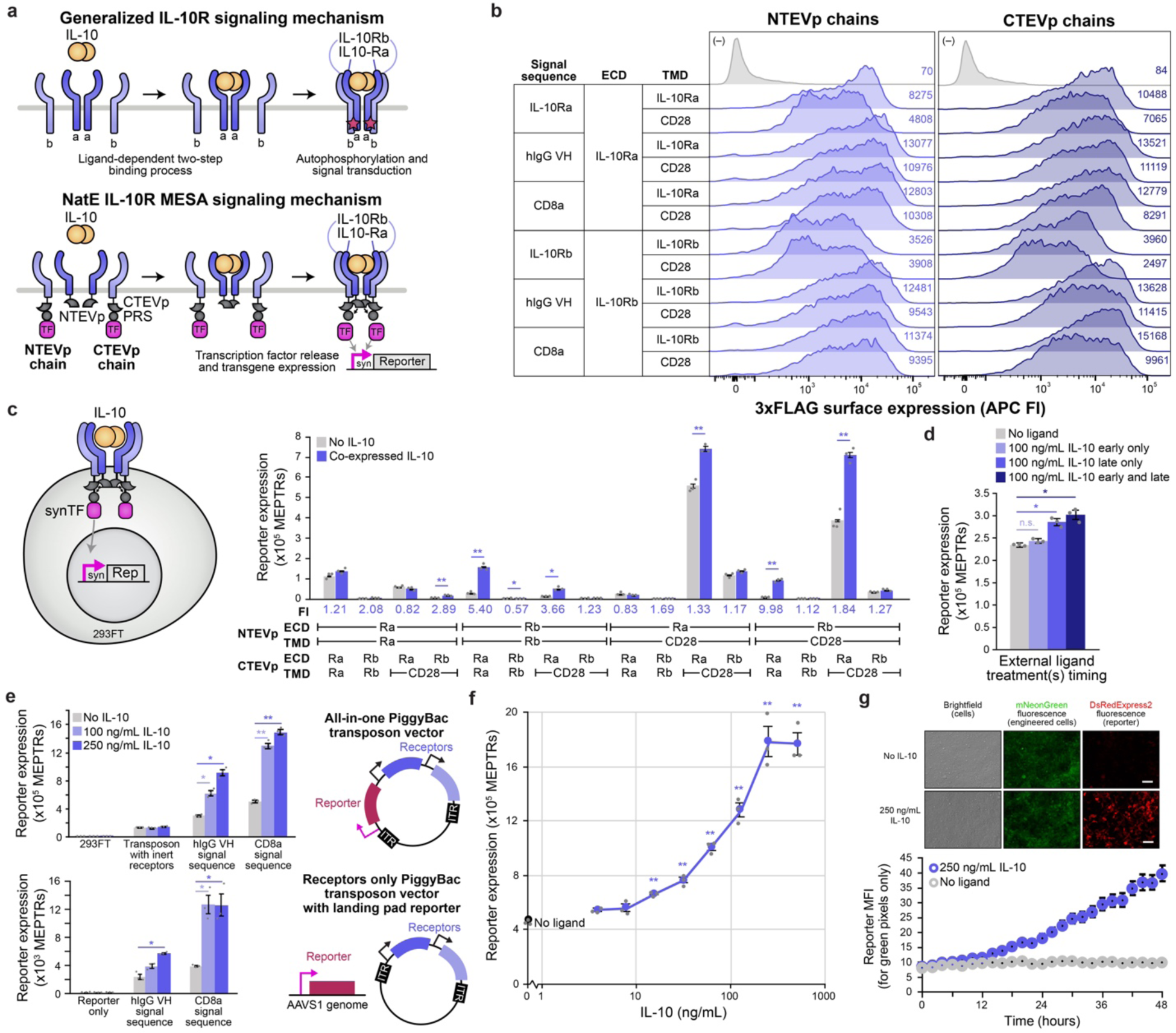
Conversion of IL-10R into an IL-10 NatE MESA receptor. **(a)** Schematic of generalized IL-10R signaling mechanism highlighting receptor interactions (top). Schematic of the proposed converted IL-10R-based NatE MESA signaling mechanism (bottom). **(b)** Surface expression of each single chain (transfected HEK293FT cells, anti-3xFLAG stain). Histograms represent transfected (fluorescent) cells and gray histograms are transfection controls (no receptor). All receptors express well on the cell surface. Mean fluorescence intensities are listed. **(c)** Functional evaluation of IL-10 NatE MESA receptor candidates (transfected HEK293FT cells) in response to autocrine human IL-10. Heteroassociative pairs exhibit ligand-inducible signaling, with fold inductions (FI: induced signal divided by background signal) shown below the plot (two-tailed Welch’s t-test results indicated above each bar pairing for * *p* < 0.05, ** *p* < 0.01, *** *p* < 0.001). Inclusion of the CD28 TMD increased background signal. All receptors include the native signal sequence that matches the ECD. **(d)** Functional evaluation of response to recombinant, exogenous ligand added at early (14 h post-transfection) and/or late (38 h post-transfection) time points (transfected HEK293FT cells). Ligand induced a significant increase in reporter expression over the non-treated condition for late treatments (two-tailed Welch’s t-test, * *p* < 0.05, ** *p* < 0.01). **(e)** Functional evaluation of IL-10 NatE MESA receptors expressed stably from the genome. HEK293FTs were modified with PiggyBac transposons encoding receptors and a reporter in the transposon (top) or encoding just receptors and introduced into a HEK293FT cell line containing a reporter integrated in the AAVS1 locus (bottom). Two receptor configurations were investigated: A CTEVp receptor with an IL-10Ra signal sequence, IL-10Ra ECD, and IL-10Ra TMD paired with a NTEVp receptor with an IL-10Rb ECD, IL-10Rb TMD, and either a hIgG VH or hCD8a signal sequence. The CD8a signal sequence NTEVp pairing exhibits more inducibility across both transposon designs. Both transposons constitutively express mNeonGreen and puromycin resistance to facilitate selection and fluorescence-based identification of engineered cells (not shown here but included in schematics in **Supplementary Figure 7f**). Data shown include all mNeonGreen+ cells. Ligand induced a significant increase in reporter expression over the non-treated condition for all receptor-expressing cell lines except for the cell line containing the inert receptors (receptors containing rapamycin-binding domains in place of IL-10R ECDs) (two-tailed Welch’s t-test indicated above each bar pairing for * *p* < 0.05, ** *p* < 0.01). **(f)** IL-10 NatE MESA dose response to recombinant human IL-10 (HEK293FT cells with genomically integrated receptor configuration: CTEVp chain with an IL-10Ra signal sequence, IL-10Ra ECD, and IL-10Ra TMD paired with an NTEVp chain with an IL-10Rb ECD, IL-10Rb TMD and hCD8a signal sequence). A dose-dependent increase in reporter expression is observed until 250 ng/mL, after which reporter expression plateaus, potentially due to saturation of receptor sites or toxicity. Ligand-induced reporter expression is significantly different from the untreated condition down to 16 ng/mL (single factor ANOVA, ** *p* < 0.01). Analogous dose responses for the same receptor configuration implemented via Sleeping Beauty transposon and when implemented via PiggyBac transposon followed by sorting for top mNeonGreen expressers are shown in **Supplementary Figure 7r**. For all bar graphs and scatter plots, bars and points represent the mean across transfected cells or transposon modified (mNeonGreen+) cells and error bars depict standard error of the mean (S.E.M.). **(g)** Microscopy images of the cell line used in panel **(f)** incubated with 250 ng/mL IL-10 for 48 h (top) and quantification of mean reporter expression over time (in engineered cells; green pixels) (bottom). After 22 h, the ligand-treated conditions are significantly different than the non-ligand treated conditions (multi-factor ANOVA, *p* < 0.05). Error bars represent the standard error across three fields of view for each time point. Abbreviations: IL-10R, interleukin-10 receptor; PRS, protease recognition sequence; TF, transcription factor; ECD, ectodomain; TMD, transmembrane domain; TEVp, Tobacco Etch Virus protease; NTEVp, N-terminal component of split, mutant TEVp; CTEVp, C-terminal component of split, mutant TEVp; APC FI, allophycocyanin fluorescence intensity; MEPTRs, molecules of equivalent PE-TexasRed; Rep, reporter; MFI, mean fluorescence intensity; FI, fold induction; Ra, IL-10Rα; Rb, IL-10Rβ.

We first evaluated expression across the full panel of design choices (**Supplementary Figure 5**). All variants were highly expressed on the cell surface (**Figure 2b**). Choice of signal sequence only modestly affected expression, and this was most pronounced for receptors with an IL-10Rβ ECD. All variants were expressed primarily as full-length receptor chains, and most receptors were expressed at similar levels within each set of NTEVp or CTEVp receptors, with the exception of IL-10Rβ receptors including the native signal sequence, which exhibited reduced total expression (**Supplementary Figure 7a**) in a manner consistent with reduced surface expression (**Figure 2b**).

We next evaluated functional performance of all receptor configurations (**Figure 2c**). We first considered a scenario in which human IL-10 was co-expressed with the receptors, and the reporter was genomically integrated (**Figure 2a**). Receptor configurations in which both chains employed the same ECD demonstrated minimal inducibility. Both background and ligand-induced signal were greatest for receptor configurations in which both ECDs were IL-10Rα, suggesting that these ECDs may mediate some pre-association. Receptor configurations pairing different ECDs (i.e., IL-10Rα on one chain and IL-10Rβ on the other chain) all demonstrated inducible signaling. The highest performing configuration (NTEVp chain containing an IL-10Rβ ECD and the CTEVp chain containing an IL-10Rα ECD) demonstrated the greatest inducibility (fold induction) and reporter output, so we selected this configuration for further analyses. Signaling required both receptor chains (**Supplementary Figure 7b-c**). Since we observed substantial differences in expression for IL-10Rβ chains as a function of signal sequences, we generated and evaluated variants of this promising receptor configuration incorporating alternative TMD and signal sequences (**Supplementary Figure 7d**). All such receptor variants demonstrated ligand-inducible signaling, and the magnitude of induced signal and sometimes fold induction could be increased by choice of signal sequences on the IL-10Rβ NTEVp chain. We opted to proceed with the CD8a signal sequence on the IL-10Rβ NTEVp chain and the IL-10Rα chain because overall magnitude of signal was highest, and we hypothesized that this property would be useful when implementing these receptors by genomic integration, for which receptor expression may be lower. As observed with the VEGF NatE MESA receptors, background signal was highest for receptors in which the CD28 TMD was included on both chains (**Figure 2b**, **Supplementary Figure 7d**), so we proceeded with the respective native TMD for the ECD on each chain.

We next evaluated whether the most promising IL-10 NatE MESA receptor can sense recombinant ligand. As was observed with VEGF NatE MESA receptors, we initially observed minimal response to recombinant ligand when IL10 NatE MESA receptors were expressed by transfection, and this conclusion held across several ligand treatment regimens (**Figure 2d Supplementary Figure 7e**). Given the improvement in performance observed when VEGF NatE MESA receptors were expressed from the genome, we hypothesized that for both systems, the complex dynamics of receptor expression, trafficking, and ligand response may render transient expression a challenging format for evaluating receptor performance in response to recombinant ligand. To address this challenge, we next stably expressed IL-10 NatE MESA receptors via PiggyBac and Sleeping Beauty transposons (**Supplementary Figure 7f**). Excitingly, we observed strong inducibility with exogenous ligand across all the designs tested. Output was greatest when the reporter was encoded in the same transposon as the receptors (consistent with our findings in the VEGF NatE MESA development), perhaps because multiple transposon copies can be integrated per cell (**Figure 2e, Supplementary Figure 7g**). Ligand-induced reporter expression was detectable by fluorescence microscopy (**Figure 2g**). In a dose response analysis, the population of biosensor-expressing cells responded differentially to IL-10 concentrations down to 16 ng/mL, with response saturating around 250 ng/mL (**Figure 2f**). The biosensor cells were also able to sense IL-10 secreted in paracrine fashion by other cells (**Supplementary Figure 7h**). Finally, we employed this framework to explore how various potential strategies for generating stable biosensor cells, including the genomic context of the biosensor-encoding cassette, impacts performance (**Supplementary Note 6**). A notable conclusion is that fold induction was relatively conserved when expressing receptors transiently or stably, suggesting that our functional assays are capturing intrinsic performance characteristics of the receptors, and that the IL-10 NatE MESA biosensor is robust to genetic context and expression level. Altogether, these data validate the fundamental feasibility and utility of this IL-10 NatE MESA receptor.

### NatE-MESA receptors can be used to construct novel, therapeutically motivated cellular functions

To demonstrate the translational utility of NatE MESA receptors, we next generated a novel engineered cellular function that could address an unmet need in cancer immunotherapy. In this proof-of-principle, we generated engineered T cells that sense IL-10 via a NatE MESA receptor and respond by activating pathways that enhance T cell cytotoxicity. First, we used the same PiggyBac transposon system used in **Figure 2e-f** to implement the biosensor, reporter, and constitutive fluorescent protein and selection marker in Jurkat T cells. The resulting cells could sense exogenous, recombinant IL-10 to induce reporter output (**Figure 3a-b**). We again observed a ligand dose-dependent increase in reporter expression that plateaued around 250 ng/mL IL-10 and produced a significant increase in reporter expression compared to background signal down to a minimal ligand dose of 32 ng/mL IL-10. Mirroring observations in HEK293FTs (**Supplementary Note 6**), sorting Jurkats for the subpopulation exhibiting greatest expression of the constitutive transposon marker (mNeonGreen) led to higher background and induced signal across all doses of IL-10 and did not change fold induction but did provide greater sensitivity for detection of IL-10 (down to 2 ng/mL), compared to the unsorted parental population (**Figure 3c**). NatE MESA can also control functional (non-reporter) outputs, which we demonstrated by implementing IL-10-induced expression of a second-generation pan-anti-ErbB CAR, T1E28z^51,91^. This circuit was validated in HEK293FTs, in which co-expression or culture with exogenous IL-10 led to a substantial increase in CAR surface expression (**Figure 3d**). When this circuit was implemented in Jurkats, CAR signaling was activated synergistically in the presence of IL-10 (recombinant or produced in trans) and antigen engagement (HER2 on co-cultured SKOV3 cells) (**Figure 3e, Supplementary Figure 8a-b**). Thus, NatE MESA receptors can be combined in series with other synthetic receptors (e.g., CARs) to integrate detection of both soluble and surface-bound cues. We also evaluated NatE MESA-driven expression of an engineered cytokine, membrane-bound IL-15 (mbIL-15), and observed an increase in mbIL-15 expression upon exposure to IL-10 (**Supplementary Figure 8c-d**)^92^. These results illustrate two general strategies for employing the IL-10 NatE MESA to control transgene expression, which could ultimately improve the specificity and safety of CAR T cell therapies by restricting expression of receptors and cytokines^93^ to sites enriched in IL-10.

**Figure 3.**
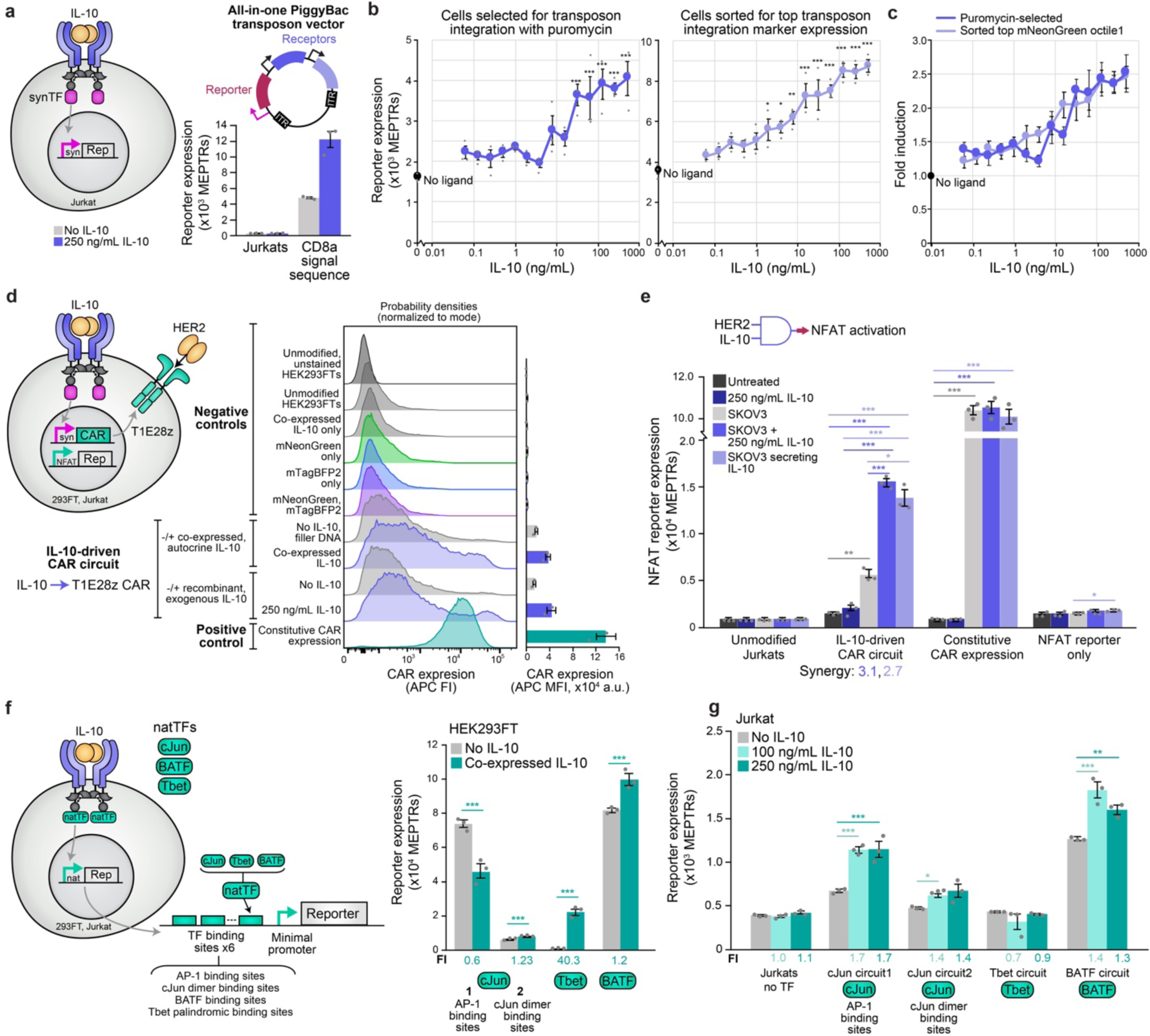
Employing IL-10 NatE MESA receptors to drive therapeutically motivated outputs. **(a)** Functional evaluation of IL-10 NatE MESA receptors integrated genomically in Jurkat T cells via PiggyBac transposon, in response to recombinant ligand. **(b)** IL-10 NatE MESA-expressing Jurkats sense IL-10 dose-dependently. Antibiotic-selected cells respond to 32 ng/mL or more, and cells sorted for high mNeonGreen (transposon-encoded, constitutive proxy) respond to 2 ng/mL or more (single-factor ANOVA, p < 0.05). Viable (DAPI-) and mNeonGreen+ cells were analyzed. Detailed schematics are in **Supplementary Figure 8j**. **(c)** Fold induction as a function of IL-10 dose (reporter signal with each IL-10 dose divided by background reporter signal with no IL-10) is conserved after sorting, suggesting that these dose response behaviors are inherent features of the receptors. With or without sorting, a maximal fold induction of 2.5 is achieved. Viable (DAPI-) and mNeonGreen+ cells were analyzed. **(d)** CAR T1E28z expression controlled by IL-10 NatE MESA receptors in HEK293FTs. Circuit components were delivered by PiggyBac transposons. Detailed schematics are in **Supplementary Figure 8j**. Negative controls include unmodified, mNeonGreen+ only (for transposon-modified cells), mTagBFP2+ only (for cells transfected with an mTagBFP2-encoding transfection control plasmid and either empty vector or an IL-10 encoding plasmid), and mNeonGreen+ and mTagBFP2+ HEK293FTs. Mean APC fluorescence intensities for 2 biological replicates are shown in the bar graph and error bars depict standard error of the mean (S.E.M.). **(e)** Implementation of an AND gate integrating soluble and surface-bound inputs. CAR expression is controlled by IL-10 NatE MESA receptors, and CAR-target binding induces signaling that induces the NFAT reporter. Circuit components were encoded on PiggyBac transposons and genomically integrated in Jurkats. Detailed schematics are in **Supplementary Figure 8j**. NFAT reporter activity increases significantly when cells are co-cultured with unmodified or IL-10-secreting SKOV3 cells (two-tailed Welch’s t-test, * *p* < 0.05, ** *p* < 0.01, *** *p* < 0.001). AND gate behavior is observed with synergy values shown below the plot. Synergy is defined in **Supplementary Note 7**. Viable (DAPI-) and mNeonGreen+ (for engineered lines, only) cells were analyzed. **(f)** IL-10 NatE MESA receptors release natural TFs in HEK293FTs. Co-expression of IL-10 with receptor chains induced natural TF-regulated reporter expression for each natural TF (two-tailed Welch’s t-test, *** *p* < 0.001). Reporter constructs were validated in **Supplementary Figure 8c-d**. **(g)** Stably expressed IL-10 NatE MESA receptors transduce IL-10 sensing to natural TF activity. Engineered Jurkats induce reporter expression for cJun constructs, but not other natural TFs (multi-factor ANOVA, * *p* < 0.05, ** *p* < 0.01, *** *p* < 0.001). Viable (DAPI-) and mNeonGreen+ cells were analyzed. Unless otherwise noted, bar graphs represent the mean across transfected or transposon-modified (mNeonGreen+) cells of three biological replicates, and error bars depict standard error of the mean (S.E.M.). Abbreviations: Rep, reporter; TF, transcription factor; MEPTRs, molecules of equivalent PE-TexasRed; CAR, chimeric antigen receptor; APC FI, allophycocyanin fluorescence intensity; MFI, mean fluorescence intensity; NFAT, nuclear factor of activated T cells; Tbet, T-box expressed in T cells; BATF, basic leucine zipper transcription factor ATF-like; AP-1, activator protein-1; FI, fold induction.

An attractive feature of engineered receptors is that their output is not limited to synthetic transcription factors but can be extended to natural transcription factors that regulate endogenous genes. As a motivating application, using natural transcription factors to boost T cell activation and prevent exhaustion to improve CAR T cell therapies is an active area of study^94,95^. Multiple transcription factors have been identified for such uses, including: c-Jun, which confers resistance to exhaustion and increased production of IL-2^94,96^; basic leucine zipper TF ATF-like (BATF), which confers resistance to exhaustion and promotes CD8+ T cell differentiation into effector T cells^97–101^; T-box expressed in T cells (T-bet), which increases the proinflammatory anti-tumor response and promotes CD4+ T cell differentiation into a T helper 1 phenotype^102,103^. Overexpressing these particular transcription factors constitutively carries oncogenic risk (for c-Jun, in particular^104^), such that restricting their activity to the tumor microenvironment could improve safety. We evaluated whether IL-10 NatE MESA receptors could be employed to sequester and release these native transcription factors to generate prototype programs that activate useful native programs in a tumor microenvironment-induced manner. We replaced the synTF with each natural TF on CTEVp-containing receptors and confirmed surface and whole cell expression (**Supplementary Figure 8e-f**). Expression of TF-containing receptors generally decreased as TF size increased. We next developed a synthetic reporter for each TF by engineering binding site arrays and pairing them with a YB_TATA minimal promoter driving DsRed-Express2 (**Figure 3f, Supplementary Figure 8g**). Reporter output increased with increasing masses of TF-encoding plasmids in HEK293FTs. Background from transfected reporter alone varied by construct because HEK293FTs express basal levels of c-Jun and BATF but not T-bet, and some reporters are subject to crosstalk with other TFs expressed in HEK293FTs^105,106^. Reporter expression increased with co-expression of IL-10 for receptors with BATF, Tbet, and cJun with a cJun dimer-specific reporter (**Figure 3f**). Reporter expression decreased with co-expression of IL-10 for the receptor with cJun and an AP-1-inducible reporter, which is responsive to other AP-1 family TFs, and this result was consistent with constitutive overexpression of free cJun (**Supplementary Figure 8g**). We hypothesize that overexpressed cJun interacts with the AP-1 proteins (cJun and cFos) basally expressed in HEK293FTs, shifting the distribution of functional dimeric TFs to more cJun homodimers than cJun-cFos heterodimers, which bind to the AP-1 response element more efficiently than do cJun homodimers^107^. When implemented stably in Jurkats via PiggyBac transposons, modest ligand-induced increases in reporter expression were observed for c-Jun and BATF but not T-bet constructs (**Figure 3g, Supplementary Figure 8h-j**), potentially due in part to the facts that receptor and reporter expression were substantially lower in Jurkats (compared to HEK293FTs) and silencing is prevalent^81^. Altogether, these proof-of-principle results demonstrate that NatE MESA receptors can be designed to regulate gene expression through the release of native transcription factors. Whether release of natural TFs from NatE MESA receptors provides adequate signal magnitude to confer meaningful functional effects via native gene regulation should be evaluated in an application specific context and is an important avenue for future investigation.

### NatE-MESA receptors can integrate information about multiple tumor microenvironment cues

The ability to engineer cell-based therapies to integrate more than one input can confer several advantages. Possible benefits include providing a desired response across varying TME cytokine signatures, or increasing safety by reducing on-target, off-tumor toxicity^14,15,108,109^. Other synthetic receptor systems have been used to perform logical evaluation of several inputs^14,15,108,109^, and receptors that signal through synthetic TF output are particularly well-suited for this purpose, but such evaluation has not yet been implemented for soluble ligands such as those found in the tumor microenvironment. We evaluated whether the two most promising NatE MESA receptors developed here (VEGF and IL-10) could be multiplexed to perform logical evaluation of cytokine inputs. Multiplexing an earlier generation of MESA receptors did not yield synergistic output^110^, and subsequent work suggests that this is attributable to both receptor and synthetic promoter characteristics^2^. We hypothesized that combining these new high-performing receptors with a minimal zinc finger domain-based synTF and cognate promoter^26^ could confer robust logical evaluation of inputs. We selected two promising receptor chain pairs for each NatE MESA receptor: for VEGF, this included the 190K CTEVp chain and both the 75S and WT NTEVp chains with native signal sequences and TMDs; for IL-10, this included the 190K CTEVp chain with the native signal sequence and TMD and 75S NTEVp chains with the hCD8a and hIgG VH signal sequences and native TMDs.

We first probed the cross-reactivity of the VEGF and IL-10 NatE MESA receptors. Most chains showed no cross-reactivity, but the VEGFR NTEVp chains showed some activity when paired with the IL-10R CTEVp chain and this was most pronounced for the VEGFR WT NTEVp (**Supplementary Figure 9a**). This cross-reactivity was reduced in the presence of ligand, potentially indicating that association between matched receptor chain pairs when bound to ligand could sequester chains to inhibit transient interactions between mismatched receptor chain pairs. Because cross-reactivity between most pairs was minimal, we concluded that VEGF and IL-10 NatE MESA receptors are likely amenable to multiplexing. We chose to evaluate each subsequent logical function with all selected VEGF and IL-10 NatE MESA receptor chain variants, including the VEGFR WT NTEVp chain, to evaluate how receptor pairs with varying degrees of orthogonality versus cross reactivity can be used to perform the intended functions.

We first employed VEGF and IL-10 NatE MESA to build an OR gate, which is defined as a circuit producing a response when either (or both) of the inputs are present, and no response when neither of the inputs are present (**Figure 4a**). To implement OR logic, both receptors were designed to release the same synTF and regulate the same output promoter. As desired, reporter expression was induced by either of the ligands, as well as both ligands together (**Figure 4b**). The magnitude of reporter expression was determined by the receptor being activated, with IL-10 inducing higher reporter levels. This trend is consistent with observations of the individual receptors, where the IL-10 NatE MESA conferred higher reporter expression levels than did the VEGF NatE MESA. Choice of IL-10R pair did not affect the function of the OR gate (reporter output with each ligand treatment), and choice of VEGFR pair did affect the magnitudes of no ligand and VEGF-only conditions (**Supplementary Figure 9b**). As observed previously in **Supplementary Figure 6f**, choosing the VEGFR WT NTEVp mutant, which has lower interfacial energy (i.e., greater propensity to reconstitute the split TEVp), increases the total signal without affecting reporter fold induction. This chain also demonstrated higher cross-reactivity with IL-10R CTEVp chains, yet it still produced successful OR gate logic here (**Supplementary Figure 9b**). Altogether, these data indicate that NatE MESA can be employed to implement OR logic, and that individual receptor characterizations can provide insights into how such gates may be tuned.

**Figure 4.**
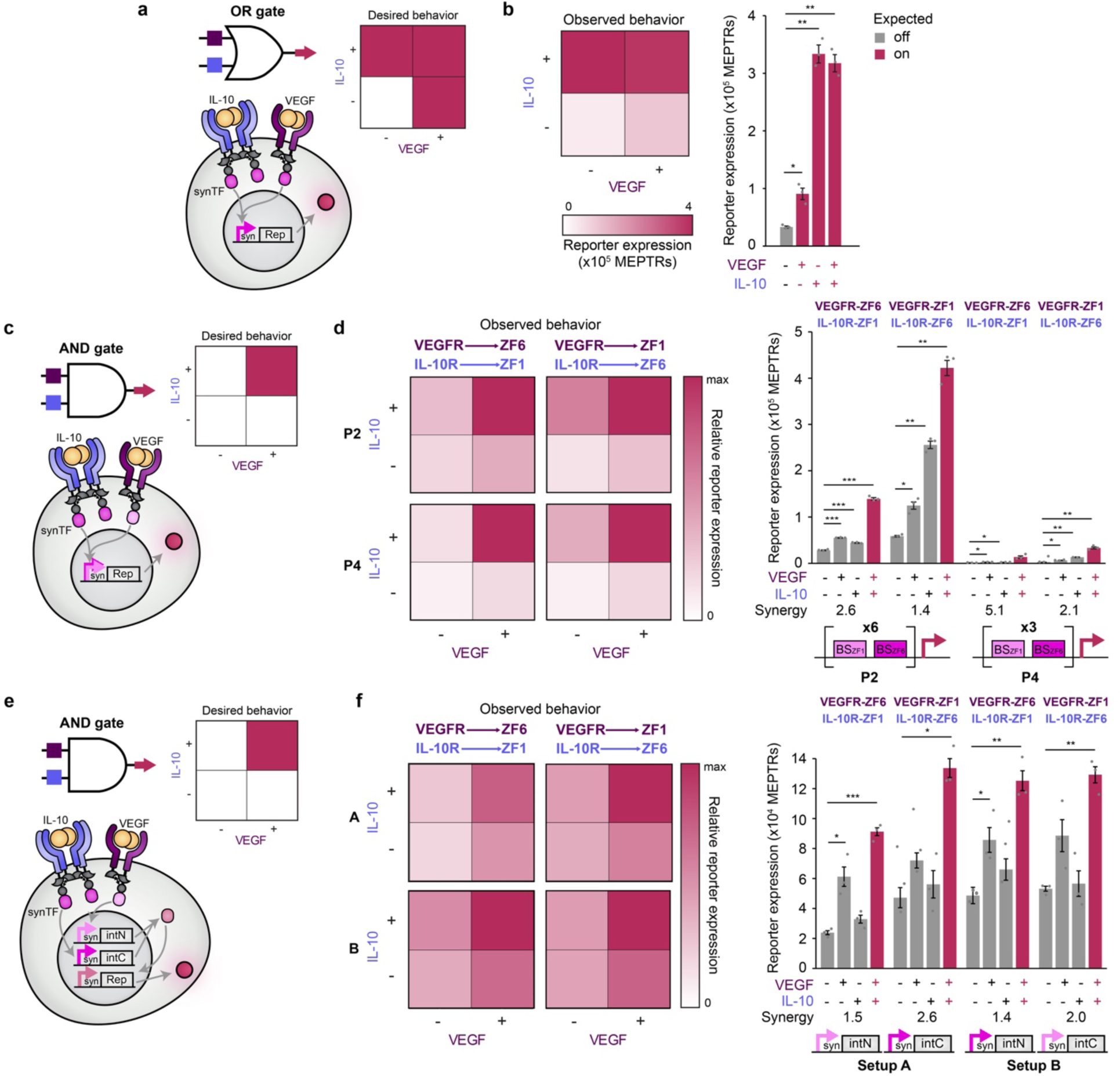
Multiplexing NatE MESA receptors to perform logical evaluation of multiple inputs. **(a)** OR gate schematic, implementation, and desired behavior. Each receptor releases the same synTF upon ligand binding, which can then bind to a cognate promoter and induce output gene expression when *either* of the inputs (IL-10 and/or VEGF) are present. **(b)** OR gate functional evaluation. IL-10 and VEGF NatE MESA receptors were expressed by co-transfection of HEK293FT reporter cells along with co-expressed ligands or empty vector DNA. The heatmap and bar graph show the same data (desired OR gate behavior) (two-tailed Welch’s t-test, * *p* < 0.05, ** *p* < 0.01, *** *p* < 0.001; insignifcant results are not displayed). See **Supplementary Figure 9b** for all configurations evaluated. **(c)** Hybrid promoter AND gate schematic, implementation, and desired behavior. Each receptor releases a different synTF upon ligand binding, which can then bind to a cognate hybrid promoter and induces output gene expression when *both* inputs are present (IL-10 and VEGF). **(d)** Hybrid promoter AND gate functional evaluation. Two promoter architectures were integrated genomically into HEK293FT-LP cells and receptors were expressed by transfection of multi-gene expression vectors (**Supplementary Figure 9c-d**), along with co-expressed ligands or empty vector. Heatmaps and bar graphs show the same data, and each heatmap scale is normalized to the maximum value within that heatmap. Synergy values are shown below the plot and this metric is defined in **Supplementary Note 7**. Some receptor-synTF pairings showed better synergy than others, although all displayed AND gate behavior (two-tailed Welch’s t-test, * *p* < 0.05, ** *p* < 0.01, *** *p* < 0.001; insignificant results are not displayed). See **Supplementary Figure 9e** for all configurations evaluated. **(e)** Split-intein synTF AND gate schematic, implementation, and desired behavior. Each receptor releases a different synTF upon ligand binding, which induces expression of one half of a split-intein synTF (activation domain fused to intN, DNA-binding domain fused to intC^111^), which then reconstitutes and induces output gene expression when *both* inputs are present (IL-10 and VEGF). **(f)** Split-intein synTF AND gate functional evaluation. Reporter setups were integrated genomically into HEK293FT-LP cells and receptors were expressed by transfection of multi-gene expression vectors (**Supplementary Figure 9c-d**), along with co-expressed ligands or empty vector. Heatmaps and bar graphs show the same data, and each heatmap scale is normalized to the maximum value within that heatmap. Some configurations showed better synergy than others, and most configurations displayed AND gate behavior (two-tailed Welch’s t-test, * *p* < 0.05, ** *p* < 0.01, *** *p* < 0.001; insignificant results are not displayed). See **Supplementary Figure 9f** for all configurations evaluated. The VEGF receptor pair used in this figure is a NTEVp receptor chain with VEGFR2 signal sequence, ECD and TMD, 75S NTEVp and a CTEVp receptor chain with VEGFR1 signal sequence, ECD and TMD, 190K CTEVp. The IL-10 receptor pair used in this figure is a NTEVp receptor chain with IgGVH signal sequence, IL-10Rb ECD and TMD, 75S NTEVp and a CTEVp receptor chain with IL-10Ra signal sequence, IL-10Ra ECD and TMD, 190K CTEVp. Bar graphs represent the mean of three biological replicates of transfected cells, and error bars depict standard error of the mean (S.E.M.). Abbreviations: MEPTRs, molecules of equivalent PE-TexasRed; Rep, reporter; synTF, synthetic transcription factor; PX, promoter design X; intN, N-terminal split intein fragment; intC, C-terminal split intein fragment; VEGF, vascular endothelial growth factor; IL-10, interleukin-10.

We next built an AND gate with NatE MESA receptors. An AND gate is defined by exhibiting synergy (i.e., the output when both inputs are present is larger than the sum of the outputs in response to each individual input). In our first AND gate design, each receptor releases a different synTF which binds to a hybrid promoter containing binding sites for both synTFs (**Figure 4c**)^110^. We designed several promoter architectures based on previous work^26^, genomically integrated them into a HEK293FT landing pad^31^, and evaluated synergistic activation via synTFs (**Supplementary Figure 9c**). Although all designs exhibited synergy, we moved forward with P2 and P4, which exhibited both high synergy and high reporter output. To evaluate each receptor individually, we created multi-gene expression vectors (MGEVs) for each receptor, including the NTEVp and CTEVp chains each driven by a separate constitutive promoter as well as a constitutive fluorescent protein proxy (**Supplementary Figure 9d**). This setup resulted in 1:1 gene copies of each chain (NTEVp and CTEVp) for each receptor (**Supplementary Figure 6**). We transfected MGEVs into the reporter cell lines containing the P2 and P4 promoters, along with vectors encoding ligands. All configurations of receptors, synTFs, and cognate promoters yielded synergistic output, even configurations employing the VEGFR WT NTEVp chain which demonstrated higher cross-reactivity with IL-10R chains (**Figure 4d, Supplementary Figure 9e**). In general, P4 exhibited more synergy than P2. We hypothesize that this is due to the mechanism by which these synTFs cooperatively recruit transcriptional machinery^26^; since P2 has more binding sites for the synTFs, each input (and corresponding synTF) can induce more transcriptional activation. This phenomenon is most pronounced for the receptor that releases the synTF for which binding sites are closer to the minimal promoter (in this case, ZF6). Interestingly, variations upon receptor design had little effect upon AND gate performance, which was mostly driven by synTF choice and promoter design (**Supplementary Figure 9e**). In sum, these data demonstrate that NatE MESA can implement AND logic and tuning at the level of synTF and promoter design rather than receptor design was most impactful.

To investigate whether the performance of our AND gates could be improved, we next implemented an alternative strategy to reduce output in response to single ligands (and improve synergy) by making the output from each receptor incapable of driving any reporter output. To this end, we employed split intein domains to separate a synTF into its activation and DNA-binding domains^27^. Each receptor releases a different synTF (ZF1 or ZF6), which can then induce the expression of one half of the split-synTF in the presence of its corresponding ligand (**Figure 4e**). We engineered a new landing pad^31^ reporter cell line incorporating the intermediate synTF-responsive promoters driving expression of the split-synTF halves, and we validated its function (**Supplementary Figure 9f**). When ligand-receptor configurations were evaluated as before, this design also exhibited synergy in some cases, although synergy was generally less than that observed for the hybrid promoter AND gate design (**Figure 4f**). In general, moving to a split synTF design successfully reduced output under the IL-10 only condition, but no similar improvement was observed for the VEGF only condition. We hypothesize that this could be explained by the higher ligand-independent background signal generally observed from the IL-10 NatE MESA (compared to the VEGF NatE MESA); due to this effect, some of the IL-10 NatE MESA-associated split synTF component is always available to complement the half released from the VEGF NatE MESA. We hypothesize that reducing background from the IL-10 NatE MESA receptors could increase AND gate synergy. An additional consideration is that the promoter controlling the reporter is driven by a single synTF, such that even a small amount of reconstituted synTF can bind near the minimal promoter and cooperatively activate transcription, whereas in the case of the hybrid promoter in **Figure 4d**, any one synTF is not capable of such cooperative activation. We again observed that variations upon receptor design had little effect upon AND gate performance compared to downstream circuit design choices (**Supplementary Figure 9g**). These results combined with those obtained for the hybrid promoter AND gate evaluations underscore the conclusion that level matching between receptor output levels (both background and induced signal) is tunable and is important for successfully achieving the desired circuit function^110^. Such a perspective also enables one to identify future design refinements that could build upon these foundational characterizations to implement suitable designs for diverse applications.

## DISCUSSION

In this study, we explored the conversion of natural receptors into synthetic biosensors that sense and respond to soluble, therapeutically relevant extracellular cues. One key lesson is that such conversion was nontrivial—not all receptor conversions were straightforward or yielded functional biosensors at all. Generally, we focused on the conversion of receptors with well-understood biophysical mechanisms, and most of our observations could be interpreted (if not predicted) using insights from receptor function. Conversion success was driven primarily by compatibility between natural receptor mechanisms and the MESA mechanism. Hetero-associative receptors were most successfully converted because MESA signaling is regulated by association of two different receptor chains. In most cases, pre-association of receptor chains (interactions that were not impacted by ligand binding) were undesired and often led to high ligand-independent signaling (e.g., IL-10Ra, TGFBR1, and TNFR1) when complementary signaling domains were paired with the same ECD. However, for receptors that pre-associate and undergo a state change upon ligand binding, pre-associations may be favorable, such as in the case of VEGFRs, where ligand-independent association is naturally observed, yet receptor pre-association did not result in ligand-independent signaling. We hypothesize that the same mechanism that prevents VEGFR kinase domain autophosphorylation in the absence of ligand^71,112^ also prevents MESA intracellular interactions, which are then enabled by the conformational change brought about upon ligand binding to the ECD. Within the successful conversions, the best performing NatE MESA receptors were receptors containing natural TMDs. In general, exploring alternative N-terminal signal peptide sequences sometimes helped in improving receptor trafficking to the cell surface and increasing expression, and in turn increasing receptor signaling, but no change in signal peptide rescued non-inducible receptor designs in our study. TMD choice, on the other hand, modulated receptor behavior and seemingly produced chain associations not observed in nature. For example, by using a CD28 TMD in TNF NatE MESA, we were able to employ a low affinity receptor variant (TNFR2 ECD) to produce an inducible signal, which is not normally observed for soluble TNF and TNFR2 (**Supplementary Note 4, Supplementary Figure 10**)^58^. Variations of other design choices, much like signal sequence choice, only resulted in modulation of overall receptor-induced signal magnitude without much effect on fold-inducibility, although exploring such variations can be useful for tuning the performance of a functional, base-case receptor to meet a specific need. We also found that exploring downstream circuits to process outputs from a given functional receptor can be a fruitful avenue for tuning overall system performance.

Though the VEGFR and IL-10R conversions were most successful and most deeply characterized here, we also evaluated conversion of TGF-βR and TNFR, which were less successful (**Supplementary Notes 4 and 5, Supplementary Figures 10 and 11**). The TGF-β NatE MESA exhibited low receptor chain expression, even with alternative signal sequences and TMDs, and extended JMDs, and no tested configuration yielded inducible signaling. The TNF NatE MESA produced minimally inducible biosensors when chains contained a CD28 TMD or split TEVp variants with high reconstitution propensity. Improvement of TNF NatE MESA into a higher performing biosensor may require tuning the interactions between intracellular domains (potentially by adding rotation via adding amino acids, akin to a method used to tune GEMS receptors^7^, or other intracellular linker variations) to bring parts together in a geometry compatible with split TEVp reconstitution. Alternatively, improvements observed with the more easily reconstituted split TEVp variants (**Supplementary Figure 10**) would likely require introduction of a new mechanism to reduce background signal. These examples highlight the importance of achieving high expression (TGF-β) and exploring alternative receptor design choices (TNF) in the conversion process. With the advancement of computational tools for predicting structures of novel proteins, it may be possible to gain additional insights into the structural basis of such challenging receptor conversions and identify opportunities for improvement. We summarize the experience gleaned through this study as a set of steps that can guide natural receptor conversion into a functional synthetic MESA receptor in **Supplementary Figure 12** and **Supplementary Note 8.**

In a key demonstration of the utility of NatE MESA receptors, we showed for the first time, to our knowledge, that mammalian cells can be engineered to integrate the sensing of multiple soluble, environmental protein ligands via synthetic receptors. We also built genetic circuits to process these inputs by performing various potentially useful operations, including implementing Boolean logic (a framing which aligns with clinical understanding of microenvironmental features) and rewiring transcriptional output to yield immunotherapy-relevant activities. An important enabling property is that NatE MESA receptors can be multiplexed in parallel (by integrating synTF outputs via logic circuits) as well as in series with other synthetic receptors (i.e., CARs). Learnings from previous work^110^ as well as general improvements in technology, including the fact that these high performing NatE MESA receptors are robust to variations in stoichiometry (compared to earlier designs^12^) and the availability of customizable toolkits of transcription parts^26^, enabled us to achieve AND gate behavior without extensive tuning. This series of demonstrations also exemplifies a longer-term trend in that the use of synthetic receptors to achieve cellular engineering goals is enabled by integration with technologies for downstream information processing^2^.

NatE MESA receptors are attractive for therapeutic translation, and this study identifies opportunities for future development. First, the receptors and transgene output modules function well when genomically integrated, and this function holds across multiple, distinct human cell types (e.g., adherent HEK293FTs and suspension-cultured Jurkat T cells). It will be important to evaluate the extent to which this trend holds across recipient cell types and delivery modalities. Preliminary evaluations of receptor performance across different populations within genomically engineered cell lines suggest that NatE MESA receptors are robust to genetic context (**Supplementary Note 6**). Second, the majority of the NatE MESA ectodomains are human-derived, which could minimize immunogenicity^113^. These properties may be complemented and improved by ongoing work including creating orthogonal human proteases^114^, and humanization of exogenous sequences through protein engineering^115^. Evaluating immunogenicity is complex and probably context-specific, and evaluating the need for de-immunization may be best addressed within a given application. Third, we demonstrate that NatE MESA receptors can be used to implement illustrative, proof-of-principle circuits that could ultimately support and enhance CAR T cell function. Future avenues for investigation include combining these NatE MESA and CAR designs in preclinical animal models to identify, and if needed address, needs for increasing and shifting bioactivity. Fourth, although NatE MESA receptors signal through orthogonal pathways, there remains a potential for interaction (or competition) with the native receptors from which NatE MESA are derived. Potential solutions include a careful matching of NatE MESA and cell type to avoid such problems, or systematically mutating NatE MESA domains to prevent interactions with endogenous receptors but maintain interactions with MESA receptor partners. Finally, although the NatE MESA developed here transduce ligand sensing into changes in gene expression, it is conceivable that split TEVp could be replaced with other split enzymes (with reconstitution tuned if needed^116^) to transduce ligand sensing into enzymatic activity. Altogether, this NatE MESA receptor conversion study may inform the development of novel, tunable biosensors for programming cellular responses to diverse physiological targets of interest.

## DATA AVAILABILITY

The data that support the findings of this study will be published along with the final version of this manuscript. The plasmids generated in this study will be deposited with and distributed by Addgene at the time of publication, including complete and annotated sequence files, at https://www.addgene.org/Joshua_Leonard/.

## ACKNOWLEDGEMENTS

We would like to thank the Neha Kamat lab for use of an Azure c280 imager for western blot analyses and members of the Leonard lab for useful discussions throughout the planning, experimental, and analysis phases of this project. We would also like to thank Gauri Bora for assistance with plasmid cloning in this study.

## AUTHOR CONTRIBUTIONS

H.I.E. and J.N.L. conceived the initial project. H.I.E., A.C., M.L.E., and A.H.M. created reagents and designed and performed experiments. H.I.E., A.C., M.L.E., A.H.M., and W.K.C. analyzed the data. H.I.E., A.C., and J.N.L. wrote the manuscript and drafted the figures. J.N.L. supervised the work. All authors edited and approved the final manuscript.

