## Supplementary Information for "Conversion of natural cytokine receptors into orthogonal synthetic biosensors"

**Contents:**

Supplementary Figures 1–12

Supplementary Tables 1–4

Supplementary Notes 1–9

References cited in this document

### SUPPLEMENTARY FIGURES

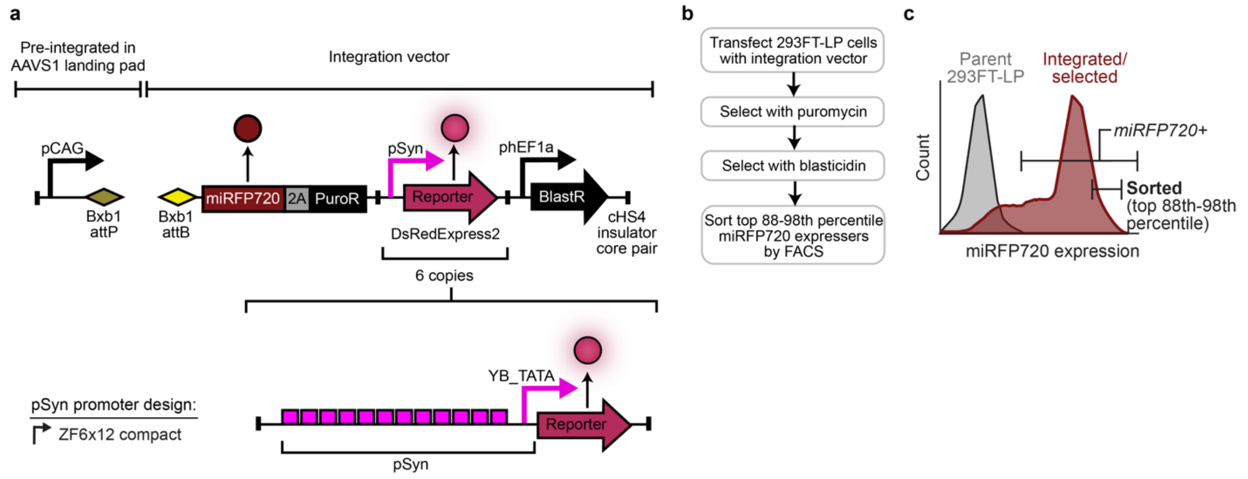


**Supplementary Figure 1. Generation of stable, genomic reporter cell lines.**
**(a)** Schematic of landing pad integration vector cargo and cell line development workflow for generating a reporter cell line. Parental HEK293FT-LP cell line contains a pre-integrated landing pad in the AAVS1 locus with a CAG promoter upstream of an attP Bxb1 recognition site[1]. Successful integration of cargo results in CAG-driven expression of the first gene in the integration vector downstream of the attB Bxb1 recognition site, here miRFP720 and the puromycin resistance gene. A secondary selection marker, the blasticidin resistance marker, is also included as a separate transcriptional unit regulated by a constitutive promoter as the last unit of the integration vector. The other transcriptional units encode one copy of a synTF-mediated promoter driving expression of a DsRedExpress2 reporter gene. **(b-c)** Parental HEK293FT-LP cells were transfected with each integration vector and a plasmid that encodes Bxb1 expression, selected first with puromycin for one week, then with blasticidin for another week before sorting for top miRFP720 expressers. Abbreviations: pCAG, cytomegalovirus (CMV) enhancer fused to the chicken beta-actin promoter (CAG); miRFP720, monomeric infrared fluorescent protein 720; 2A, P2A peptide; PuroR, puromycin resistance gene; phEF1a, human elongation factor alpha promoter; BlastR, blasticidin resistance gene; cHS4, chicken hypersensitivity site 4 insulator; LP, landing pad; ZFX, zinc finger X.

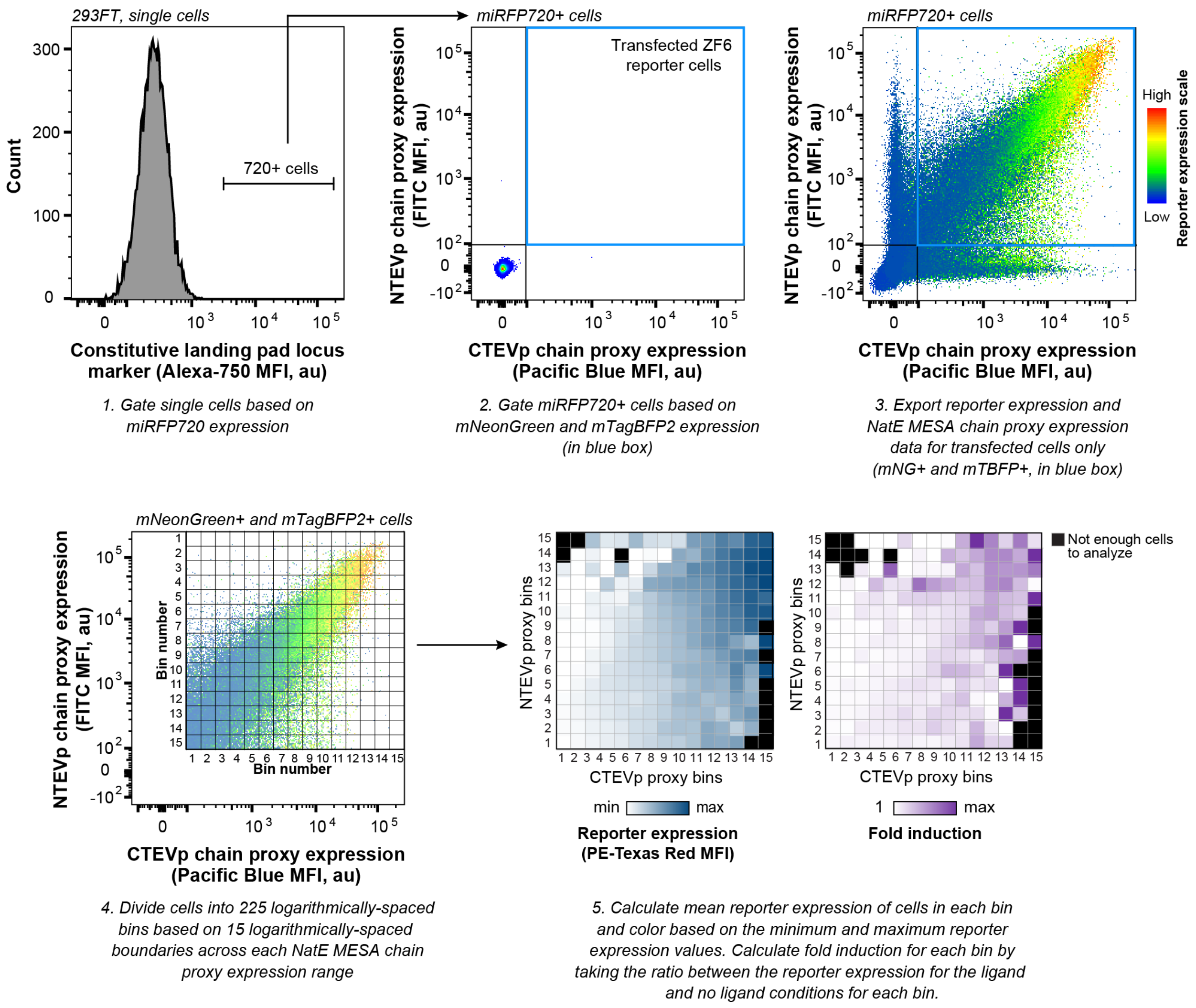


**Supplementary Figure 2. Poly-transfection workflow summary.**
Workflow for poly-transfection data analysis. First, single cells were gated based on miRFP720 fluorescence to identify cells with an active Landing Pad locus containing the reporter construct. Reporter cells were co-transfected with separate plasmids containing the NatE MESA chains co-expressed with a fluorescent proxy (mNeonGreen for NTEVp chains, mTagBFP2 for CTEVp chains). Data for reporter and proxy expression was then exported from the double transfected population (mNeonGreen+ and mTagBFP+) quadrant highlighted in blue. These cells were divided into 225 logarithmically spaced bins based on 15 logarithmically spaced boundaries across NTEVp chain and CTEVp chain proxy expression range. Finally, mean reporter expression was calculated for each bin and fold induction was determined as the ratio between the reporter expression in the ligand case vs the no ligand case.

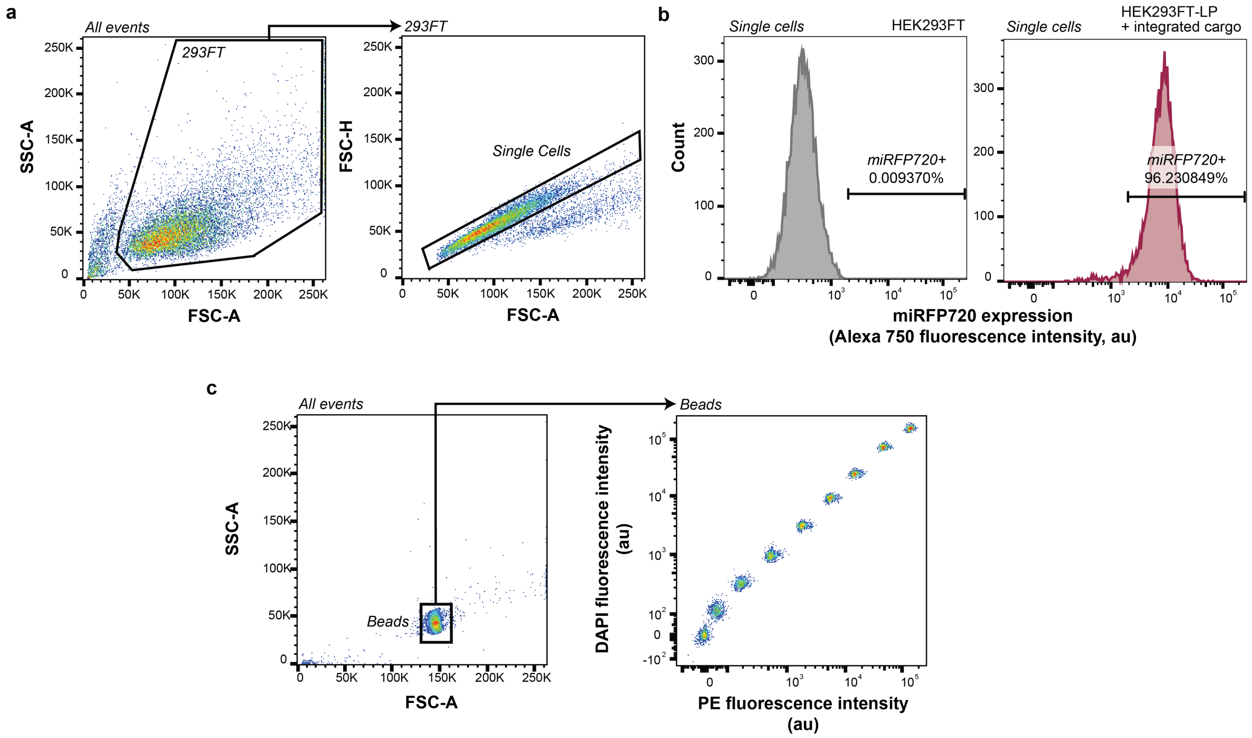


**Supplementary Figure 3. General flow cytometry gating strategies.**
**(a)** Illustration of the flow cytometry gating strategy used to identify single HEK293FT cells for a representative sample of cells. **(b)** Illustration of the flow cytometry gating strategy used to identify cells that are expressing miRFP720 from the landing pad locus (for identifying reporter cells with an accessible landing pad locus cells). A gate is drawn on unmodified HEK293FT cells (left) to include <0.1% of cells in the Alexa750 fluorescence channel. This gate encompasses most landing pad-modified cells (right). A similar approach is used with the FITC channel to identify transposon modified HEK293FT cells that are constitutively expressing mNeonGreen (not shown). **(c)** Calibration of fluorescence intensities to absolute units requires inclusion of a sample of Spherotech UltraRainbow Calibration Particles (URCP) in each experiment. These beads have nine fluorescent bead populations. Beads are identified based on the FSC-A vs. SSC-A profile (left). For each experiment, two fluorescent channels were used to identify all nine bead populations (right). The mean fluorescence intensities (MFIs) of each population in the relevant channel(s) are exported and plotted against manufacturer-provided absolute values of fluorophores per bead for each population (for example, Molecules of Equivalent Fluorescein MEFLs for mNeonGreen, Molecules of Equivalent PE-TexasRed MEPTRs for DsRedExpress2). To generate the calibration curve, a linear regression was performed with the constraint that the y-intercept equals zero. This calibration is done for each experiment, and then exported MFI values (which have arbitrary fluorescence units) are converted to absolute units using the multiplier obtained from the regression. Error from the linear regression is also appropriately propagated. Abbreviations: FSC-H, forward scatter height; FSC-A, forward scatter area; SSC-H, side scatter height; SSC-A, side scatter area; miRFP720, monomeric infrared fluorescent protein 720; au, arbitrary units; LP, landing pad.

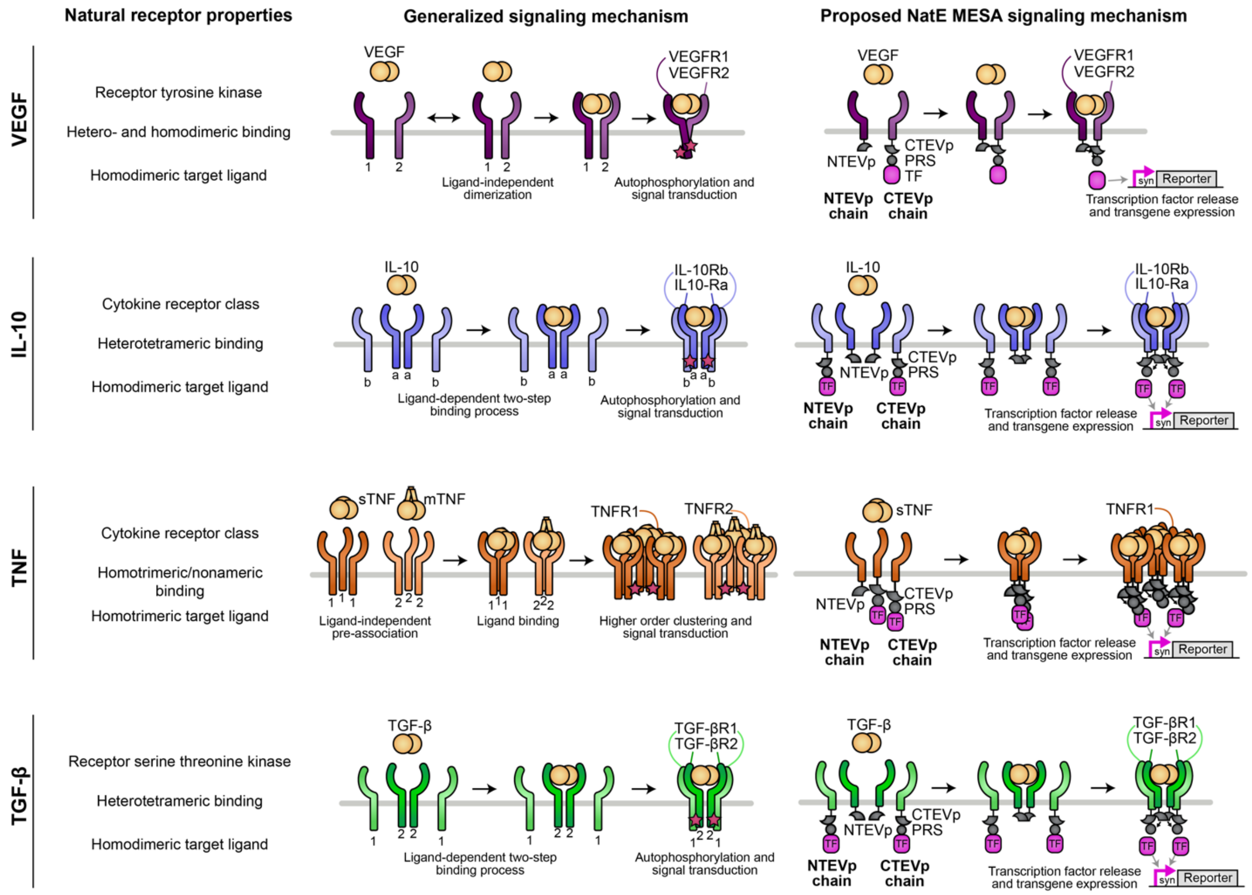


**Supplementary Figure 4. Summary of human receptors explored in this study.**
From left to right, classification of each selected natural receptor system and its receptor and ligand multimeric states, overview of what is currently known about the natural receptor mechanisms for each system, and proposed NatE MESA mechanism for each system. Abbreviations: VEGF, vascular endothelial growth factor; IL-10, interleukin-10; TNF, tumor necrosis factor; TGF-B, transforming growth factor beta; TEVp, Tobacco Etch Virus protease; NTEVp, N-terminal component of split, mutant TEVp; CTEVp, C-terminal component of split, mutant TEVp; PRS, protease recognition sequence.

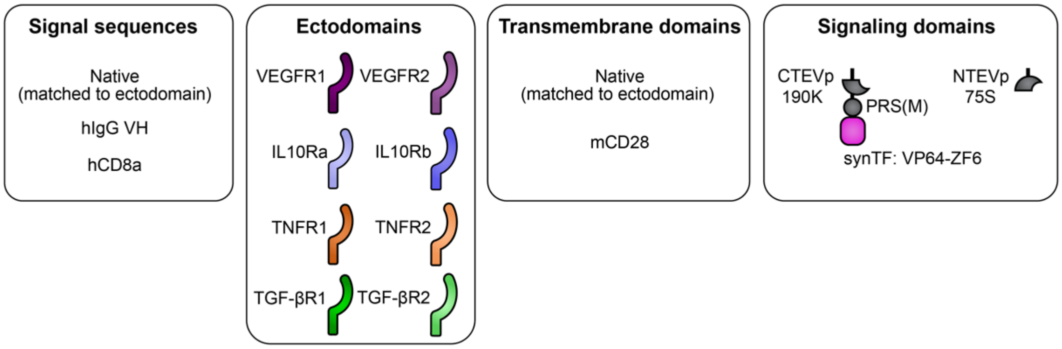


**Supplementary Figure 5. Summary of the synthetic receptor design space explored across all four NatE MESA receptors included in this study.**
Overview of the consistent design choices explored for all receptors. Some receptors employed additional design choices as needed (including split TEVp mutants and human CD28 TMD variations), which are not covered here. Abbreviations: VEGFR, vascular endothelial growth factor receptor; IL-10R, interleukin-10 receptor; TNFR, tumor necrosis factor receptor; TGF-BR, transforming growth factor beta receptor; TEVp, Tobacco Etch Virus protease; NTEVp, N-terminal component of split, mutant TEVp; CTEVp, C-terminal component of split, mutant TEVp; PRS(M), protease recognition sequence with methionine in P1' position; synTF, synthetic transcription factor; ZFX zinc finger X; mCD28, murine CD28.

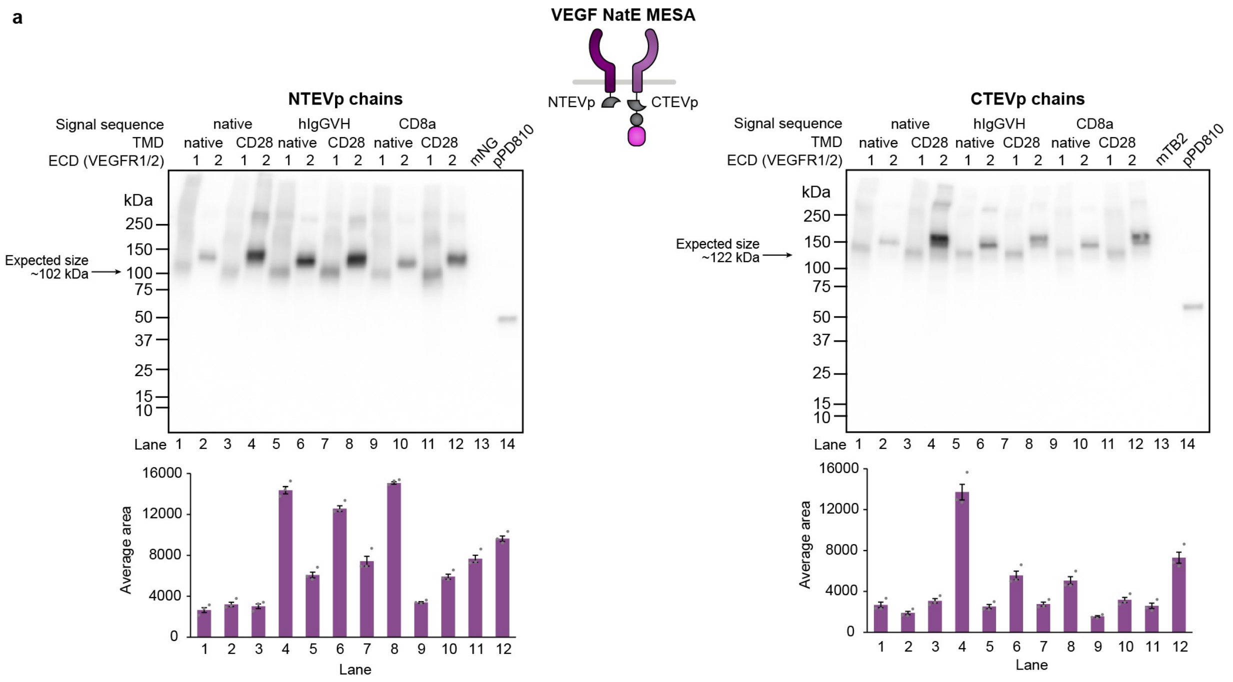


**Supplementary Figure 6. Supplementary information for the conversion of VEGFR.**
**(a)** Whole-cell expression of each VEGF NatE MESA receptor was measured via western blotting of a 3xFLAG epitope tag fused to the N-terminus of receptors. Soluble mNeonGreen (mNG) or mTagBFP2 (mTB2) were included as negative controls because each receptor chain-expressing plasmid is a poly-transfection vector that also contains either a constitutively expressed mNG or mTB2 in the backbone. A rapamycin-sensing MESA receptor, pPD810 was included as an internal positive control for 3xFLAG-tagged protein expression[2]. Quantification of expression levels from bands in each western blot is shown below each gel image. To quantify band intensity, western blot images were imported into ImageJ and analyzed using the analyze gel feature. Three images with different exposure times and no saturated pixels were used for analysis. Intensities of all bands (area under the curve) in each lane excluding the band representing the pPD810 control were summed and averaged across three exposure times. Error bars depict the standard error of the mean (S.E.M.) for these three measurements. Abbreviations: VEGFR, vascular endothelial growth factor receptor; ECD, ectodomain; TMD, transmembrane domain; TEVp, Tobacco Etch Virus protease; NTEVp, N-terminal component of split, mutant TEVp; CTEVp, C-terminal component of split, mutant TEVp.

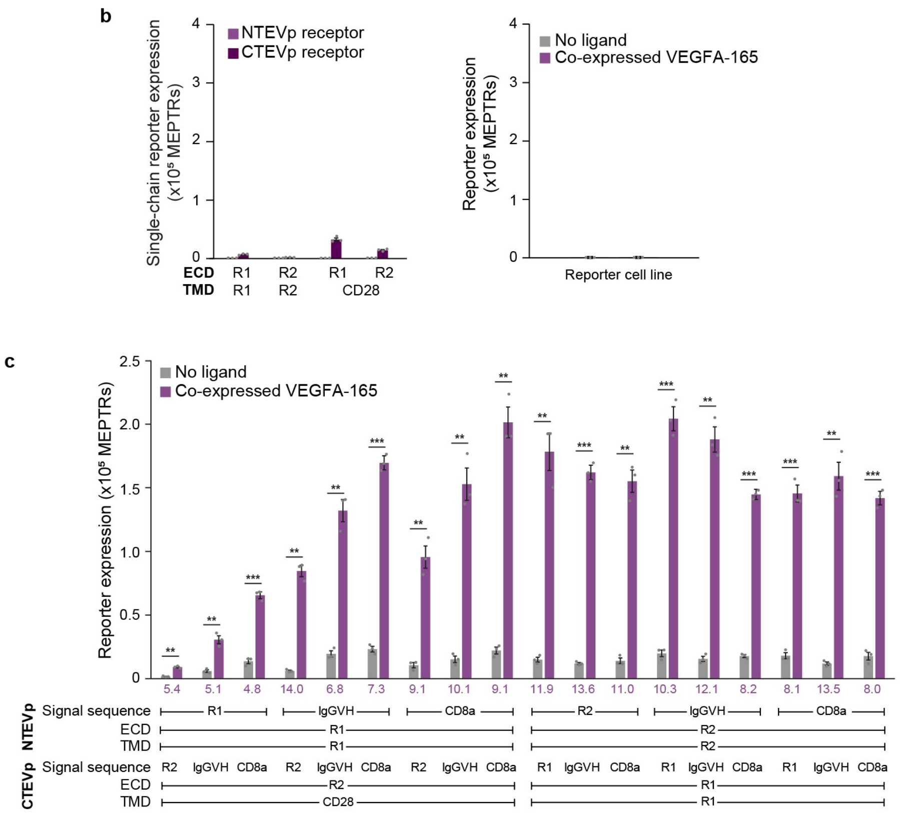


**Supplementary Figure 6. Supplementary information for the conversion of VEGFR.**
**(b)** The reporter cell line was transfected with plasmids encoding each receptor chain individually to assess single receptor-induced reporter expression in the absence of ligand (left). No reporter expression is observed with NTEVp chains, which do not contain a synTF. Minimal reporter expression is observed with CTEVp chains (e.g., from spontaneous release of the synTF from the membrane). In addition, the reporter cell line was transfected with plasmids expressing VEGF only to show that no reporter expression is induced only in the presence of co-expressed VEGF (right). The y-axis of each plot is scaled to match that of **Figure 1c**. **(c)** Evaluating the effect of receptor signal sequence on reporter expression with receptors expressed via transfection and co-expressed VEGF. Two inducible pairs from the initial panel were selected and pairwise combinations of receptors with varying signal sequences were evaluated in the presence and absence of VEGF. Fold induction values are indicated below each pair of bars for each receptor chain combination, indicating the ratio between reporter expression in the presence and absence of ligand. Both receptor pairs were inducible across all signal sequence pairings (two-tailed Welch's t-test: * $p < 0.05$, ** $p < 0.01$, *** $p < 0.001$) and signal sequence choice impacted both background and ligand-induced reporter expression (multi-factor ANOVA, $p < 0.05$). Each bar represents the mean of transfected cells across three biologic replicates and error bars indicate the standard error of the mean (S.E.M). Abbreviations: VEGFR, vascular endothelial growth factor receptor; ECD, ectodomain; TMD, transmembrane domain; TEVp, Tobacco Etch Virus protease; NTEVp, N-terminal component of split, mutant TEVp; CTEVp, C-terminal component of split, mutant TEVp; R1, VEGFR1; R2, VEGFR2; MEPTRs, molecules of equivalent PE-TexasRed.

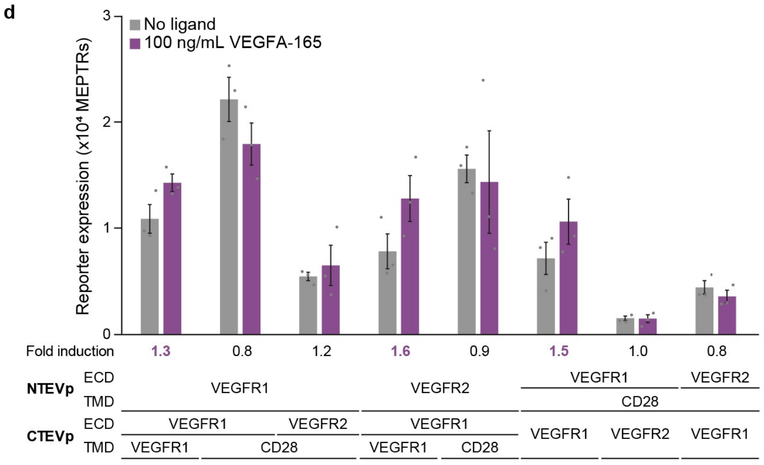


**Supplementary Figure 6. Supplementary information for the conversion of VEGFR.**
**(d)** Reporter expression with receptors expressed via transfection and cells treated with recombinant VEGF. The eight inducible pairs from the functional evaluation performed using co-expressed VEGF were selected for further analysis here. All receptors contain their natural signal sequence. Fold inductions are shown below each pair of bars. Each bar represents the mean of transfected cells across three biologic replicates and error bars indicate standard error of the mean (S.E.M). No statistical significance was found with a Welch's T-test between the no ligand and VEGFA conditions for each receptor combination. Abbreviations: VEGFR, vascular endothelial growth factor receptor; ECD, ectodomain; TMD, transmembrane domain; TEVp, Tobacco Etch Virus protease; NTEVp, N-terminal component of split, mutant TEVp; CTEVp, C-terminal component of split, mutant TEVp; MEPTRs, molecules of equivalent PE-TexasRed.

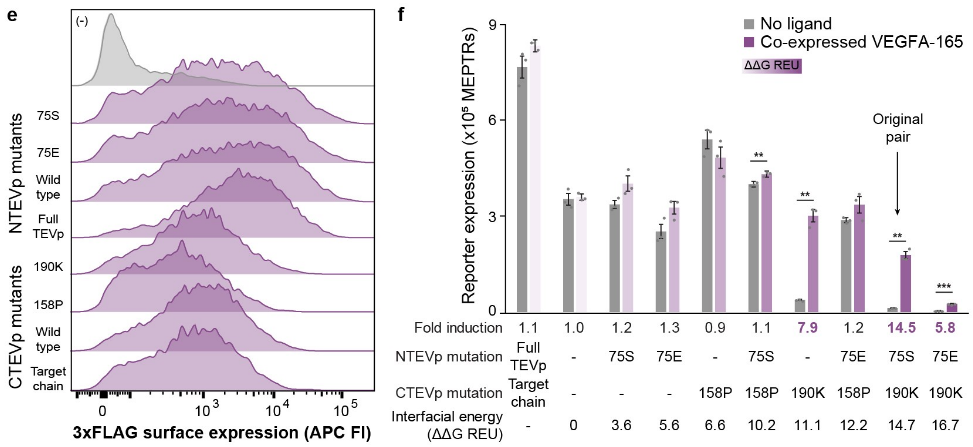


**Supplementary Figure 6. Supplementary information for the conversion of VEGFR.**
**(e)** Surface expression of receptor chains with different NTEVp and CTEVp mutant domains was measured via immunohistochemistry. Receptors which employ only the *trans*-cleavage MESA mechanism (full TEVp on one chain, protease recognition sequence and synTF on the other chain) were included as a reference to show that splitting the TEVp does not ablate surface expression. Histograms included all transfected cells based on expression of a constitutive fluorescent protein in the receptor-encoding plasmid backbone. **(f)** Transfected receptor combinations with varying split TEVp mutants yielding a range of interfacial energies (Rosetta Energy Units, REU)[3] were tested for functionality with co-expressed and exogenous VEGF. Darker bars indicate higher interfacial energy, resulting in lower propensity for TEVp reconstitution. High performing receptors with co-expressed ligand (left) also detected exogenous ligand (right). Fold inductions are shown below each pair of bars. Results for two-tailed Welch's t-test are indicated above corresponding bars for significant pairs only (** $p < 0.01$, *** $p < 0.001$). The difference between the 75S/190K and WT/190K pairs is statistically significant (** $p < 0.01$) as determined via a multi-factor ANOVA and Tukey's HSD Test. Interfacial energy shows a significant effect on reporter expression across ligand treatments (multi-factor ANOVA, *** $p < 0.001$). Each bar represents the mean of transfected cells across three biologic replicates and error bars indicate standard error of the mean (S.E.M). Abbreviations: VEGFR, vascular endothelial growth factor receptor; ECD, ectodomain; TMD, transmembrane domain; TEVp, Tobacco Etch Virus protease; NTEVp, N-terminal component of split, mutant TEVp; CTEVp, C-terminal component of split, mutant TEVp; R1, VEGFR1; R2, VEGFR2; MEPTRs, molecules of equivalent PE-TexasRed; APC FI, allophycocyanin fluorescence intensity; REU, Rosetta energy units.

g

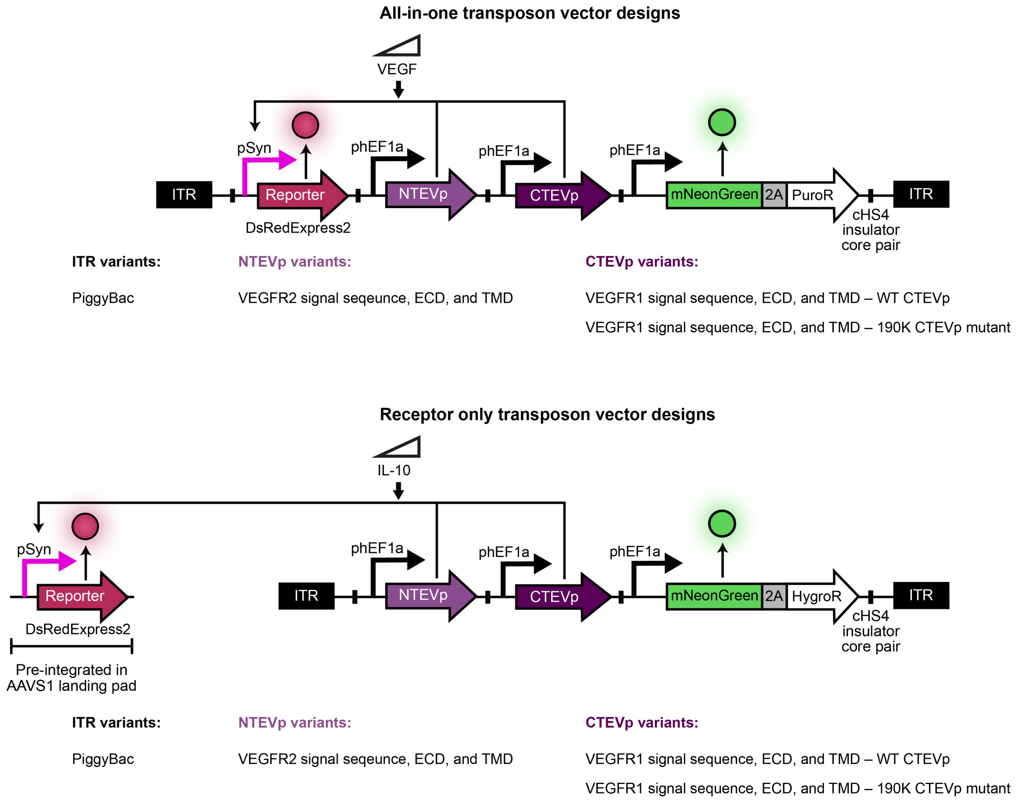


**Supplementary Figure 6. Supplementary information for the conversion of VEGFR.**
**(g)** Schematics detailing the composition of the transposon vectors used to generate stable cell lines expressing VEGF NatE MESA receptors. For the “all-in-one” vector, the reporter, two receptor chains, and selection marker were all included in the same transposon vector and used to transduce HEK293FT cells. For the “reporter-less” vector, the reporter was excluded, and the vector was used to transduce the previously engineered HEK293FT landing pad reporter cells. In both vector designs, transcriptional units (containing the promoter, the gene, and the terminator) are flanked by a pair of cHS4 insulators. Abbreviations: VEGFR, vascular endothelial growth factor receptor; ECD, ectodomain; TMD, transmembrane domain; TEVp, Tobacco Etch Virus protease; NTEVp, N-terminal component of split, mutant TEVp; CTEVp, C-terminal component of split, mutant TEVp; ITR, inverted terminal repeat; 2A, P2A peptide; PuroR, puromycin resistance gene; HygroR, hygromycin resistance gene; cHS4, chicken hypersensitive site 4 insulator; hEF1a, human elongation factor alpha.

h

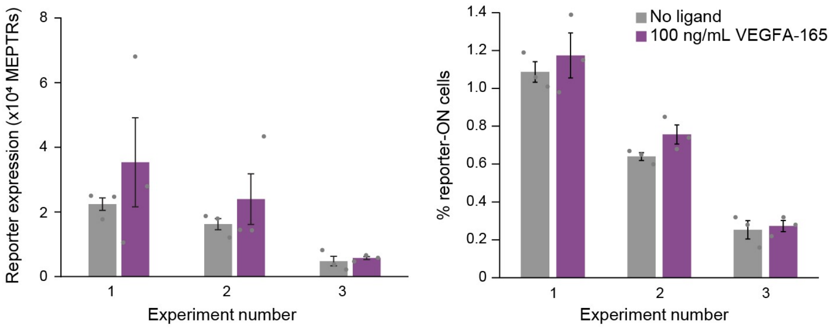


**Supplementary Figure 6. Supplementary information for the conversion of VEGFR.**
**(h)** Post-hoc analysis of silencing effects on all-in-one transposon cell lines. Reporter expression as well as percent of cells with an active reporter (defined by non-engineered HEK293FTs) were compared across three different experiments (with 1 being the earliest and 3 being the latest), with and without exogenous VEGF. Reporter signal and percent of cells with an active reporter decrease over experiment replicates (multi-factor ANOVA, $p < 0.001$). No statistical significance was found with a Welch's t-test between the no ligand and VEGFA treatments for reporter expression and percent of cells with an active reporter for each experiment. Each bar represents the mean across three biologic replicates and error bars indicate standard error of the mean (S.E.M). The receptor pair used in this panel is a NTEVp receptor chain with VEGFR2 signal sequence, ECD and TMD, WT NTEVp and a CTEVp receptor chain with VEGFR1 signal sequence, ECD and TMD, 190K CTEVp. Abbreviations: VEGFA, vascular endothelial growth factor A; MEPTRs, molecules of equivalent PE-TexasRed.

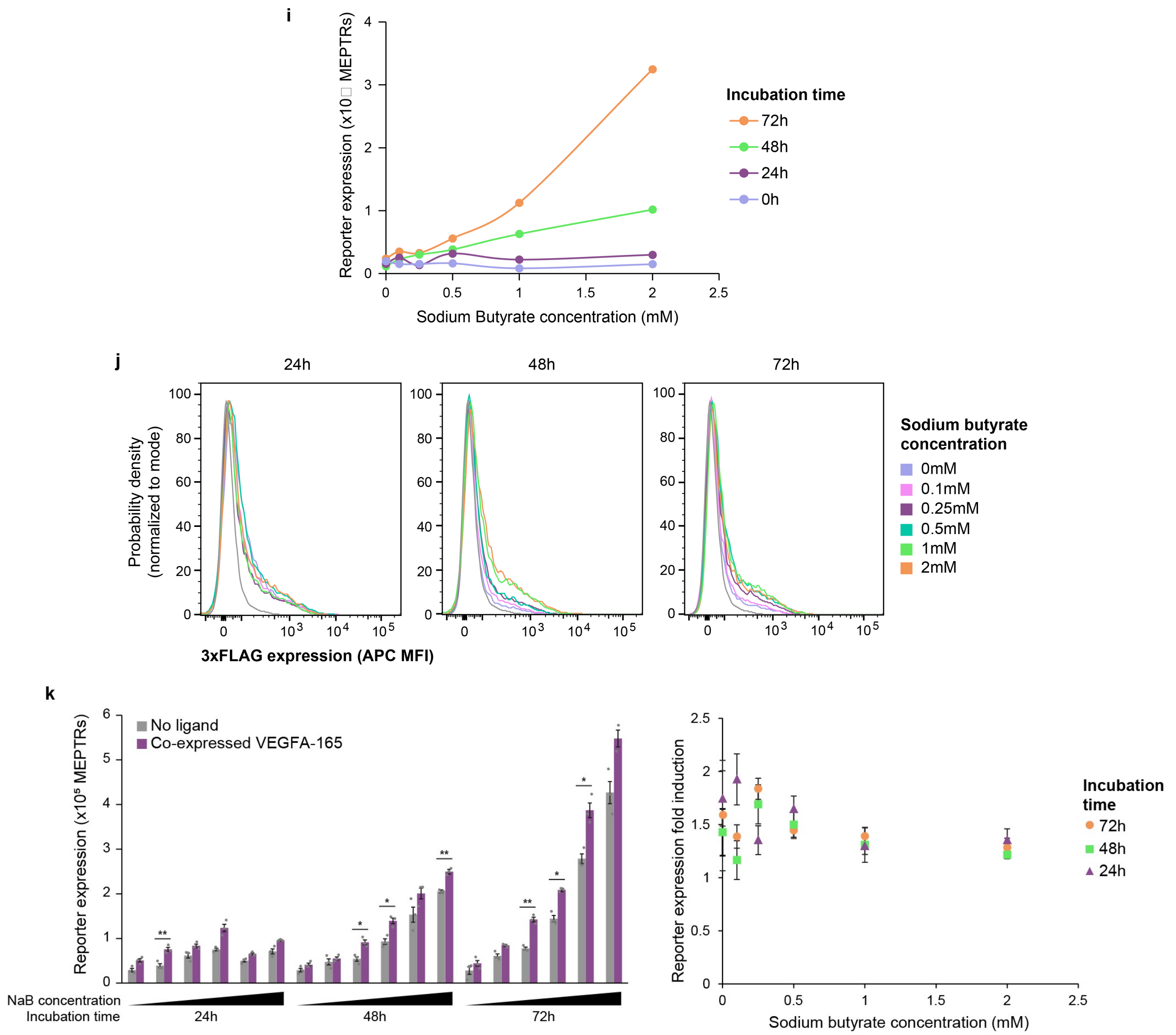


**Supplementary Figure 6. Supplementary information for the conversion of VEGFR.**
**(i)** Sodium butyrate (NaB) was employed to reverse silencing to confirm the cause of a reduction in signaling inducibility across experiments. Effect of NaB concentration and incubation time on VEGF-independent reporter expression in the "all-in-one" VEGF NatE MESA cell line. Each dot represents a single data point (replicate). **(j)** Effect of NaB concentration and incubation time on receptor surface expression as measured via 3xFLAG epitope tag. Note that this is total receptor expression—NTEVp and CTEVp chain expression cannot be differentiated in this experiment because they contain the same 3xFLAG epitope tag. **(k)** Effect of NaB concentration and incubation time on reporter expression in the presence and absence of transfected VEGF. Fold inductions (ratio of reporter expression in the presence and absence of VEGF) were plotted as a function of NaB concentration and incubation time. Reporter expression is significantly impacted by NaB dose, NaB incubation time, and the interaction between those variables (multi factor ANOVA, $p < 0.001$). Results for two-tailed Welch's t-test are indicated above corresponding bars for significant pairs only (* $p < 0.05$, ** $p < 0.01$). Each bar represents the mean across three biologic replicates and error bars indicate standard error of the mean (S.E.M). The receptor pair used in these panels is a NTEVp receptor chain with VEGFR2 signal sequence, ECD and TMD, WT NTEVp and a CTEVp receptor chain with VEGFR1 signal sequence, ECD and TMD, 190K CTEVp. Abbreviations: MEPTRs, molecules of equivalent PE-TexasRed; APC FI, allophycocyanin fluorescence intensity; NaB, sodium butyrate.

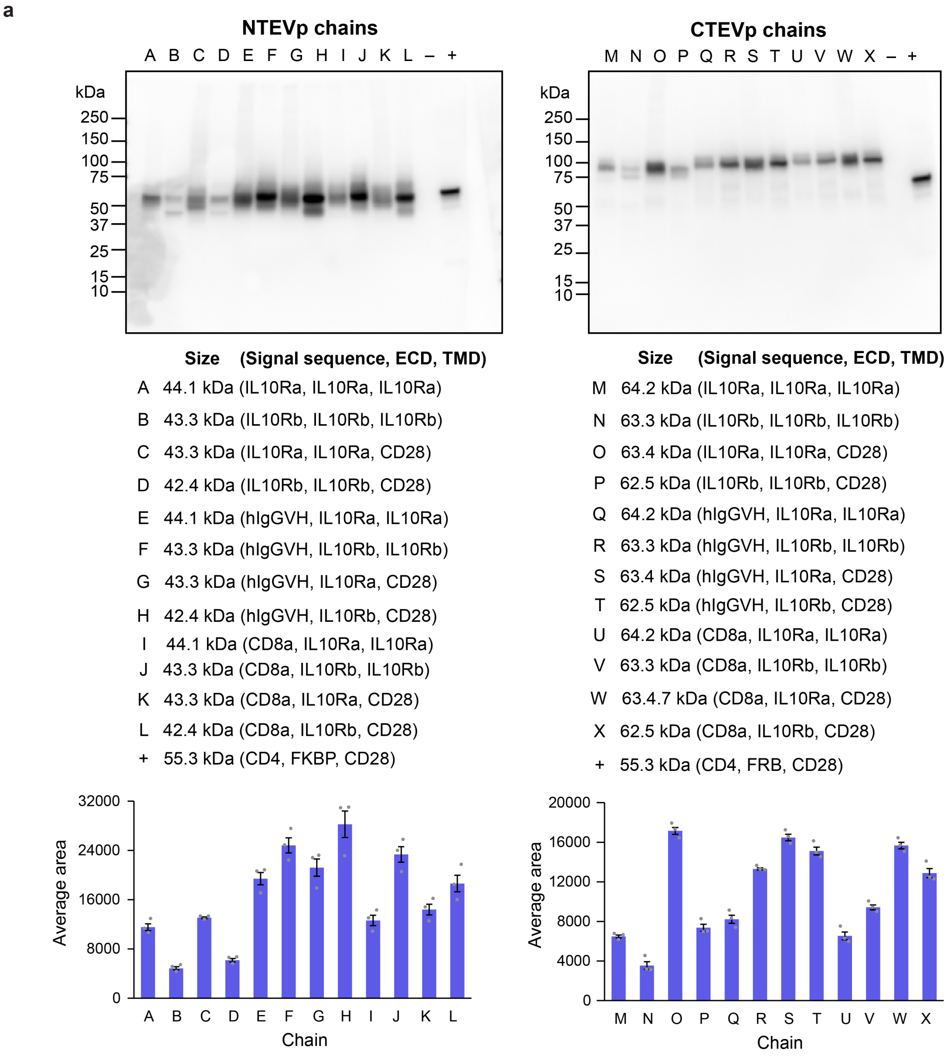


**Supplementary Figure 7. Supplementary information for the conversion of IL-10R.**
**(a)** Whole-cell expression of each IL-10 NatE MESA receptor chain was measured via western blotting of a 3xFLAG epitope tag fused to the N-terminus of receptors. Soluble mNeonGreen (mNG) or mTagBFP2 (mTB2) were included as negative controls (–) because each receptor chain-expressing plasmid is a poly-transfection vector that also contains either a constitutively expressed mNG or mTB2 in the backbone. A rapamycin-sensing MESA receptor, pPD810 was included as an internal control for 3xFLAG-tagged protein expression (+)[2]. Quantification of expression levels from bands in western blot is shown below each gel image. To quantify band intensity, western blot images were imported into ImageJ and analyzed using the analyze gel feature. Three images with different exposure times and no saturated pixels were used for analysis. Intensities of all bands (area under the curve) in each lane excluding the band representing the pPD810 control were summed and averaged across three exposure times. Error bars depict the standard error of the mean (S.E.M.) for these three measurements. Abbreviations: IL10R, interleukin-10 receptor; ECD, ectodomain; TMD, transmembrane domain; TEVp, Tobacco Etch Virus protease; NTEVp, N-terminal component of split, mutant TEVp; CTEVp, C-terminal component of split, mutant TEVp.

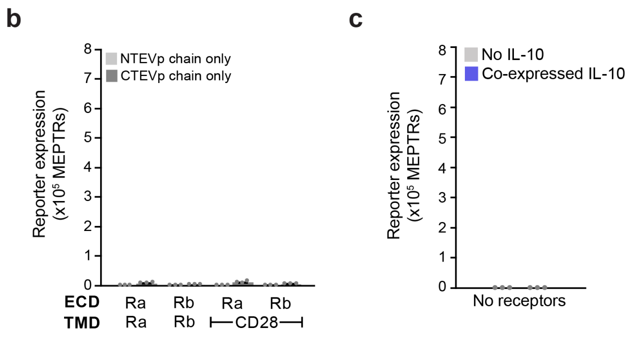


**Supplementary Figure 7. Supplementary information for the conversion of IL-10R.**
**(b)** The reporter cell line was transfected with plasmids encoding each receptor chain individually to assess single receptor chain-induced reporter expression in the absence of ligand. No reporter expression is observed with NTEVp chains, which do not contain a synTF. Minimal reporter expression is observed with CTEVp chains (e.g., from spontaneous release of the synTF from the membrane). The y-axis is scaled to match that of **Figure 2c. (c)** The reporter cell line was transfected to express IL-10 only to show that no reporter expression is induced only in the presence of IL-10. Each bar represents the mean of transfected cells across three biologic replicates and error bars indicate standard error of the mean (S.E.M). Abbreviations: IL-10R, interleukin-10 receptor; ECD, ectodomain; TMD, transmembrane domain; TEVp, Tobacco Etch Virus protease; NTEVp, N-terminal component of split, mutant TEVp; CTEVp, C-terminal component of split, mutant TEVp; Ra, IL-10Ra; Rb, IL-10Rb; MEPTRs, molecules of equivalent PE-TexasRed.

d

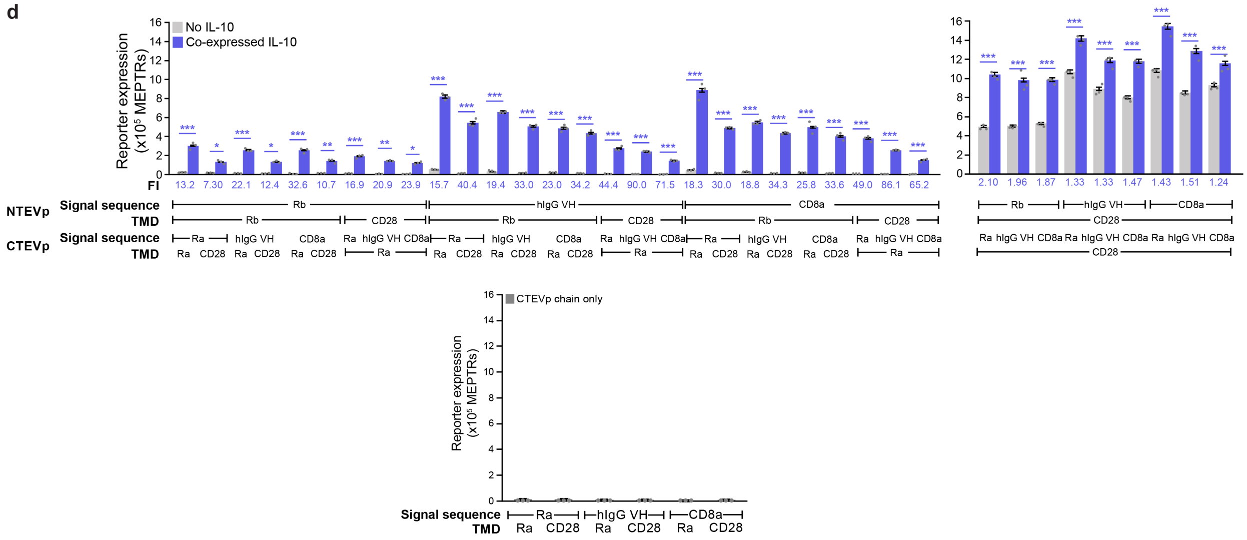


**Supplementary Figure 7. Supplementary information for the conversion of IL-10R.**
**(d)** Evaluation of the effect of NatE MESA receptor signal sequence on reporter expression with receptors expressed by transfection and co-expressed IL-10. One inducible ECD configuration from the initial panel was selected (NTEVp-IL-10Rb and CTEVp-IL-10Ra), and pairwise combinations of receptor chains with varying signal sequences and TMDs were evaluated in the presence and absence of co-expressed IL-10. Fold induction (FI) values are indicated below each pair of bars for each receptor chain combination, indicating the ratio between reporter expression in the presence of and absence of ligand. All receptor configurations were inducible across all signal sequence and transmembrane domain pairings (multi factor ANOVA, * $p < 0.05$, ** $p < 0.01$, *** $p < 0.001$). Signal sequence choice impacted both background and induced expression (multi factor ANOVA, $p < 0.05$). The use of a CD28-based transmembrane domain on both receptor chains in a pair leads to substantially higher background signal. The bottom plot shows that expression of each single CTEVp chain alone drives minimal reporter expression. Each bar represents the mean of transfected cells across three biologic replicates and error bars indicate standard error of the mean (S.E.M). Abbreviations: IL-10R, interleukin-10 receptor; ECD, ectodomain; TMD, transmembrane domain; TEVp, Tobacco Etch Virus protease; NTEVp, N-terminal component of split, mutant TEVp; CTEVp, C-terminal component of split, mutant TEVp; Ra, IL-10Ra; Rb, IL-10Rb; MEPTRs, molecules of equivalent PE-TexasRed.

e

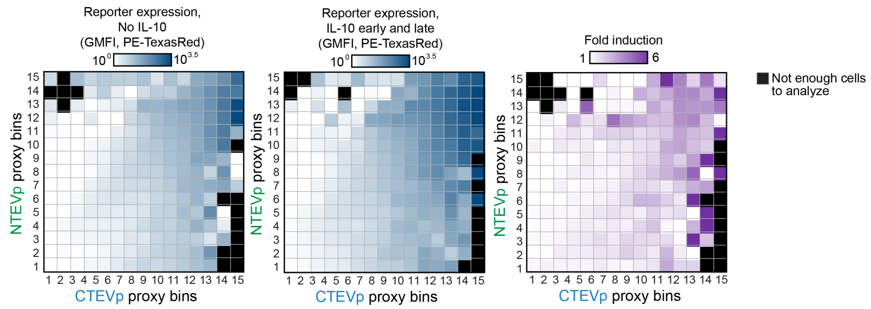


**Supplementary Figure 7. Supplementary information for the conversion of IL-10R.**
**(e)** Post-hoc analysis of cells poly-transfected to express IL-10 NatE MESA receptors at various expression levels and treated with recombinant IL-10 (250 ng/mL) at early (14 h post-transfection) and late (38 h post-transfection) time points (transfected HEK293FT cells). Expression of each single receptor chain was quantified via a proxy fluorescent protein included in the receptor-encoding plasmid under a separate constitutive promoter. In this panel, cells were transfected using the poly-transfection technique in which individual receptor-encoding plasmids are complexed with transfection reagents separately and applied to cells as separate transfection particles[4]. Cells were binned based on the expression level of each fluorescent proxy and both reporter output and fold induction (ratio of geometric mean of reporter expression in the presence and absence of IL-10) were calculated for each bin. Bins where the cell count was below 100 are blacked out and were not analyzed. Highest inducibility is observed with bins corresponding to cells that received the most plasmids encoding both receptor chains (upper right-hand corner). A visual description of this poly-transfection workflow is included in **Supplementary Figure 2**. These plots display data for one representative biological replicate of each treatment. The receptor pair used in this panel is a NTEVp receptor chain with CD8a signal sequence, IL-10Rb ECD and TMD, 75S NTEVp and a CTEVp receptor chain with IL-10Ra signal sequence, IL-10Ra ECD and TMD, 190K CTEVp. Abbreviations: IL-10R, interleukin-10 receptor; TEVp, Tobacco Etch Virus protease; NTEVp, N-terminal component of split, mutant TEVp; CTEVp, C-terminal component of split, mutant TEVp; GMFI, geometric mean of fluorescence intensity.

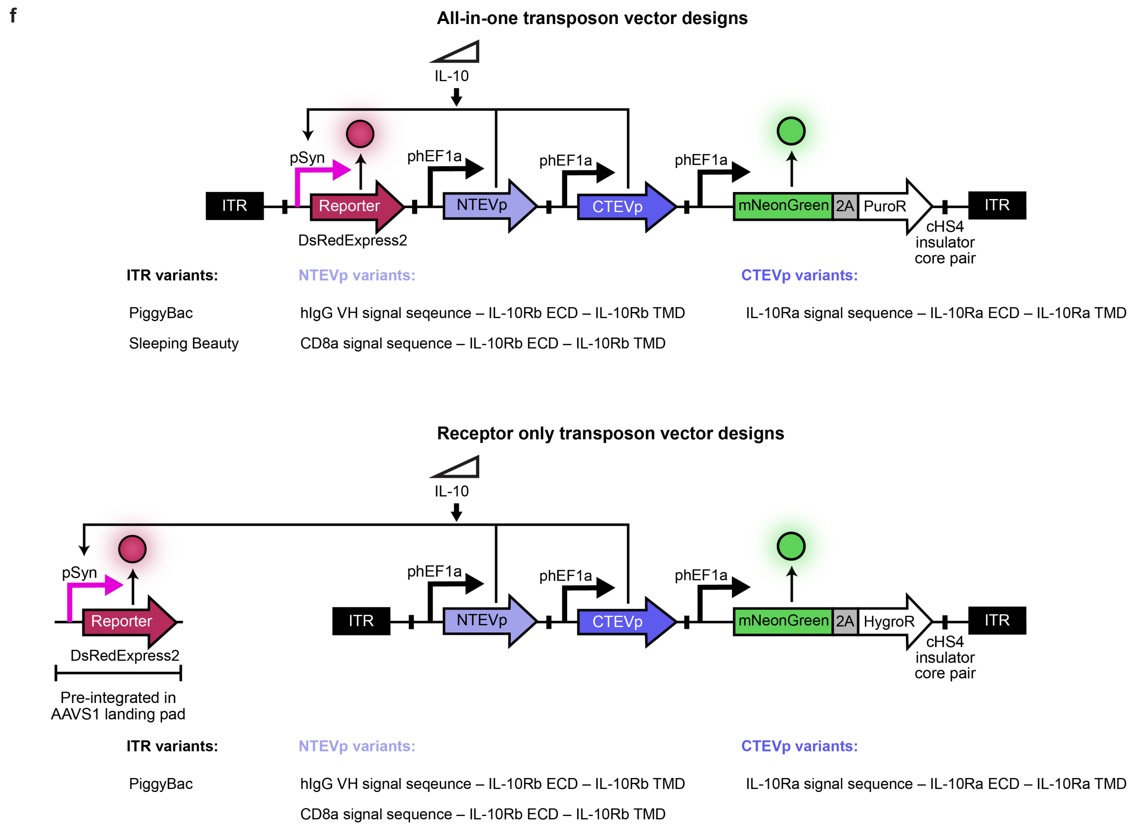


**Supplementary Figure 7. Supplementary information for the conversion of IL-10R.**
**(f)** Schematics detailing the composition of the transposon vectors used to generate stable cell lines expressing IL-10 NatE MESA receptors. For the "all-in-one" vector, the reporter, both receptors, and a selection marker were all included in the same transposon vector and used to transduce HEK293FT cells. For the "reporter-less" vector, the reporter was excluded, and the vector was used to transduce the previously engineered HEK293FT landing pad reporter cells. In both vector designs, transcriptional units (containing the promoter, the gene, and the terminator) are flanked by a pair of cHS4 insulators. Abbreviations: IL-10R, interleukin-10 receptor; ECD, ectodomain; TMD, transmembrane domain; TEVp, Tobacco Etch Virus protease; NTEVp, N-terminal component of split, mutant TEVp; CTEVp, C-terminal component of split, mutant TEVp; ITR, inverted terminal repeat; 2A, P2A peptide; PuroR, puromycin resistance gene; HygroR, hygromycin resistance gene; cHS4, chicken hypersensitive site 4 insulator; hEF1a, human elongation factor alpha.

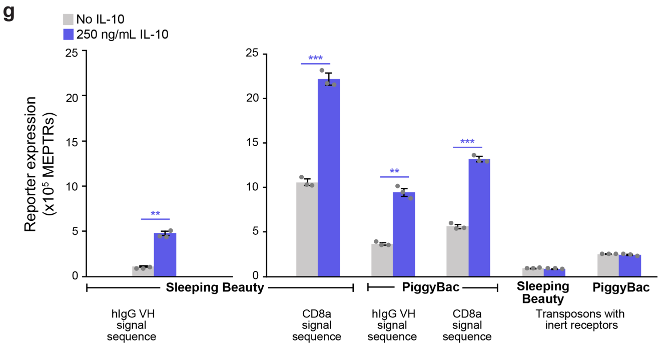


**Supplementary Figure 7. Supplementary information for the conversion of IL-10R.**
**(g)** Stable cell lines engineered with two IL-10 NatE MESA receptor chains (which differ in the signal sequence on the NTEVp receptor) and with two different transposon types were cultured with and without exogenous human IL-10 to evaluate ability to sense recombinant ligand. All transposon designs yield an IL-10-induced increase in reporter expression (two-tailed Welch’s t test, *** $p < 0.001$, ** $p < 0.01$). Two cell lines that contain receptors that do not sense IL-10 (inert receptors in this assay; receptors containing rapamycin-binding domains in place of IL-10R ECDs) were also included as negative controls to show that exogenous IL-10 ligand itself does not activate the reporter (two-tailed Welch’s t test, $p > 0.05$, not significant). Each bar represents the mean of mNeonGreen+ cells (constitutive transposon marker) across three biologic replicates and error bars indicate standard error of the mean (S.E.M). The receptor pairs used in these panels include a constant CTEVp receptor chain with IL-10Ra signal sequence, IL-10Ra ECD and TMD, 190K CTEVp and NTEVp receptor chains with varying signal sequences, IL-10Rb ECD and TMD, 75S NTEVp. Abbreviations: IL-10R, interleukin-10 receptor; MEPTRs, molecules of equivalent PE-TexasRed.

h

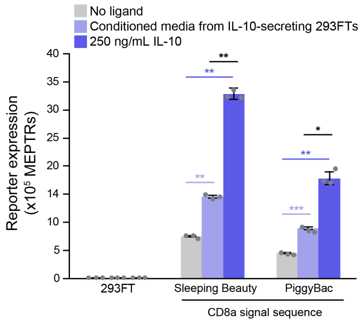


**Supplementary Figure 7. Supplementary information for the conversion of IL-10R.**
**(h)** Stable cell lines engineered with two IL-10 NatE MESA receptor pairs (which differ in the signal sequence on the NTEVp receptor) via a Piggy Bac transposon can detect IL-10 in conditioned media from HEK293FT cells engineered to secrete human IL-10 (two-tailed Welch's t test, *** $p < 0.001$, ** $p < 0.01$, * $p < 0.05$). Each bar represents the mean of mNeonGreen+ cells (constitutive transposon marker) across three biologic replicates and error bars indicate standard error of the mean (S.E.M). The receptor pair used in this panel is a NTEVp receptor chain with CD8a signal sequence, IL-10Rb ECD and TMD, 75S NTEVp and a CTEVp receptor chain with IL-10Ra signal sequence, IL-10Ra ECD and TMD, 190K CTEVp. Abbreviations: IL-10R, interleukin-10 receptor; MEPTRs, molecules of equivalent PE-TexasRed.

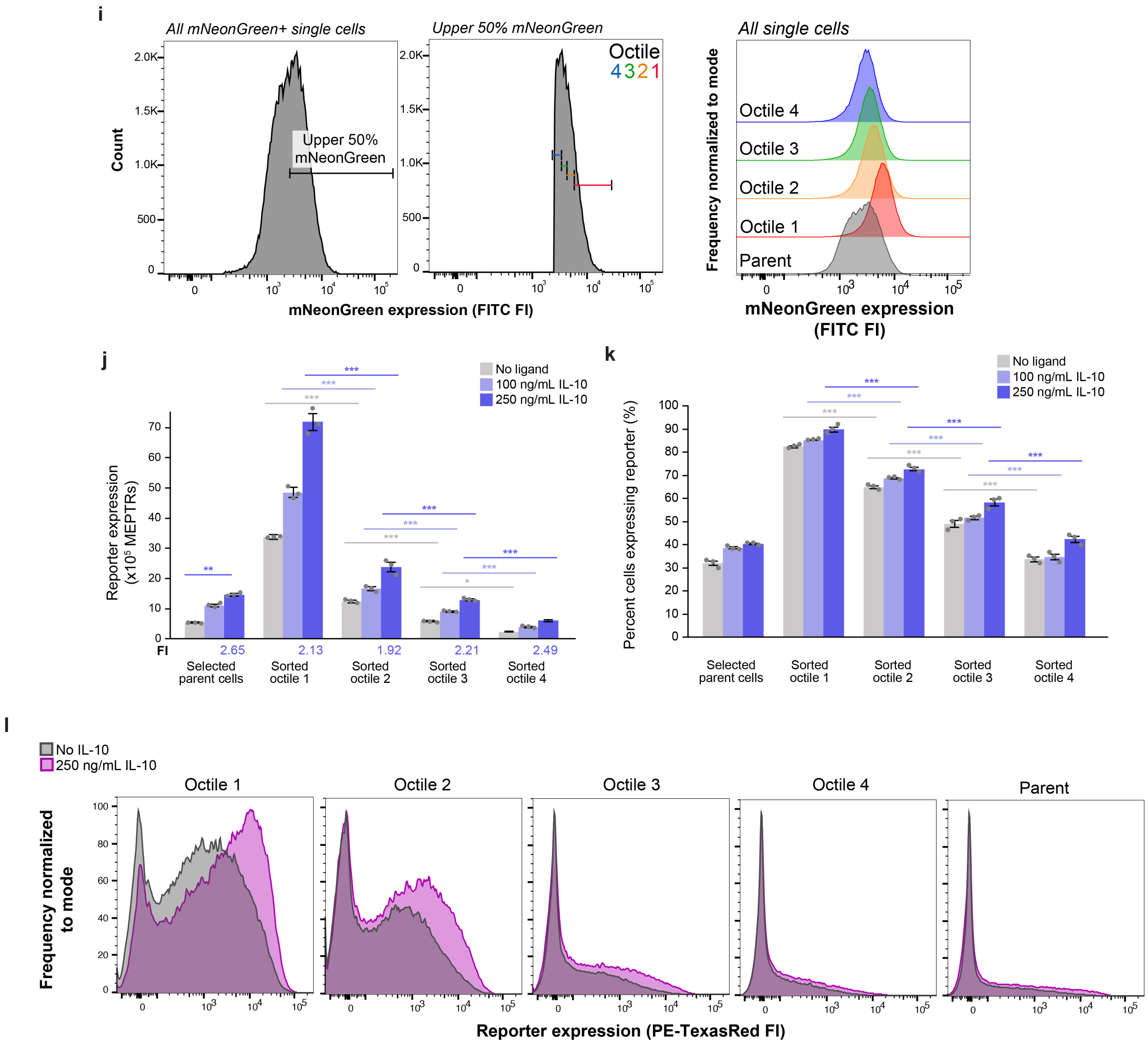


**Supplementary Figure 7. Supplementary information for the conversion of IL-10R.**
**(i)** A Fluorescence-Assisted Cell Sorting (FACS) strategy was devised to evaluate how different transposon copy numbers contribute to bulk population signaling with IL-10 NatE MESA receptors stably expressed in HEK293FTs. The distribution of mNeonGreen expression (constitutive transposon marker) was broken into eight octiles that each contained 12.5% of the total mNeonGreen+ cells, and the top 4 octiles were isolated. One week after sorting, the expanded octile populations retain differences in mNeonGreen expression. The four octiles were cultured with and without exogenous, recombinant IL-10 to evaluate **(j)** inducibility and **(k)** percent of cells expressing any reporter (defined by non-engineered HEK293FTs). Both reporter expression and percent of cells signaling decreases as octile number increases (mNeonGreen expression decreases), but fold induction is not substantially changed (multi-factor ANOVA, * $p < 0.05$, *** $p < 0.001$). Each bar represents the mean of mNeonGreen+ cells (constitutive transposon marker) across three biologic replicates and error bars indicate standard error of the mean (S.E.M). **(l)** The distribution of reporter expression across each expanded octile population suggests that in general, cells that dimly express reporter in the absence of ligand express more reporter when the ligand is present. This is evidenced by a growth in the shoulder of the histograms as opposed to a full population shift. The receptor pair used in these panels is a NTEVp receptor chain with CD8a signal sequence, IL-10Rb ECD and TMD, 75S NTEVp and a CTEVp receptor chain with IL-10Ra signal sequence, IL-10Ra ECD and TMD, 190K CTEVp. Abbreviations: IL-10R, interleukin-10 receptor; MEPTRs, molecules of equivalent PE-TexasRed; FITC or PE-TexasRed FI, FITC or PE-TexasRed fluorescence intensity; FI, fold induction.

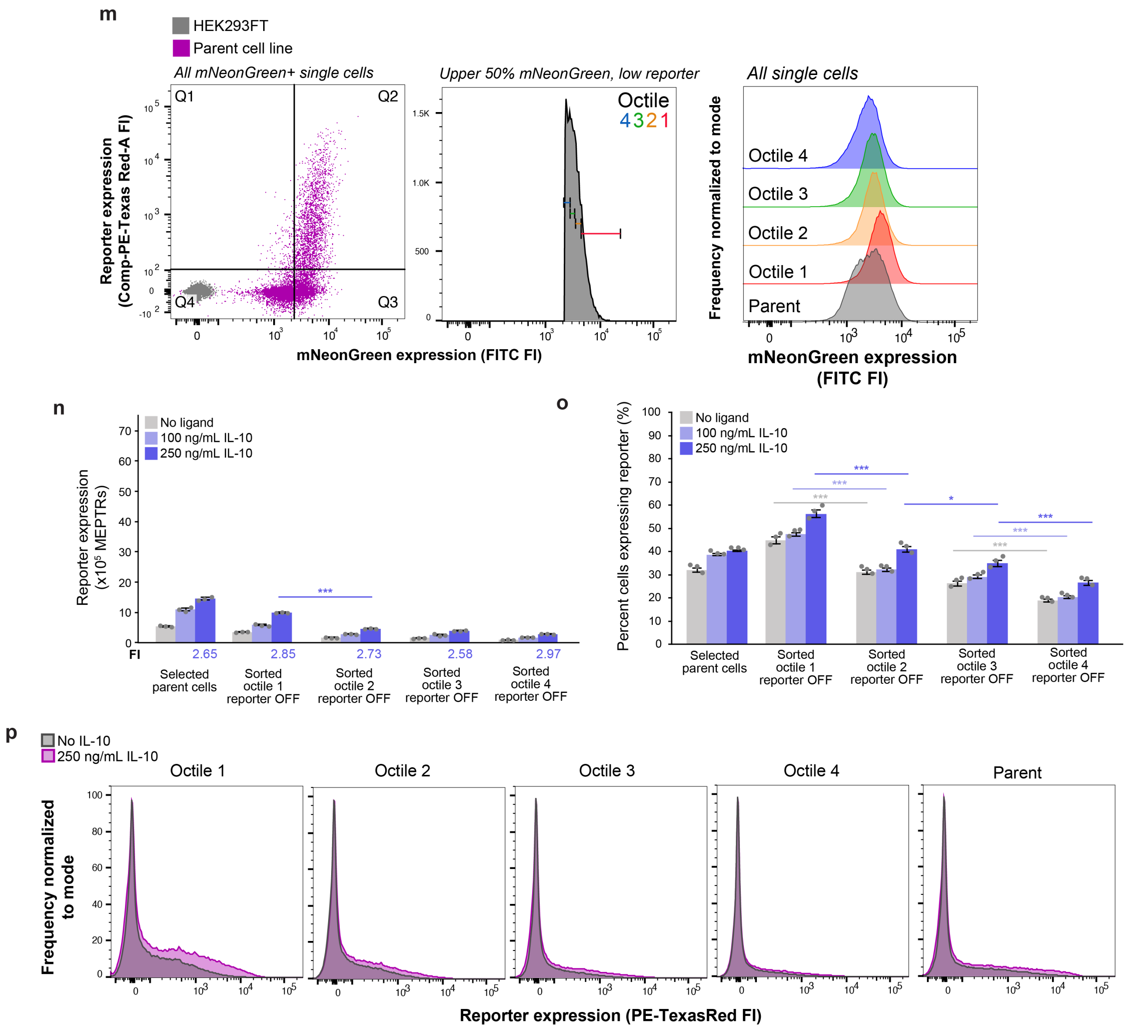


**Supplementary Figure 7. Supplementary information for the conversion of IL-10R.**
**(m)** A FACS strategy was devised to evaluate i) how different transposon copy numbers contribute to bulk population signaling and ii) whether a low-background population could be isolated for IL-10 NatE MESA receptors stably expressed in HEK293FTs. First, a quadrant gate was applied to identify low reporter expressing cells (cells were not treated with ligand before this sort) and the upper half of mNeonGreen expression (constitutive transposon marker). From here, mNeonGreen expression was broken into quartiles that each contained 12.5% of the total mNeonGreen+ cells (25% of the top half of mNeonGreen+ cells) and isolated. One week after sorting, the expanded octile populations retain differences in mNeonGreen expression. The four octiles were cultured with and without exogenous, recombinant IL-10 to evaluate **(n)** inducibility and **(o)** percent of cells expressing any reporter (defined by non-engineered HEK293FTs). The percent of cells signaling when treated with ligand decreases as octile number increases (mNeonGreen expression decreases), reporter expression decreases between octiles 1 and 2, and fold inductions are not substantially changed (multi-factor ANOVA, * $p < 0.05$, *** $p < 0.001$). Compared to sorted populations in **Supplementary Figure 7 i-l,** overall signal and percent of signaling cells are much lower. Each bar represents the mean of mNeonGreen+ cells (constitutive transposon marker) across three biologic replicates and error bars indicate standard error of the mean (S.E.M). **(p)** The distribution of reporter expression across each expanded octile population shows that in general, cells that are dimly expressing reporter in the absence of ligand express more reporter when the ligand is present. This is evidenced by a growth in the shoulder of the histograms as opposed to a full population shift. The receptor pair used in these panels is a NTEVp receptor chain with CD8a signal sequence, IL-10Rb ECD and TMD, 75S NTEVp

and a CTEVp receptor chain with IL-10Ra signal sequence, IL-10Ra ECD and TMD, 190K CTEVp. Abbreviations: IL-10R, interleukin-10 receptor; MEPTRs, molecules of equivalent PE-TexasRed; FITC or PE-TexasRed FI, FITC or PE-TexasRed fluorescence intensity; FI, fold induction.

q

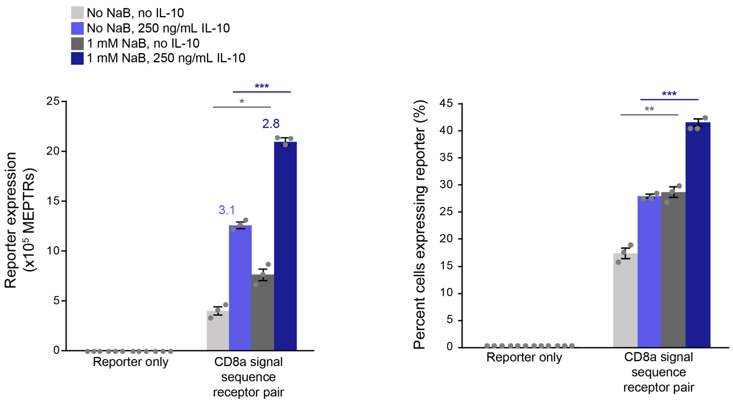


**Supplementary Figure 7. Supplementary information for the conversion of IL-10R.**
**(q)** To evaluate whether reversal of silencing could increase the proportion of cells that can signal, stable transposon-engineered cells with reporter only or with reporter and IL-10 NatE MESA receptors were treated with 1 mM NaB for 48 h before being treated with or without exogenous, recombinant IL-10. Background signal, induced signal, and percent of cells expressing reporter (defined by non-engineered HEK293FTs) increased with NaB treatment (two-tailed Welch's t-test, * $p < 0.05$, ** $p < 0.01$, *** $p < 0.001$). The receptor pair used in this panel is a NTEVp receptor chain with CD8a signal sequence, IL-10Rb ECD and TMD, 75S NTEVp and a CTEVp receptor chain with IL-10Ra signal sequence, IL-10Ra ECD and TMD, 190K CTEVp. Each bar represents the mean of mNeonGreen+ cells (constitutive transposon marker) across three biologic replicates and error bars indicate standard error of the mean (S.E.M). Abbreviations: IL-10R, interleukin-10 receptor; MEPTRs, molecules of equivalent PE-TexasRed.

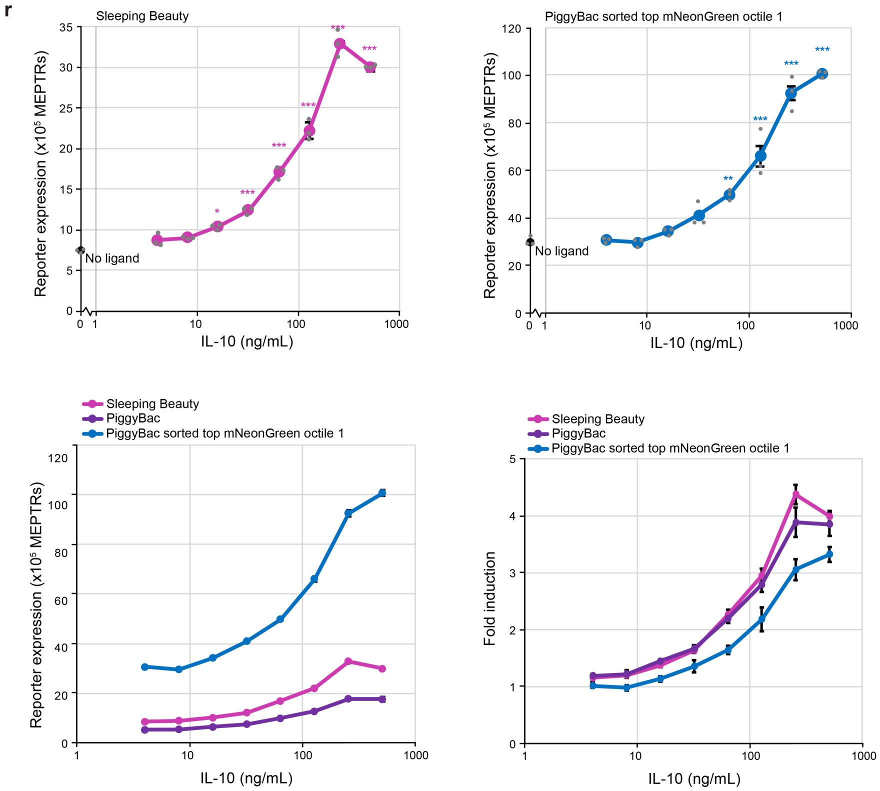


**Supplementary Figure 7. Supplementary information for the conversion of IL-10R.**
**(r)** Reporter expression dose response data for stable cells engineered with the reporter and the IL-10 NatE MESA receptor pair (NTEVp receptor chain with CD8a signal sequence, IL-10Rb ECD and TMD, 75S NTEVp; CTEVp receptor chain with IL-10Ra signal sequence, IL-10Ra ECD and TMD, 190K CTEVp)via a Sleeping Beauty transposon (antibiotic selected cells) or Piggy Bac transposon (antibiotic selected cells and top mNeonGreen octile). The Sleeping Beauty transposon cell line exhibits a ligand dose-dependent increase in reporter expression until 250 ng/mL, after which reporter expression decreases, likely due to ligand toxicity. Ligand-induced reporter expression is significantly different from the untreated condition down to 16 ng/mL (single factor ANOVA, * $p < 0.05$, *** $p < 0.001$). The top octile Piggy Bac transposon cell line demonstrates a dose-dependent increase in reporter expression through 500 ng/mL. Ligand-induced reporter expression is significantly different from the untreated condition down to 64 ng/mL (single factor ANOVA, ** $p < 0.01$, *** $p < 0.001$). Though signal magnitude changes across these cell lines, fold induction (ratio of induced to background signal) is relatively conserved. Each point represents the mean of mNeonGreen+ cells (constitutive transposon marker) across three biologic replicates and error bars indicate standard error of the mean (S.E.M). Abbreviations: IL-10R, interleukin-10 receptor; MEPTRs, molecules of equivalent PE-TexasRed.

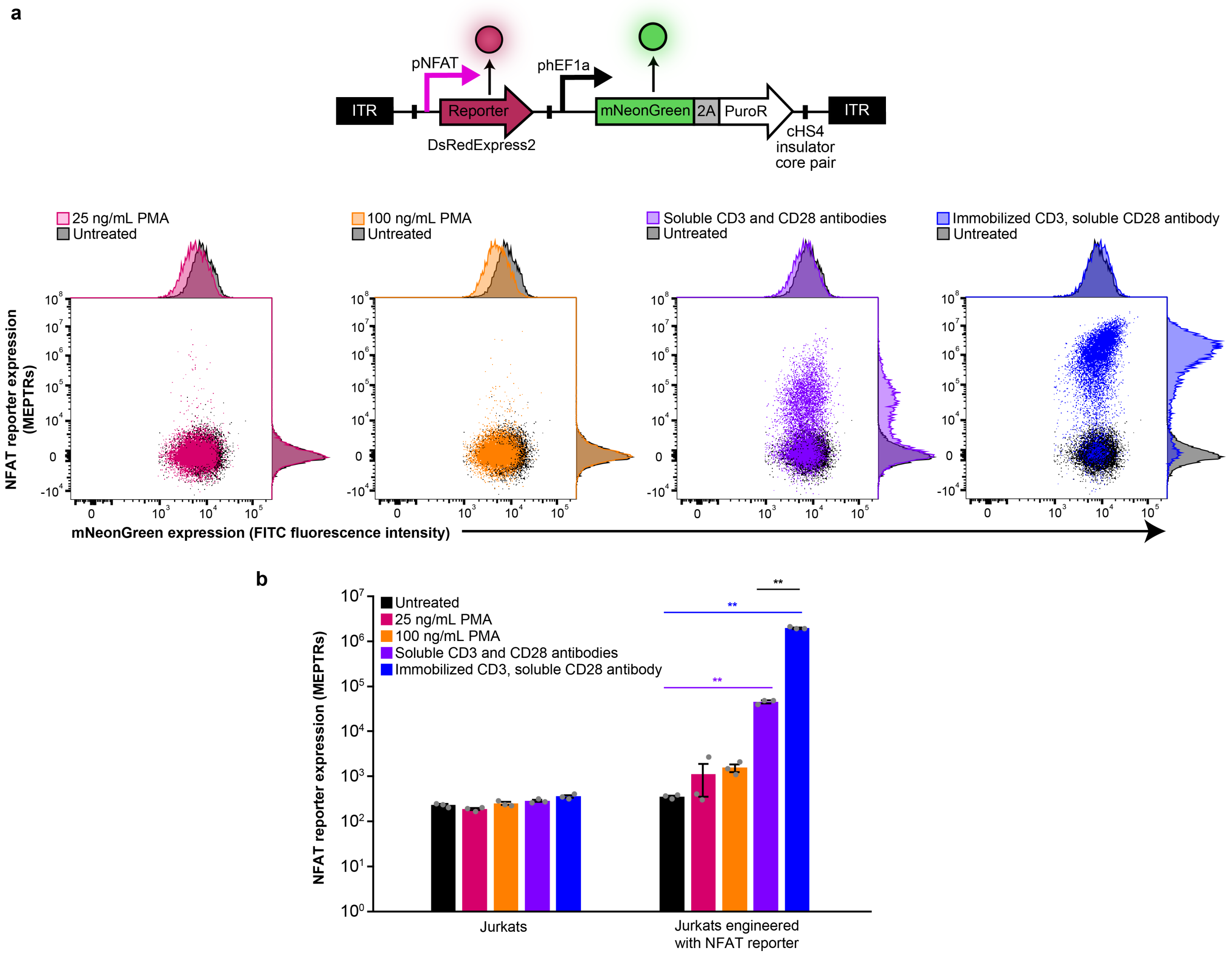


**Supplementary Figure 8. Supplementary information for NatE-MESA receptors can be used to construct novel, therapeutically motivated cellular functions.**
**(a)** Evaluation of engineered NFAT reporter activation. Jurkats were stably engineered with the PiggyBac transposon vector shown at the top containing an engineered NFAT-inducible promoter (6 NFAT response elements upstream of a YB_TATA minimal promoter) that drives expression of DsRedExpress2 as well as a constitutive promoter driving expression of mNeonGreen and a puromycin resistance gene. The NFAT reporter used in this construct was used in experiments shown in **Figure 3**. Cells were selected with puromycin and sorted for the top 12.5% of mNeonGreen expressing cells. To evaluate activation, engineered cells were cultured for 48 h with 25 ng/mL PMA, 100 ng/mL PMA, soluble anti-CD3 and anti-CD28 antibodies, or immobilized anti-CD3 and soluble anti-CD28 antibodies before assay. A representative sample for each treatment is shown. **(b)** NFAT reporter expression in parent Jurkats and engineered cells with each treatment were quantified. Both concentrations of PMA did not induce a statistically significant increase in reporter expression but both antibody treatments did activate reporter expression (two-tailed Welch's t test, * $p < 0.05$, ** $p < 0.01$). Culturing cells with an immobilized anti-CD3 antibody and a soluble anti-CD28 antibody yielded the strongest activation. Bars represents the mean of single cells (for Jurkats) or single mNeonGreen+ cells (for engineered cells) across three biologic replicates and error bars indicate standard error of the mean (S.E.M). Abbreviations: ITR, inverted terminal repeat; cHS4, chicken hypersensitive site 4 insulator; NFAT, nuclear factor of activated T cells; 2A, P2A peptide; PuroR, puromycin resistance gene; PMA, phorbol 12-myristate 13-acetate; MEPTRs, molecules of equivalent PE-TexasRed; hEF1a, human elongation factor alpha.

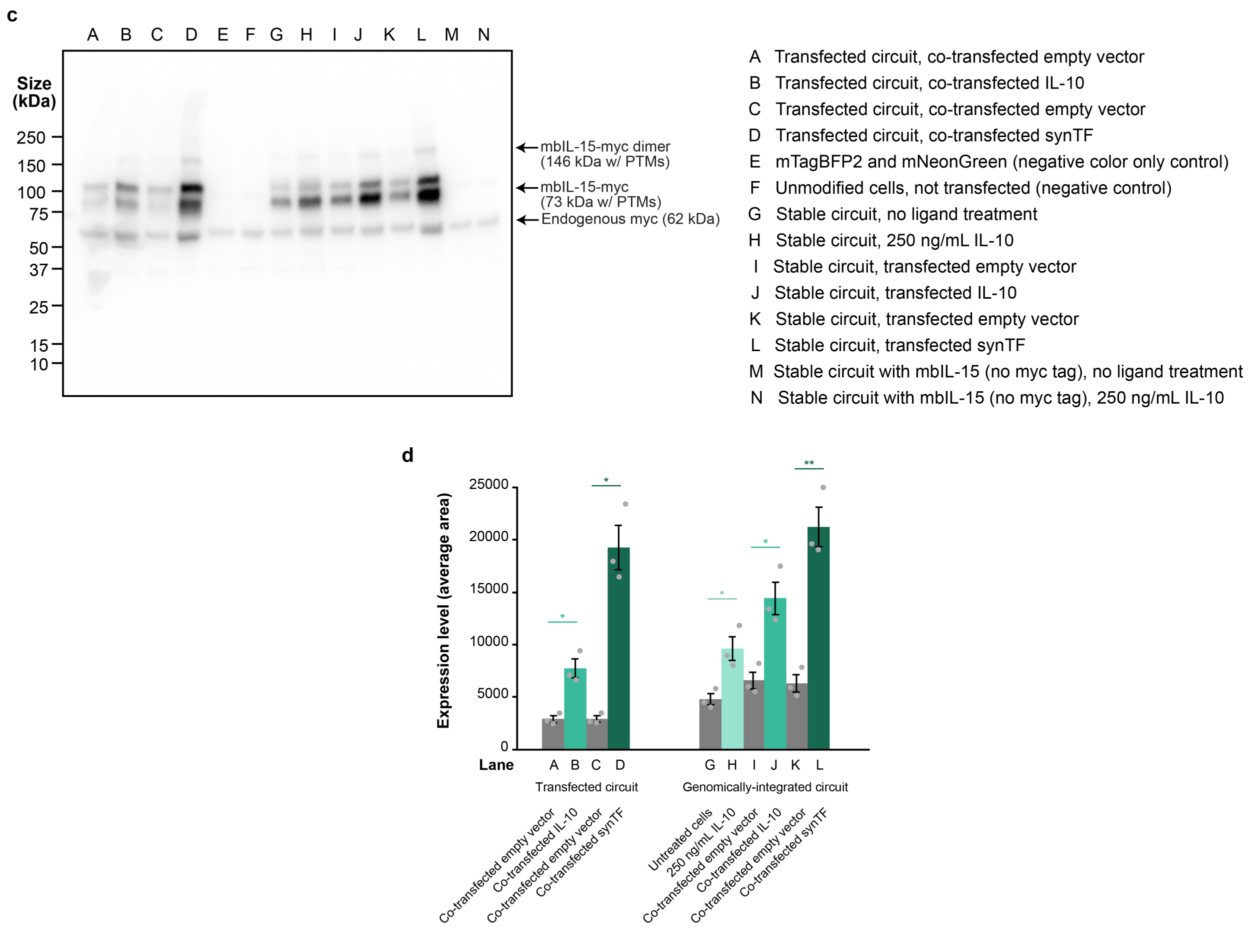


**Supplementary Figure 8. Supplementary information for NatE-MESA receptors can be used to construct novel, therapeutically motivated cellular functions.**
**(c)** Western blot showing expression of membrane-bound IL-15 (mbIL-15) in HEK293FTs. Sample descriptions for each lane are on the right. Arrow markers for relevant protein sizes are also shown. The expected dominant band size for mbIL-15 is 73 kDa including post-translational modifications (PTMs)[5]. A C-terminal myc epitope tag was included on the mbIL-15 for detection. A consistent band for endogenous myc protein expression appears in all lanes. Lanes A-D show that co-expressed IL-10 and syn-TF lead to an increase in mbIL-15 expression when the biosensor circuit is transfected via a plasmid. Lanes G-L show that treatment with recombinant IL-10 or co-transfection with IL-10 and syn-TF in cells with stably integrated biosensor circuits yield an increase in mbIL-15 expression. Lanes E, F, M, N are negative controls. **(d)** Quantification of expression levels from bands in western blot shown in **(c)**. To quantify band intensity, western blot images were imported into ImageJ and analyzed using the analyze gel feature. Three images with different exposure times and no saturated pixels were used for analysis. Intensities of all bands (area under the curve) in each lane excluding the band representing the 62 kDa endogenous myc protein were summed and averaged across three exposure times. Error bars depict the standard error of the mean for these three measurements. Transfection with a plasmid encoding IL-10, transfection with a plasmid encoding a synTF, or treatment with 250 ng/mL recombinant IL-10 led to significant increases in mbIL-15 expression over conditions transfected with empty vector or untreated (two-tailed Welch's t test, * $p < 0.05$, ** $p < 0.01$). The receptor pair used in these panels is a NTEVp receptor chain with CD8a signal sequence, IL-10Rb ECD and TMD, 75S NTEVp and a CTEVp receptor chain with IL-10Ra signal sequence, IL-10Ra ECD and TMD, 190K CTEVp. Abbreviations: IL-10, interleukin-10; PTM, post-translational modification; mbIL-15, membrane-bound IL-15; synTF, synthetic transcription factor.

e

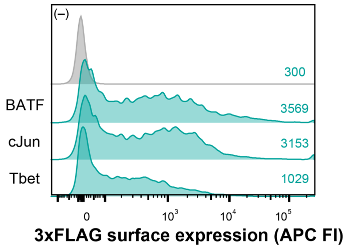


f

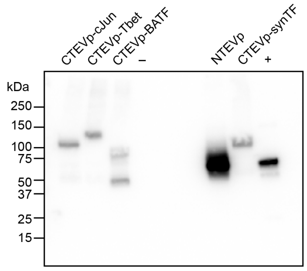


**Supplementary Figure 8. Supplementary information for NatE-MESA receptors can be used to construct novel, therapeutically motivated cellular functions.**
**(e)** Surface expression of IL-10 NatE MESA CTEVp receptors with different natural transcription factors (BATF, cJun, and Tbet) was measured via immunohistochemistry to detect a surface expressed 3xFLAG epitope tag on the N-terminus of each receptor. Histograms include all transfected cells based on expression of a constitutive transfection control fluorescent protein on a separate co-transfected plasmid, and the negative control is a sample transfected with just the transfection control plasmid. Mean APC fluorescence intensity for each sample is listed. **(f)** Whole-cell expression of each IL-10 NatE MESA receptor chain was measured via western blotting of the 3xFLAG epitope tag fused to the N-terminus of receptors. With each receptor chain , a plasmid encoding mTagBFP2 was included as a transfection control, so soluble mTagBFP2 was included alone as a negative control (–). A rapamycin-sensing MESA receptor, pPD810 was included as an internal control for 3xFLAG-tagged protein expression (+)[2]. Natural transcription factor-containing receptor chains produced a band at the expected size and the BATF-containing receptor produced a prominent, small cleavage product. The receptor pair used in these panels is a NTEVp receptor chain with CD8a signal sequence, IL-10Rb ECD and TMD, 75S NTEVp and a CTEVp receptor chain with IL-10Ra signal sequence, IL-10Ra ECD and TMD, 190K CTEVp. Abbreviations: BATF, basic leucine zipper transcription factor ATF-like; Tbet, T-box expressed in T cells; APC FI, allophycocyanin fluorescence intensity; TEVp, Tobacco Etch Virus protease; NTEVp, N-terminal component of split, mutant TEVp; CTEVp, C-terminal component of split, mutant TEVp; synTF, synthetic transcription factor.

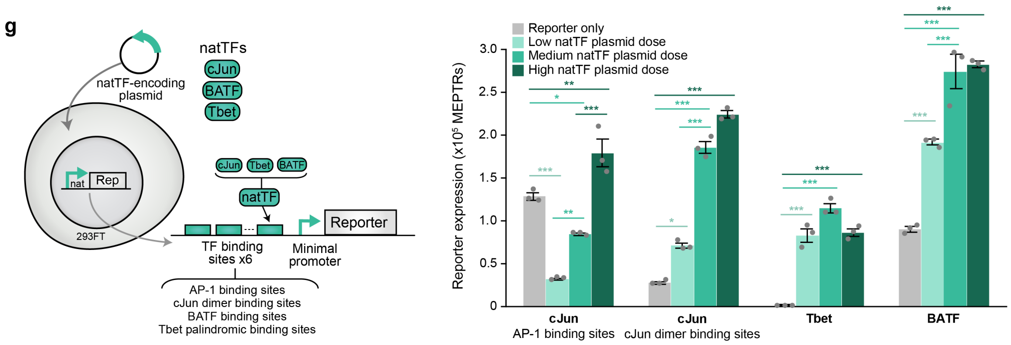


**Supplementary Figure 8. Supplementary information for NatE-MESA receptors can be used to construct novel, therapeutically motivated cellular functions**.

**(g)** Synthetic promoters were engineered to detect release of natural transcription factors. Six copies of binding sites for each transcription factor were placed upstream of a YB_TATA minimal promoter. These reporters were validated by co-transfecting three doses of transcription factor-encoding plasmid along with each reporter plasmid in HEK293FTs. Each reporter demonstrates a natTF plasmid-dependent increase in reporter expression (multi factor ANOVA, * $p < 0.05$, ** $p < 0.01$, *** $p < 0.001$). The cJun reporter with AP-1 binding sites shows a significant decrease in reporter expression at low and medium natTF plasmid doses compared to the reporter only condition. We hypothesize that this is because overexpressed cJun interacts with the AP-1 proteins (cJun and cFos) basally expressed in HEK293FTs, shifting the distribution of functional dimeric TFs to more cJun homodimers than cJun-cFos homodimers, which bind to the AP-1 response element more efficiently than cJun homodimers[6] The level of background reporter output from transfection of the reporter plasmid alone varies across designs likely because HEK293FTs express varying levels of cJun (and other AP-1 transcription factors) and BATF (and other ATF-like transcription factors) but not Tbet, endogenously. Each bar represents the mean of transfected cells across three biologic replicates and error bars indicate standard error of the mean (S.E.M). The receptor pair used in these panels is a NTEVp receptor chain with CD8a signal sequence, IL-10Rb ECD and TMD, 75S NTEVp and a CTEVp receptor chain with IL-10Ra signal sequence, IL-10Ra ECD and TMD, 190K CTEVp. Abbreviations: BATF, basic leucine zipper transcription factor ATF-like; Tbet, T-box expressed in T cells; MEPTRs, molecules of equivalent PE-TexasRed; Rep, reporter; TEVp, Tobacco Etch Virus protease; NTEVp, N-terminal component of split, mutant TEVp; CTEVp, C-terminal component of split, mutant TEVp; synTF, synthetic transcription factor; natTF, natural transcription factor; AP-1, activator protein-1.

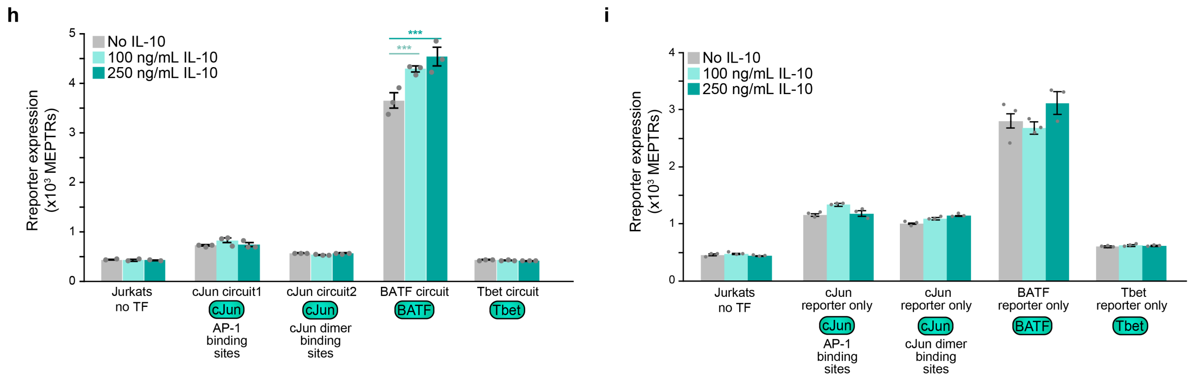


**Supplementary Figure 8. Supplementary information for NatE-MESA receptors can be used to construct novel, therapeutically motivated cellular functions.**
**(h)** Jurkats engineered with natural transcription factor containing IL-10 NatE MESA receptors were cultured with or without recombinant IL-10 for 48 h to assay synthetic reporter output from engineered natural transcription factor reporters. This is the earlier timepoint data that corresponds to data shown in **Figure 3f**. Ligand treatments within 48 h only produced a significant increase in reporter expression for the BATF circuit (multi-factor ANOVA, *** $p < 0.001$). **(i)** Jurkats engineered with just the synthetic reporters for each natural transcription factor were cultured with or without recombinant IL-10 for 48 h to assay how IL-10 alone (no receptors) impacts synthetic reporter output. Ligand treatments did not produce statistically significant changes in reporter expression. Each bar represents the mean of mNeonGreen+ cells (constitutive transposon marker) across three biologic replicates and error bars indicate standard error of the mean (S.E.M). The receptor pair used in these panels is a NTEVp receptor chain with CD8a signal sequence, IL-10Rb ECD and TMD, 75S NTEVp and a CTEVp receptor chain with IL-10Ra signal sequence, IL-10Ra ECD and TMD, 190K CTEVp. Abbreviations: BATF, basic leucine zipper transcription factor ATF-like; Tbet, T-box expressed in T cells; MEPTRs, molecules of equivalent PE-TexasRed; IL-10, interleukin-10; TF, transcription factor; AP-1, activator protein-1.

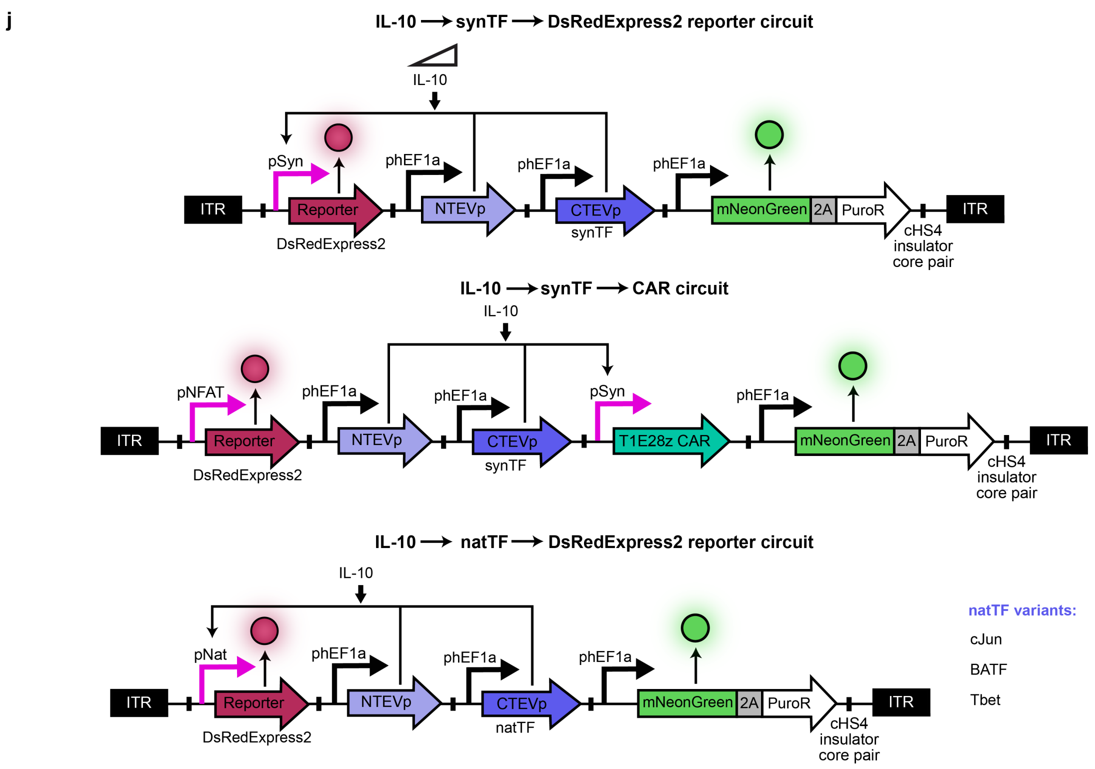


**Supplementary Figure 8. Supplementary information for NatE-MESA receptors can be used to construct novel, therapeutically motivated cellular functions.**
**(j)** Schematics detailing the composition of the Piggy Bac transposon vectors used to generate stable cell lines expressing IL-10 NatE MESA receptors in Jurkats and HEK293FTs for experiments related to **Figure 3**. In all vector designs, transcriptional units (containing the promoter, the gene, and the terminator) are flanked by a pair of cHS4 insulators. Abbreviations: ITR, inverted terminal repeat; cHS4, chicken hypersensitive site 4 insulator; NFAT, nuclear factor of activated T cells; 2A, P2A peptide; PuroR, puromycin resistance gene; TEVp, Tobacco Etch Virus protease; NTEVp, N-terminal component of split, mutant TEVp; CTEVp, C-terminal component of split, mutant TEVp; synTF, synthetic transcription factor; natTF, natural transcription factor; BATF, basic leucine zipper transcription factor ATF-like; Tbet, T-box expressed in T cells; IL-10, interleukin-10; hEF1a, human elongation factor alpha.

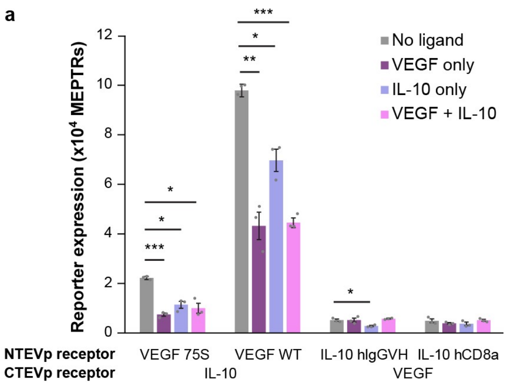


**Supplementary Figure 9. Supplementary information for multiplexing NatE MESA receptors to perform logical evaluation of multiple inputs.**

**(a)** Cross-reactivity of VEGF and IL-10 NatE MESA receptors. Indicated receptor chains were expressed by co-transfection of HEK293FT reporter cells along with co-expressed ligands or empty vector DNA. Cross reactivity was observed between VEGFR NTEVp and IL-10R CTEVp chains. Results for two-tailed Welch's t-test between different ligand conditions are indicated above corresponding bars for significant pairs only (* $p$ < 0.05, ** $p$ < 0.01, *** $p$ < 0.001). Each bar represents the mean of transfected cells across three biologic replicates and error bars indicate standard error of the mean (S.E.M). The VEGF receptor pair used in this panel is a NTEVp receptor chain with VEGFR2 signal sequence, ECD and TMD, WT/75S NTEVp and a CTEVp receptor chain with VEGFR1 signal sequence, ECD and TMD, 190K CTEVp. The IL-10 receptor pair used in this panel is a NTEVp receptor chain with CD8a/IgGVH signal sequence, IL-10Rb ECD and TMD, 75S NTEVp and a CTEVp receptor chain with IL-10Ra signal sequence, IL-10Ra ECD and TMD, 190K CTEVp. Abbreviations: VEGF, vascular endothelial growth factor; IL-10, interleukin-10; MEPTRs, molecules of equivalent PE-TexasRed; TEVp, Tobacco Etch Virus protease; NTEVp, N-terminal component of split, mutant TEVp; CTEVp, C-terminal component of split, mutant TEVp.

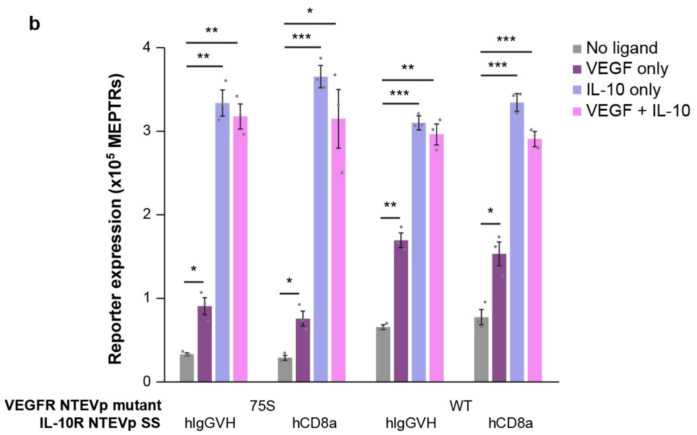


**Supplementary Figure 9. Supplementary information for multiplexing NatE MESA receptors to perform logical evaluation of multiple inputs.**

**(b)** Full panel of OR gate design choices. VEGF and IL-10 NatE MESA receptors were co-expressed in HEK293FTs along with the different co-expressed ligands (or empty vector filler DNA in the no ligand condition) via transfection. The main figure shows the combination of the VEGF receptor with NTEVp mutant 75S and the IL-10 receptor with NTEVp signal sequence hIgG VH (first combination on this plot). CTEVp chains were not varied between each of the receptor pairs. Choice of IL-10 receptor does not have an effect on OR gate performance (reporter expression with any ligand treatment), and choice of VEGF receptor does affect reporter expression with no ligand and VEGF-only (multi-factor ANOVA, $p < 0.05$). Results for two-tailed Welch's t-test between different ligand conditions are indicated above corresponding bars for significant pairs only (* $p < 0.05$, ** $p < 0.01$, *** $p < 0.001$). Each bar represents the mean of transfected cells across three biologic replicates and error bars indicate standard error of the mean (S.E.M). The VEGF receptor pair used in this panel is a NTEVp receptor chain with VEGFR2 signal sequence, ECD and TMD, WT/75S NTEVp and a CTEVp receptor chain with VEGFR1 signal sequence, ECD and TMD, 190K CTEVp. The IL-10 receptor pair used in this panel is a NTEVp receptor chain with CD8a/IgGVH signal sequence, IL-10Rb ECD and TMD, 75S NTEVp and a CTEVp receptor chain with IL-10Ra signal sequence, IL-10Ra ECD and TMD, 190K CTEVp. Abbreviations: VEGF, vascular endothelial growth factor; IL-10, interleukin-10; MEPTRs, molecules of equivalent PE-TexasRed; TEVp, Tobacco Etch Virus protease; NTEVp, N-terminal component of split, mutant TEVp; CTEVp, C-terminal component of split, mutant TEVp; WT, wildtype; SS, signal sequence.

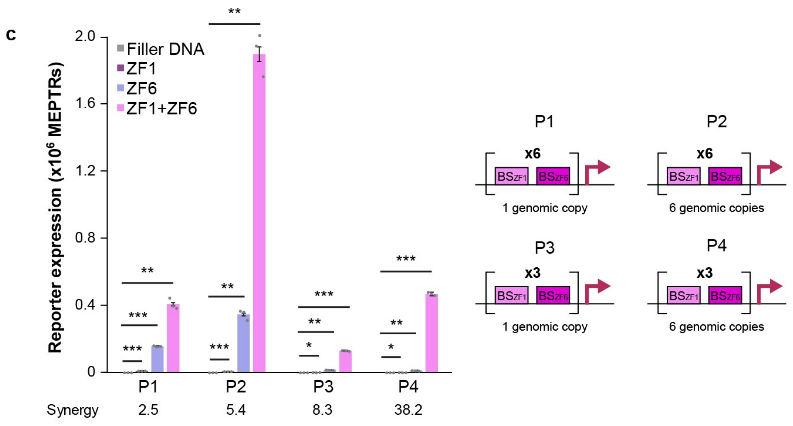


**Supplementary Figure 9. Supplementary information for multiplexing NatE MESA receptors to perform logical evaluation of multiple inputs.**
**(c)** Synthetic hybrid promoter development for AND gate implementation. Four promoter architectures (P1-P4) were built and inserted into the AAVS1 safe harbor locus of Landing Pad cells (HEK293FT-LP). The promoters have interspersed binding sites for COMET synTFs ZF1 and ZF6, with a total of either 6 or 12 total binding sites[7]. Soluble transcription factors were expressed by transfection in the engineered cell lines and reporter expression was measured to evaluate the synergistic potential of the hybrid promoter architectures. Synergy is defined in **Supplementary Note 7**. Results for two-tailed Welch's t-test between different ZF conditions are indicated above corresponding bars for significant pairs only (* $p < 0.05$, ** $p < 0.01$, *** $p < 0.001$). Each bar represents the mean of transfected cells across three biologic replicates and error bars indicate standard error of the mean (S.E.M). Abbreviations: ZFX, zinc finger X; PX, promoter design X; MEPTRs, molecules of equivalent PE-TexasRed.

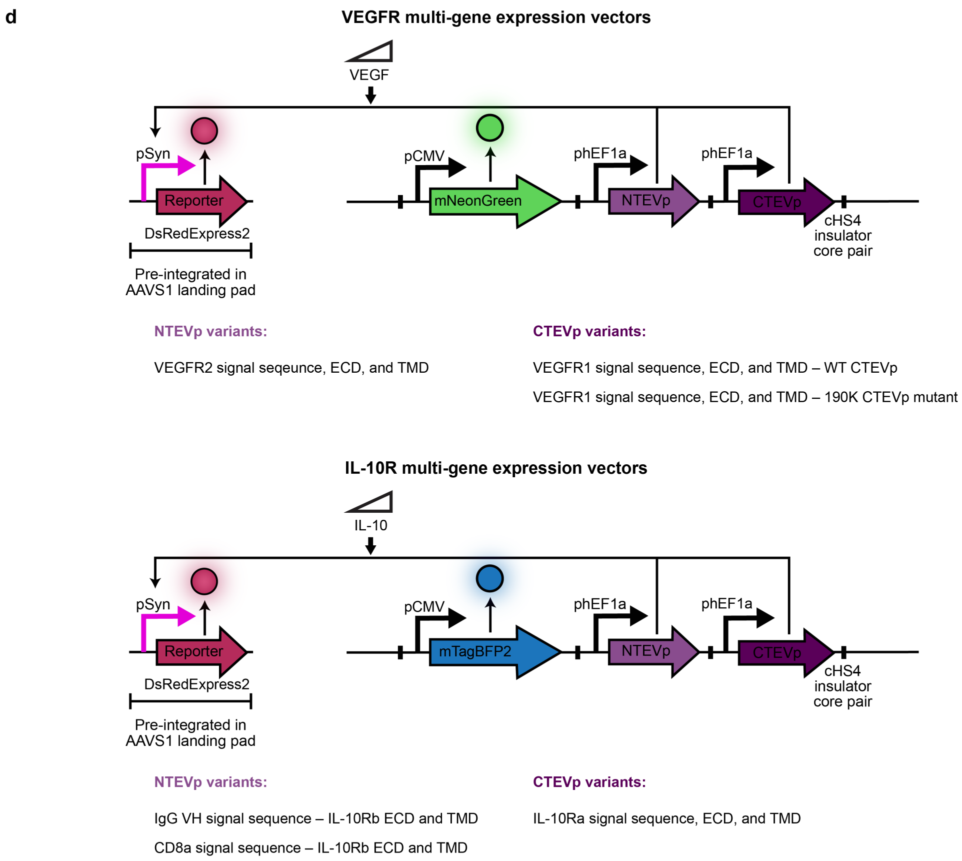


**Supplementary Figure 9. Supplementary information for multiplexing NatE MESA receptors to perform logical evaluation of multiple inputs.**
**(d)** Multi-gene expression vector (MGEV) design for the evaluation of AND gate logic. Each receptor (VEGF and IL-10 NatE MESA) was encoded on a different MGEV containing both the NTEVp and CTEVp chains, as well as a fluorescent proxy (mNeonGreen for VEGF, mTagBFP2 for IL-10). In all vector designs, transcriptional units (containing the promoter, the gene, and the terminator) are flanked by a pair of cHS4 insulators. See **Supplementary Figure 9c** for synthetic promoter design choices. Abbreviations: cHS4, chicken hypersensitive site 4 insulator; TEVp, Tobacco Etch Virus protease; NTEVp, N-terminal component of split, mutant TEVp; CTEVp, C-terminal component of split, mutant TEVp; VEGFR, vascular endothelial growth factor receptor; IL-10R, interleukin-10 receptor; CMV, cytomegalovirus; hEF1a, human elongation factor alpha; ECD, ectodomain; TMD, transmembrane domain.

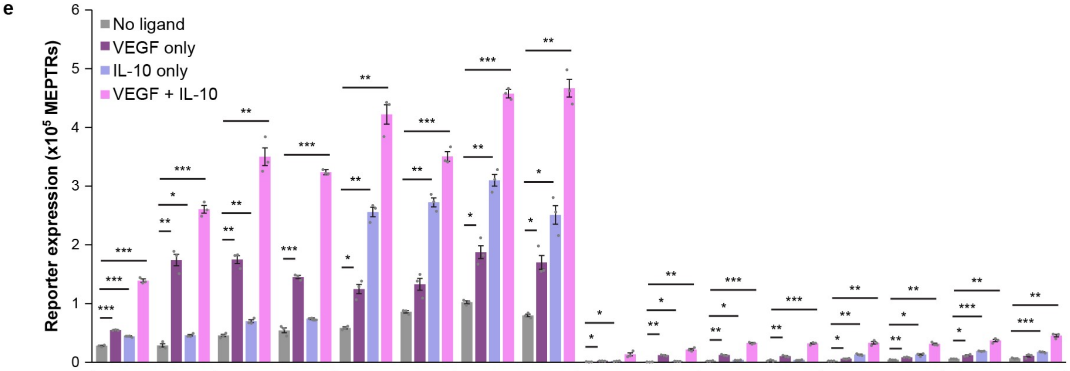


| | | | | | | | | | | | | | | | | |
|---|---|---|---|---|---|---|---|---|---|---|---|---|---|---|---|---|
| **VEGF NTEVp** | 75S | | WT | | 75S | | WT | | 75S | | WT | | 75S | | WT | |
| **VEGF ZF** | ZF6 | | | | ZF1 | | | | ZF6 | | | | ZF1 | | | |
| **IL-10 signal sequence** | hIgGVH | hCD8a | hIgGVH | hCD8a | hIgGVH | hCD8a | hIgGVH | hCD8a | hIgGVH | hCD8a | hIgGVH | hCD8a | hIgGVH | hCD8a | hIgGVH | hCD8a |
| **IL-10 ZF** | ZF1 | | | | ZF6 | | | | ZF1 | | | | ZF6 | | | |
| **Reporter** | P2 | | | | | | | | P4 | | | | | | | |
| **Synergy** | 2.6 | 1.4 | 2.0 | 2.4 | 1.4 | 1.1 | 1.2 | 1.5 | 5.1 | 1.7 | 2.7 | 3.6 | 2.1 | 2.1 | 1.6 | 2.6 |

**Supplementary Figure 9. Supplementary information for multiplexing NatE MESA receptors to perform logical evaluation of multiple inputs.**

**(e)** Full panel of hybrid promoter AND gate design choices. VEGF and IL-10 NatE MESA receptor-encoding MGEVs (see **Supplementary Figure 9d**) were co-transfected in the engineered P2 and P4 reporter cell lines along with the different co-expressed ligands (or empty vector filler DNA in the no ligand condition). Reporter expression was measured for a panel of design choices. The VEGF receptor pair used in this panel is a NTEVp receptor chain with VEGFR2 signal sequence, ECD and TMD, WT/75S NTEVp and a CTEVp receptor chain with VEGFR1 signal sequence, ECD and TMD, 190K CTEVp. The IL-10 receptor pair used in this panel is a NTEVp receptor chain with CD8a/IgGVH signal sequence, IL-10Rb ECD and TMD, 75S NTEVp and a CTEVp receptor chain with IL-10Ra signal sequence, IL-10Ra ECD and TMD, 190K CTEVp. CTEVp chains were not varied between each of the receptors. Synergy values are shown below the plot and synergy is defined in **Supplementary Note 7**. Results for two-tailed Welch's t-tests between different ligand conditions are indicated above corresponding bars for significant pairs only (* $p < 0.05$, ** $p < 0.01$, *** $p < 0.001$). Each bar represents the mean of transfected cells across three biologic replicates and error bars indicate standard error of the mean (S.E.M). Abbreviations: VEGF, vascular endothelial growth factor; IL-10, interleukin-10; TEVp, Tobacco Etch Virus protease; NTEVp, N-terminal component of split, mutant TEVp; CTEVp, C-terminal component of split, mutant TEVp; ZFX, zinc finger X; WT, wildtype; MEPTRs, molecules of equivalent PE-TexasRed; MGEV, multi-gene expression vector.

f

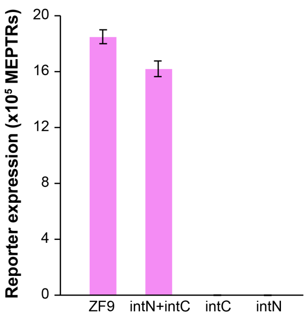


**Supplementary Figure 9. Supplementary information for multiplexing NatE MESA receptors to perform logical evaluation of multiple inputs.**
**(f)** Validation of split intein synTF (ZF9) function. Plasmids expressing soluble whole ZF9 and intN/intC split ZF9 were transfected into a reporter cell line (DsRedExpress2 reporter driven by pZF9), and reporter expression was measured. When expressed individually, neither of the split ZF9 halves induce any reporter expression. When co-expressed, the split ZF9 halves can reconstitute and induce reporter expression at a level comparable to the whole ZF9 protein. Each bar represents the mean of transfected cells across one biologic replicate and error bars indicate standard error of the mean (S.E.M) across the population. Abbreviations: ZF, zinc finger; MEPTRs, molecules of equivalent PE-TexasRed; intN, N-terminal component of split intein; intC, C-terminal component of split intein.

**g**

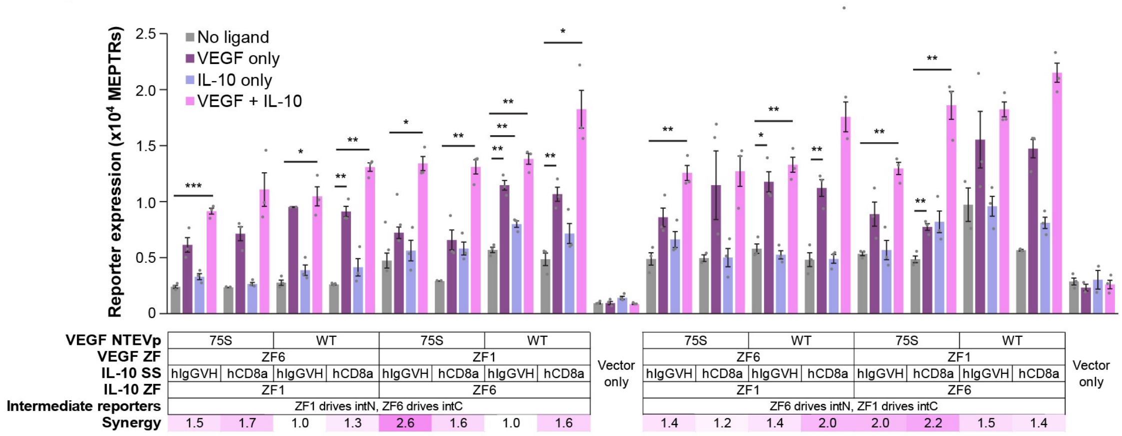


**Supplementary Figure 9. Supplementary information for multiplexing NatE MESA receptors to perform logical evaluation of multiple inputs.**

**(g)** Full panel of split intein synTF AND gate design choices. VEGF and IL-10 NatE MESA-encoding MGEVs (see **Supplementary Figure 9d**) were co-transfected in the engineered reporter cell lines (see **Figure 4f-g**) along with plasmids encoding the different co-expressed ligands (or empty vector filler DNA in the no ligand condition). Reporter expression was measured for a panel of design choices. The VEGF receptor pair used in this panel is a NTEVp receptor chain with VEGFR2 signal sequence, ECD and TMD, WT/75S NTEVp and a CTEVp receptor chain with VEGFR1 signal sequence, ECD and TMD, 190K CTEVp. The IL-10 receptor pair used in this panel is a NTEVp receptor chain with CD8a/IgGVH signal sequence, IL-10Rb ECD and TMD, 75S NTEVp and a CTEVp receptor chain with IL-10Ra signal sequence, IL-10Ra ECD and TMD, 190K CTEVp. CTEVp chains were not varied between each of the receptor pairs. Synergy values are shown below the plot and synergy is defined in **Supplementary Note 7**. Results for two-tailed Welch's t-tests between different ligand conditions are indicated above corresponding bars for significant pairs only (* $p$ < 0.05, ** $p$ < 0.01, *** $p$ < 0.001). Each bar represents the mean of transfected cells across three biologic replicates and error bars indicate standard error of the mean (S.E.M). Abbreviations: VEGF, vascular endothelial growth factor; IL-10, interleukin-10; TEVp, Tobacco Etch Virus protease; NTEVp, N-terminal component of split, mutant TEVp; CTEVp, C-terminal component of split, mutant TEVp; SS, signal sequence; ZFX, zinc finger X; WT, wildtype; intN, N-terminal component of split intein; intC, C-terminal component of split intein; MEPTRs, molecules of equivalent PE-TexasRed; MGEV, multi-gene expression vector.

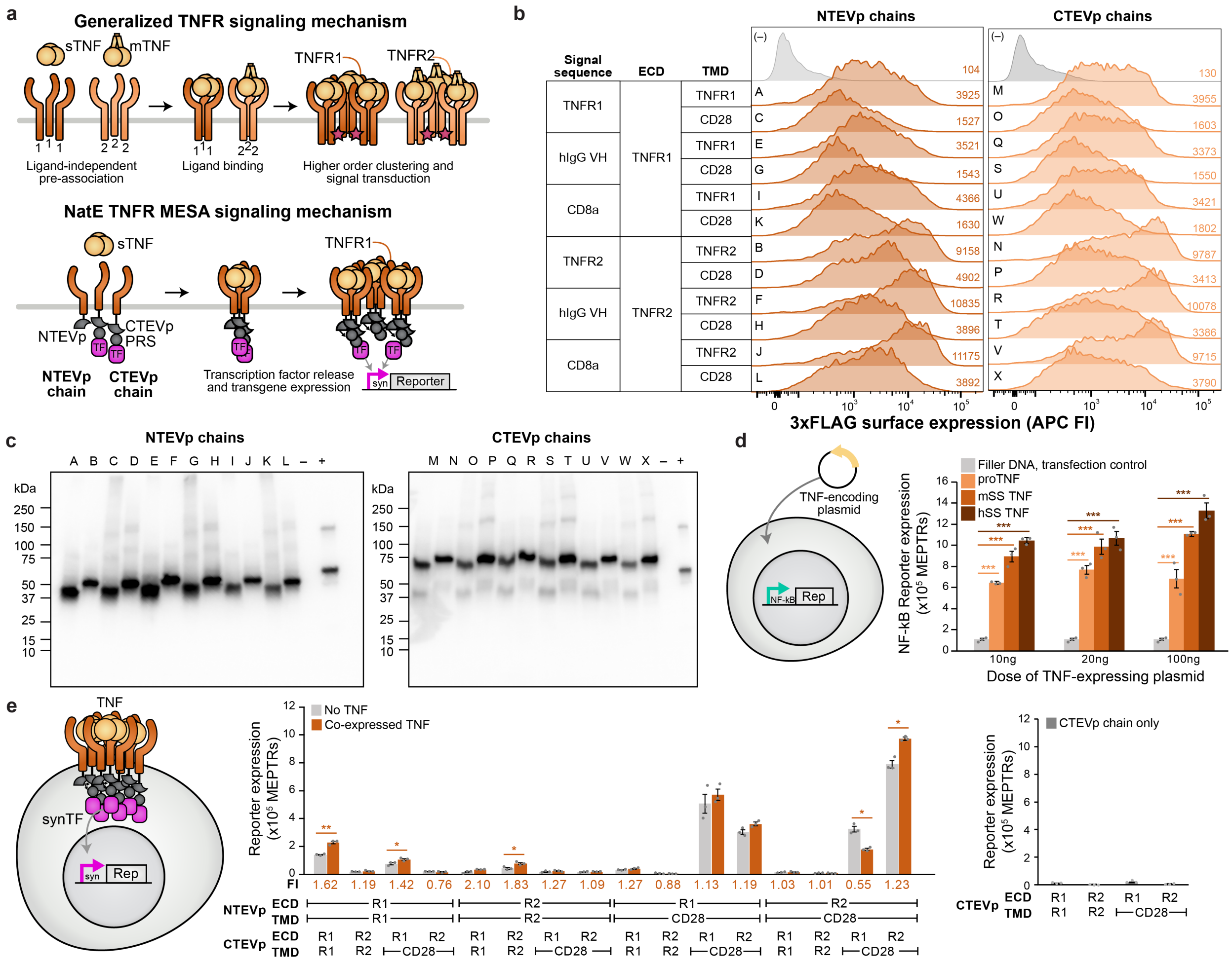


**Supplementary Figure 10. Supplementary information for conversion of TNFR.**
**(a)** Schematic of generalized TNFR signaling mechanism highlighting receptor interactions (top) and schematic of the proposed converted TNF NatE MESA signaling mechanism (bottom). **(b)** Surface expression of each single chain expressed in HEK293FT cells alone. Histograms show data for transfected (fluorescent) cells and the gray histograms in each column are transfection controls (no receptor). Mean APC fluorescent intensity for each sample is listed. **(c)** Whole-cell expression of each TNF NatE MESA receptor chain was measured via western blotting of a 3xFLAG epitope tag fused to the N-terminus of receptors. Soluble mNeonGreen (mNG) or mTagBFP2 (mTB2) were included as negative controls (–) because each receptor-expressing plasmid is a poly-transfection vector that also contains either a constitutively expressed mNG or mTB2 in the backbone. A rapamycin-sensing MESA receptor, pPD810 was included as an internal control for 3xFLAG-tagged protein expression (+)[2]. Letters indicate receptor variants labeled in detail in panel **(b). (d)** To validate bioactivity of the co-expressed TNF ligand, we transfected TNF-encoding plasmids into a previously validated stable cell line with a genomically integrated fluorescent NF-kB reporter. In HEK293FTs, TNF would be expected to activate a NF-kB reporter through endogenous TNF receptors. The three expression systems included proTNF (membrane bound version that is reflective of how natural TNF is produced before cleavage), and mature TNF that includes only the soluble portion with either a mouse or human Ig *κ* light chain leader sequence for secretion. All expression systems induced a significant increase in reporter expression (multi-factor ANOVA, *** $p < 0.001$). We proceeded with mature TNF with the human Ig *κ* light chain leader sequence. **(e)** Functional evaluation of TNF NatE MESA receptors across all mixed and matched ECDs, TMDs, and signaling domain pairings with and without co-expressed human TNF. CD28 mediates increased background and induced signal (multi-factor ANOVA, $p < 0.05$). Most configurations show low signal, with a moderate increase in signal for few pairs (two-tailed Welch's t-test results indicated above each bar pairing for * $p < 0.05$, ** $p < 0.01$). Single

CTEVp chains expressed alone were also tested and produced minimal reporter output (right). Abbreviations: TNFR, tumor necrosis factor receptor; sTNF, soluble TNF; mTNF, membrane-bound TNF; PRS, protease recognition sequence; TF, transcription factor; ECD, ectodomain; TMD, transmembrane domain; TEVp, Tobacco Etch Virus protease; NTEVp, N-terminal component of split, mutant TEVp; CTEVp, C-terminal component of split, mutant TEVp; APC FI, allophycocyanin fluorescence intensity; MEPTRs, molecules of equivalent PE-TexasRed; Rep, reporter; MFI, mean fluorescence intensity; FI, fold induction; R1, TNFR1; R2, TNFR2; proTNF, precursor TNF; mSS, murine signal sequence; hSS, human signal sequence.

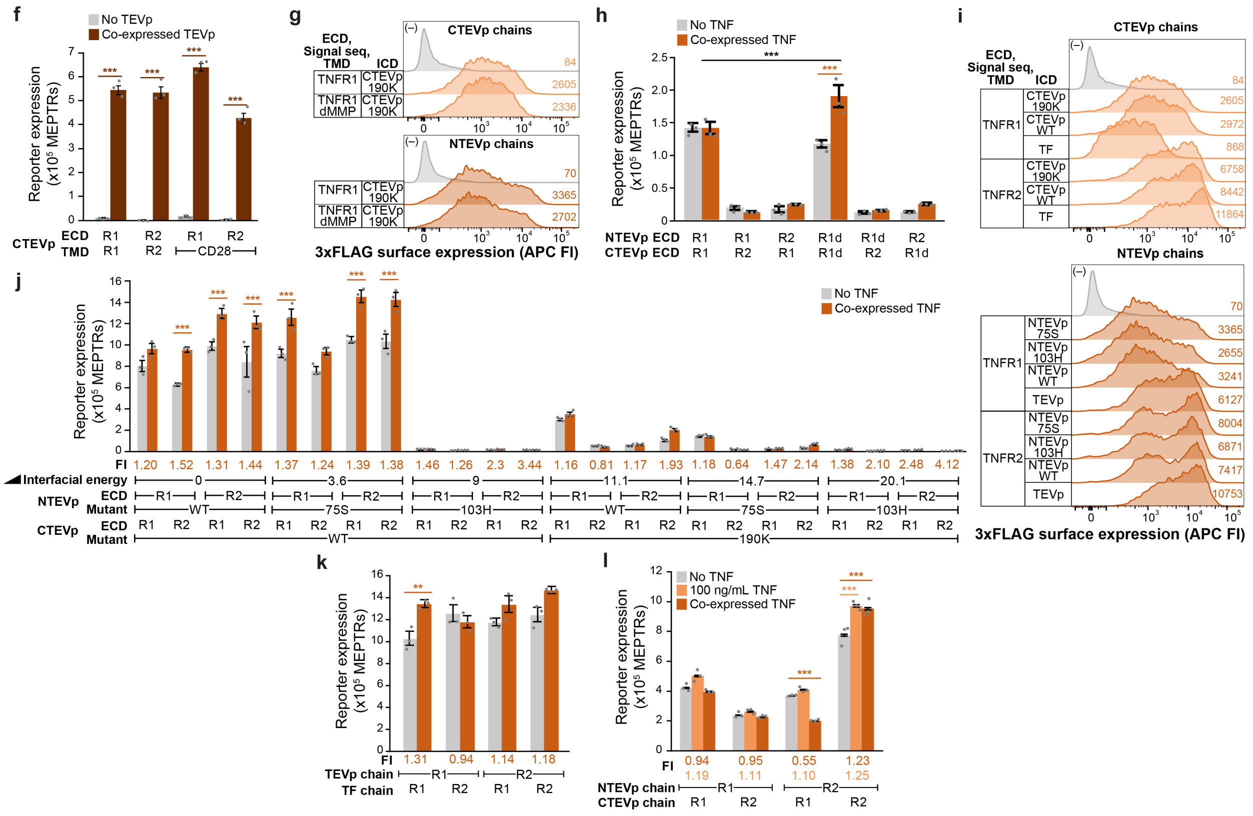

**Supplementary Figure 10. Supplementary information for conversion of TNFR.**
**(f)** To validate that CTEVp receptor chains could be cleaved by a reconstituted TEVp, we co-expressed each chain with or without soluble TEVp with a nuclear exclusion tag (NES) and found that all receptors could release synTFs to induce reporter expression in the presence of TEVp (multi-factor ANOVA, *** $p < 0.001$). **(g)** Removal of a MMP cleavage site was removed from TNFR1-based receptors to evaluate if reporter output could be increased. Surface expression to detect the 3xFLAG epitope tag was unchanged by these mutations. Histograms included all transfected cells based on expression of a constitutive fluorescent protein in the receptor-encoding plasmid backbone. **(h)** Induced reporter output was increased with mutations to remove a MMP cleavage site in TNFR1 domains when this mutated TNFR1 domain (R1d) is used on both the NTEVp and CTEVp receptor chains (multi-factor ANOVA, *** $p < 0.001$). **(i)** Surface expression to detect the 3xFLAG epitope tag of receptor chains with different NTEVp and CTEVp mutant domains was measured. Histograms included all transfected cells based on expression of a constitutive fluorescent protein in the receptor-encoding plasmid backbone. **(j)** Functional evaluation of TNF NatE MESA receptors across different NTEVp/CTEVp mutant pairings with and without co-expressed human TNF. Receptor pairs with NTEVp/CTEVp pairings with low interfacial energy conferred higher, inducible signaling (multi-factor ANOVA, *** $p < 0.001$). All receptors contain their native signal sequence and transmembrane domains. Interfacial energy is measured in Rosetta energy units (REUs). **(k)** Receptor pairings that employ a *trans*-cleavage signaling mechanism with a full TEVp on one chain and a recognition sequence on the other chain, which do not have to reconstitute the TEVp to signal, also displayed higher signaling. These receptors contain their native signal sequence and transmembrane domains. **(l)** Functional evaluation of TNF NatE MESA receptors treated with and without 250 ng/mL recombinant, exogenous ligand and co-transfected with TNF. All receptors contain a CD28 transmembrane domain. A transfection assay with extended timing (late ligand addition after a passage post transfection) shows some inducibility for the TNFR2-TNFR2 pairing (multi factor ANOVA, *** $p < 0.001$). For all bar graphs, bars represent the mean across transfected cells of three biologic replicates and error bars depict standard error of the mean (S.E.M.). Abbreviations: TNFR, tumor necrosis factor receptor; sTNF, soluble TNF; mTNF, membrane-bound TNF; PRS, protease recognition sequence; TF, transcription factor; ECD, ectodomain; TMD,

transmembrane domain; ICD, intracellular domain; TEVp, Tobacco Etch Virus protease; NTEVp, N-terminal component of split, mutant TEVp; CTEVp, C-terminal component of split, mutant TEVp; APC FI, allophycocyanin fluorescence intensity; MEPTRs, molecules of equivalent PE-TexasRed; Rep, reporter; MFI, mean fluorescence intensity; FI, fold induction; R1, TNFR1; R2, TNFR2; R1d, TNFR1 with deleted matrix metalloproteinase site; dMMP, deleted matrix metalloproteinase site.

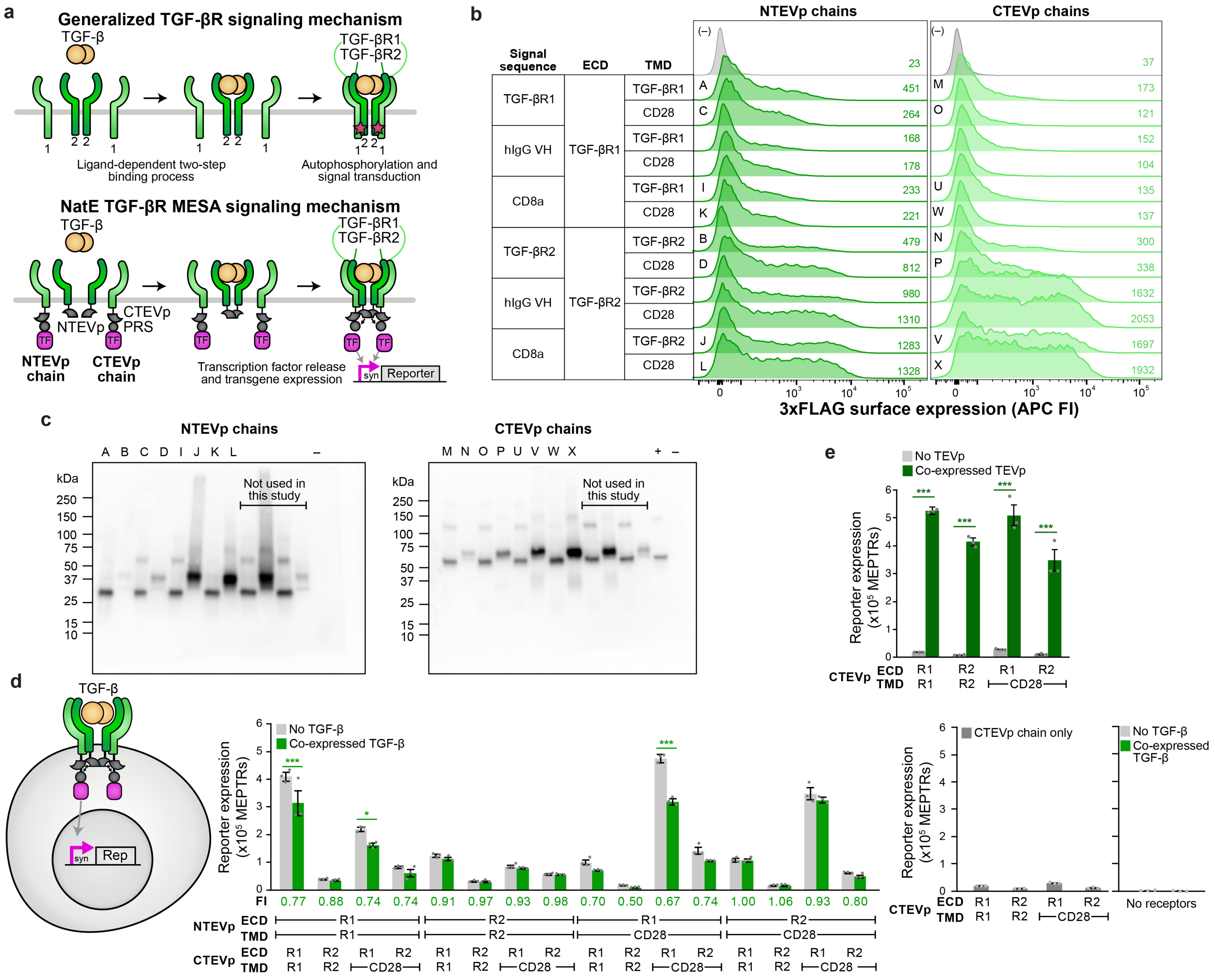

**Supplementary Figure 11. Supplementary information for conversion of TGF-βR.**
**(a)** Schematic of generalized TGF-βR signaling mechanism highlighting receptor interactions (top), and schematic of the proposed converted TGF-β NatE MESA signaling mechanism (bottom). **(b)** Surface expression of each single chain expressed by transfection in HEK293FT cells alone. Histograms show data for transfected (fluorescent) cells and the gray histograms in each column are transfection controls (no receptor). Mean APC fluorescence intensity for each sample is listed. **(c)** Whole-cell expression of each TGF-β NatE MESA chain was measured via western blotting of a 3xFLAG epitope tag fused to the N-terminus of receptors. Soluble mNeonGreen (mNG) or mTagBFP2 (mTB2) were included as negative controls (–) because each receptor-expressing plasmid is a poly-transfection vector that also contains either a constitutively expressed mNG or mTB2 in the backbone. A rapamycin-sensing MESA receptor, pPD810 was included as an internal control for 3xFLAG-tagged protein expression (+)[2]. Letters indicate receptor variants labeled in more detail in panel **(b). (d)** Functional evaluation of TGF-β NatE MESA receptors across all mixed and matched ECDs, TMDs, and signaling domain pairings with and without co-expressed human TGF-β. Using the CD28 TMD confers increased background signal, and most configurations show low signal, with a moderate increase in signal when two TGF-βR1 ECDs are employed. Single CTEVp chains transfected alone were also tested and produced minimal reporter output (right). **(e)** To validate that CTEVp receptors could be cleaved by a reconstituted TEVp, we co-expressed each chain with or without soluble TEVp and found that all receptors could release synTFs to induce reporter expression in the presence of TEVp. Abbreviations: TGF-BR, transforming growth factor beta receptor; PRS, protease recognition sequence; TF, transcription factor; ECD, ectodomain; TMD, transmembrane domain; TEVp, Tobacco Etch Virus protease; NTEVp, N-terminal component of split, mutant TEVp; CTEVp, C-terminal component of

split, mutant TEVp; APC FI, allophycocyanin fluorescence intensity; MEPTRs, molecules of equivalent PE-TexasRed; Rep, reporter; MFI, mean fluorescence intensity; FI, fold induction; R1, TGF-BR1; R2, TGF-BR2.

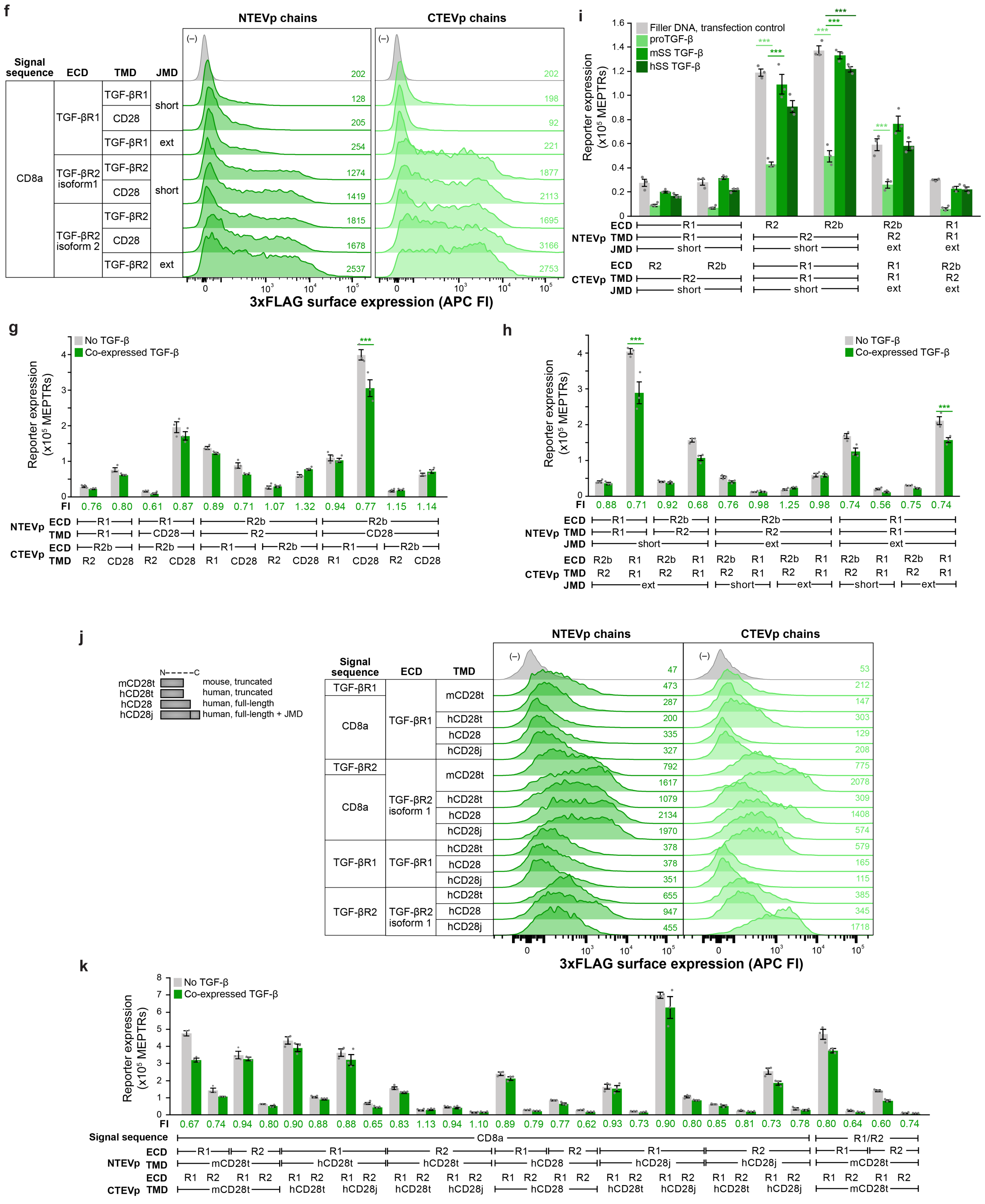


**Supplementary Figure 11. Supplementary information for conversion of TGF-βR.**

**(f)** Because TGF-β NatE MESA receptors demonstrated poorer surface expression than other NatE MESA receptors explored in this study, to improve surface expression, we investigated whether extending the juxtamembrane sequence to retain additional residues from the native receptor sequence could improve surface expression. We also evaluated receptors with an ectodomain derived from TGF-βR2 isoform 2. We found that switching to TGF-βR2 isoform 2 did increase surface expression moderately when incorporated on an NTEVp or CTEVp chain. The combination of switching from TGF-βR2 isoform 1 to isoform 2 and extending the JMD improved surface expression the most. For TGF-βR1, which was the poorer surface expresser, extending the JMD only minimally increased surface expression. **(g)** Functional evaluation of TGF-β NatE MESA receptors including the TGF-βR2 isoform 2 chains. No ligand-inducible receptor pairs were identified (multi-factor ANOVA, *** $p < 0.001$). **(h)** Functional evaluation of TGF-β NatE MESA receptors including the TGF-βR2 isoform 2 chains and mixed and matched JMD lengths. Again, no ligand-inducible increases in reporter expression were observed (multi-factor ANOVA, *** $p < 0.001$). **(i)** Evaluation of different TGF-β1 ligand co-expression systems. The three expression systems included proTGF-β (a pre-processed version with the native signal sequence and latency associated peptide domain), and mature TGF-β that includes only the soluble portion with either a mouse or human Ig *κ* light chain leader sequence for secretion. None of the expression systems induced a significant increase in reporter expression when tested with receptors with mismatched ectodomains (multi-factor ANOVA, *** $p < 0.001$). Some of the receptors demonstrated ligand de-inducibility when co-expressed with proTGF-β, where reporter expression with the ligand is significantly less than without the ligand (multi-factor ANOVA, *** $p < 0.001$). **(j)** To improve surface expression, we also evaluated switching to a human CD28 transmembrane domain with either the same truncated length used in other receptors employing a mouse CD28 transmembrane domain in this study, the full domain, or the full domain plus juxtamembrane domain. In general, changing the CD28 transmembrane domain variant did not substantially change surface expression. ECD and signal sequence have the biggest impact on surface expression. Histograms show data for transfected (fluorescent) cells and the gray histograms in each column are transfection controls (no receptor). Mean APC fluorescence intensity for each sample is listed. **(k)** Functional evaluation TGF-β NatE MESA receptors across different CD28 transmembrane domain variations. All receptors are not inducible with co-expressed TGF-β1. For all bar graphs, bars represent the mean across transfected cells of three biologic replicates and error bars depict standard error of the mean (S.E.M.). Abbreviations: TGF-BR, transforming growth factor beta receptor; PRS, protease recognition sequence; TF, transcription factor; ECD, ectodomain; TMD, transmembrane domain; JMD, juxtamembrane domain; TEVp, Tobacco Etch Virus protease; NTEVp, N-terminal component of split, mutant TEVp; CTEVp, C-terminal component of split, mutant TEVp; APC FI, allophycocyanin fluorescence intensity; MEPTRs, molecules of equivalent PE-TexasRed; Rep, reporter; MFI, mean fluorescence intensity; FI, fold induction; R1, TGF-BR1; R2, TGF-BR2; R2b, TGF-BR2 isoform 2; mCD28t, murine CD28 transmembrane domain truncated; hCD28t, human CD28 transmembrane domain truncated; mCD28j, murine CD28 transmembrane and juxtamembrane domain; hCD28j, human CD28 transmembrane and juxtamembrane domain; pro TGF-B, precursor TGF-B; mSS, murine signal sequence; hSS, human signal sequence.

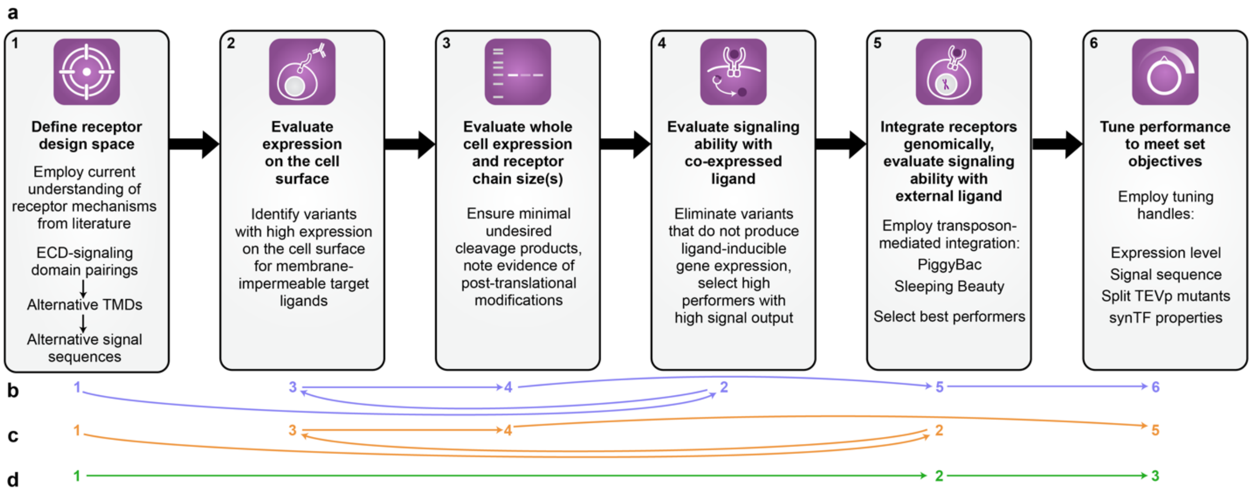


**Supplementary Figure 12. Synthesizing design principles for conversion of natural human receptors into synthetic biosensors.**
(**a-d**) A proposed workflow for converting natural receptors into synthetic NatE MESA receptors. **(a)** Black outlined boxes and black numbers depict the process as it was implemented in this study. This workflow employs a strategy to down-select receptor variants based on surface and whole cell expression before testing function. **(b-d)** Colored arrows and corresponding numbers depict alternative workflows depending on desired properties and throughput. **(b)** The blue workflow prioritizes testing receptor function (in co-expressed ligand setup with most optimal receptor-ligand interactions for signaling) before characterizing expression properties and doing so only for select variants or to explain functional differences. **(c)** The orange workflow prioritizes testing receptor function first in the application context (e.g., genomically integrated receptors and exogenous, recombinant ligand) and only evaluating expression properties for functional hits. The green workflow **(d)** only focuses on biosensor function in the application context (e.g., genomically integrated receptors and exogenous, recombinant ligand). The alternative workflows might facilitate higher throughput explorations because surface expression and western blotting can limit throughput, though they provide useful information about which receptor variants have desired properties of surface expression and correct protein size. All workflows start with considerations for defining the receptor design space and end with options for tuning performance.

### SUPPLEMENTARY TABLES

**Supplementary Table 1. Soluble ligand concentrations used throughout this study.**

| Ligand | Product information | Vehicle | Stock concentration | Working concentration |
|---|---|---|---|---|
| hVEGF-165 | Biolegend #583706 | PBS | 100 µg/mL | 100 ng/mL |
| | Biolegend #583704 | PBS | 200 µg/mL | 100 ng/mL |
| hIL-10 | Biolegend #573206 | PBS | 200 µg/mL | 100, 250 ng/mL |
| hTNF | ACROBiosystems, ActiveMax #TNA-H4211 | PBS | 25 µg/mL | 250 ng/mL |

**Supplementary Table 2. Instrument specifications for analytical flow cytometry.**

| Instrument | Fluorescent protein | Parameter/Channel name | Excitation laser | Filter set |
|---|---|---|---|---|
| BD LSR Fortessa | mTagBFP2 | Pacific Blue | Violet, 405 nm | 450/50 |
| | mNeonGreen | FITC | Blue, 488 nm | 505LP, 530/30 |
| | DsRedExpress2 | PE-Texas Red | Yellow Green, 550 nm | 600LP, 610/20 |
| | miRFP720 | Alexa Fluor 750 | Far Red, 690 nm | 690LP, 730/45 |

**Supplementary Table 3. Instrument specifications for fluorescence-activated cell sorting.**

| Instrument | Fluorescent protein | Parameter/Channel name | Excitation laser | Filter set |
|---|---|---|---|---|
| BD FacsAria IIu | mNeonGreen | FITC | Blue, 488 nm | 505LP, 530/30 |
| | DsRedExpress2 | PE-Texas Red | Yellow Green, 561 nm | 600LP, 610/20 |
| | miRFP720 | APC-Cy7 | Red, 633 nm | 690LP, 730/45 |

**Supplementary Table 4. UniProt entries for natural receptors investigated in this study.**

| Receptor name | Gene name | Species | UniProt ID |
|---|---|---|---|
| VEGFR1 | FLT1 | Human | P17948 |
| VEGFR2 | KDR | | P35968 |
| IL-10Rα | IL10RA | | Q13651 |
| IL-10Rβ | IL10RB | | Q08334 |
| TNFR1 | TNFRSF1A | | P19438 |
| TNFR2 | TNFRSF1B | | P20333 |
| TGF-βR1 | TGFBR1 | | P36897 |
| TGF-βR2 | TGFBR2 | | P37173 |

### SUPPLEMENTARY NOTES

#### Supplementary Note 1. *Detailed statistical results for single factor ANOVAs and Tukey's HSD tests*

Below are the outcomes from single factor ANOVAs and Tukey's HSD tests. Null hypotheses were that there existed no effects of ligand treatment on the measured reporter expression.

Functional signaling assay in **Figure 1d**
VEGFR-based NatE MESA, transfected receptors (VEGFR2 NTEVp, VEGFR1 CTEVp)

- Treatment $p = 2\times10^{-16}$
  - Plasmid doses of co-expressed VEGF165 above and equal to 2 ng induced a significant increase in reporter expression compared to the 0 ng case (all $p < 0.05$).

Functional signaling assay in **Figure 2f**, **Supplementary Figure 7r**
IL-10R-based NatE MESA, PiggyBac transposon cell line

- Treatment $p = 2.11\times10^{-13}$
  - Doses of recombinant IL-10 above and equal to 16 ng/mL induced a significant increase in reporter expression compared to the 0 ng/mL case (all $p < 0.05$).
  - Doses of recombinant IL-10 less than and equal to 8 ng/mL did not induce a significant increase in reporter expression compared to the 0 ng/mL case (all $p > 0.05$).

IL-10R-based NatE MESA, Sleeping Beauty transposon cell line

- Treatment $p < 2\times10^{-16}$
  - Doses of recombinant IL-10 above and equal to 16 ng/mL induced a significant increase in reporter expression compared to the 0 ng/mL case (all $p < 0.05$).
  - Doses of recombinant IL-10 less than and equal to 8 ng/mL did not induce a significant increase in reporter expression compared to the 0 ng/mL case (all $p > 0.05$).

IL-10R-based NatE MESA, sorted PiggyBac transposon cell line

- Treatment $p = 1.68\times10^{-13}$
  - Doses of recombinant IL-10 above and equal to 64 ng/mL induced a significant increase in reporter expression compared to the 0 ng/mL case (all $p < 0.05$).
  - Doses of recombinant IL-10 less than and equal to 32 ng/mL did not induce a significant increase in reporter expression compared to the 0 ng/mL case (all $p > 0.05$).

Functional signaling assay in **Figure 3b**
IL-10R-based NatE MESA, PiggyBac Jurkat transposon cell line (puromycin-selected)

- Treatment $p = 5.45\times10^{-8}$
  - Doses of recombinant IL-10 above and equal to 32 ng/mL induced a significant increase in reporter expression compared to the 0 ng/mL case (all $p < 0.05$).
  - Doses of recombinant IL-10 less than and equal to 16 ng/mL did not induce a significant increase in reporter expression compared to the 0 ng/mL case (all $p > 0.05$).

IL-10R-based NatE MESA, PiggyBac Jurkat transposon cell line (sorted mNeonGreen octile 1)

- Treatment $p = 1.28\times10^{-11}$
  - Doses of recombinant IL-10 above and equal to 2 ng/mL induced a significant increase in reporter expression compared to the 0 ng/mL case (all $p < 0.05$).
  - Doses of recombinant IL-10 less than and equal to 1 ng/mL did not induce a significant increase in reporter expression compared to the 0 ng/mL case (all $p > 0.05$).

Below are the outcomes from single-factor ANOVAs and Tukey's HSD tests. Null hypotheses were that there existed no effects of ECD, TMD, or their interaction on the measured reporter expression with or without co-expressed ligand.

Functional signaling assay in **Supplementary Figure 10e**
No ligand (background signal)

- NTEVp ECD $p = 0.1160$ (n.s.)

- NTEVp TMD $p < 2\times10^{-16}$
  - Differences between all TMDs were significant (all $p < 0.05$).

- CTEVp ECD $p = 0.3370$ (n.s.)
- CTEVp TMD $p < 2\times10^{-16}$
  - Differences between all TMDs were significant (all $p < 0.05$).
- Interaction between NTEVp ECD and CTEVp ECD $p = 8.08\times10^{-13}$
- Interaction between NTEVp TMD and CTEVp ECD $p = 2.73\times10^{-8}$
- Interaction between NTEVp ECD and CTEVp TMD $p = 6.80\times10^{-9}$
- Interaction between NTEVp TMD and CTEVp TMD $p < 2\times10^{-16}$

**Supplementary Note 2.** ***Detailed statistical results for multi-factor ANOVAs and Tukey's HSD tests***

Below are the outcomes from multi-factor ANOVAs and Tukey's HSD tests. Null hypotheses were that there existed no effects on the measured reporter expression of NTEVp ECD, NTEVp TMD, CTEVp ECD, CTEV TMD, co-expressed ligand, interactions between: NTEVp ECD and CTEVp ECD, NTEVp TMD and CTEVp ECD, NTEVp ECD and CTEVp TMD, NTEVp TMD and CTEVp TMD, NTEVp ECD and treatment, NTEVp TMD and treatment, CTEVp ECD and treatment, CTEVp TMD and treatment, NTEVp ECD and CTEVp ECD and treatment, NTEVp TMD and CTEVp ECD and treatment, NTEVp ECD and CTEVp TMD and treatment, NTEVp TMD and CTEVp TMD and treatment.

Functional signaling assay in **Figure 1c**

- Treatment $p < 2\times10^{-16}$
  - Each secreted ligand induces a statistically significant increase in reporter expression compared to the no ligand condition ($p < 0.05$).
  - The reporter expression with VEGF165 is significantly different than the reporter expression with VEGF 121 ($p < 0.05$).
- NTEVp ECD $p = 0.2729$ (not significant, n.s.)
- NTEVp TMD $p < 2\times10^{-16}$
  - The differences between VEGFR1 and CD28 TMDs, as well as VEGFR2 and CD28 TMDs are statistically significant ($p < 0.05$).
  - No statistical significance was found between VEGFR1 and VEGFR2 TMDs ($p$ = 0.9733).
- CTEVp ECD $p < 2\times10^{-16}$
  - The differences between VEGFR1 and VEGFR2 ECDs are significant ($p < 0.05$).
- CTEVp TMD $p < 2\times10^{-16}$
  - The differences between VEGFR1 and CD28 TMDs, as well as VEGFR2 and CD28 TMDs are statistically significant ($p < 0.05$).
  - No statistical significance was found between VEGFR1 and VEGFR2 TMDs ($p$ = 0.1328).
- All two-way interactions between independent variables were found to be statistically significant ($p < 0.05$) except NTEVp ECD and treatment interaction.

Below are the outcomes from multi-factor ANOVAs and Tukey's HSD tests. Null hypotheses were that there existed no effects of signal sequence pairing, co-expressed ligand, or their interaction on the measured reporter expression.

Functional signaling assay in **Supplementary Figure 6c**

- Signal sequence pairing $p < 2\times10^{-16}$
- Ligand treatment $p < 2\times10^{-16}$
- Interaction between signal sequence pairing and ligand treatment $p < 2\times10^{-16}$

Below are the outcomes from multi-factor ANOVAs and Tukey's HSD tests. Null hypotheses were that there existed no effects of interfacial energy between split TEVp halves, ligand treatment, or their interaction on the measured reporter expression.

Functional signaling assay in **Supplementary Figure 6f**

- Interfacial energy $p < 2\times10^{-16}$
- Ligand treatment $p = 7.33\times10^{-11}$
- Interaction between interfacial energy and ligand treatment $p = 3.35\times10^{-10}$
  - The ligand-treated condition is significantly different for the WT NTEVp/190K CTEVp pair and the 75S NTEVp/190K CTEVp pair (ligand-treated condition, interfacial energy 11.1 vs. 14.7, $p = 0.0004733$).

Below are the outcomes from multi-factor ANOVAs and Tukey's HSD tests. Null hypotheses were that there existed no effects of experiment replicate, ligand treatment, or their interaction on reporter expression or the percent of cells with an active reporter.

Functional signaling assay in **Supplementary Figure 6h**
Effect on reporter expression:

- Experiment replicates $p = 0.0375$
  - Only experiment 1 and 3 had a statistically significant difference ($p = 0.0317$).
- Ligand treatment $p = 0.2943$ (not significant, n.s.)
- Interaction between interfacial energy and ligand treatment $p = 0.7633$ (n.s.)

Effect on percent of cells with an active reporter:

- Experiment replicates $p = 3.77\times10^{-8}$
  - All differences were significant (all $p < 0.05$).
- Ligand treatment $p = 0.167$ (not significant, n.s.)
- Interaction between interfacial energy and ligand treatment $p = 0.733$ (n.s.)

Below are the outcomes from multi-factor ANOVAs and Tukey's HSD tests. Null hypotheses were that there existed no effects of sodium butyrate concentration, timepoint of sodium butyrate addition, ligand treatment, or their interaction on the percent of cells with an active reporter.

Functional signaling assay in **Supplementary Figure 6i**

- Timepoint of sodium butyrate addition $p = 5.19\times10^{-15}$
  - All differences were significant (all $p < 0.05$).
- Sodium butyrate concentration $p < 2\times10^{-16}$
  - All differences were significant (all $p < 0.05$).
- Ligand treatment $p = 0.000887$
- Interaction between timepoint and concentration of sodium butyrate $p = 6x10^{-12}$
- All other interaction terms were not significant ($p > 0.05$).

Below are the outcomes from multi-factor ANOVAs and Tukey's HSD tests. Null hypotheses were that there existed no effects of ligand treatment, time after treatment, or their interaction on the measured reporter expression.

Functional signaling assay in **Figure 2g**

- Treatment $p < 2\times10^{-16}$
- Time after treatment $p < 2\times10^{-16}$
  - A significant increase in reporter expression compared to the 0 timepoint is observed at timepoints of 18 h and later for ligand-treated cells (ligand-treated condition, time 0 vs. time 18 h and later, all $p < 0.05$).
  - No significant increase in reporter expression compared to the 0 timepoint is observed at any timepoint for untreated cells (no treatment condition, time 0 vs. no treatment condition at all other times, all $p > 0.05$).
- Interaction between treatment and time after treatment $p < 2\times10^{-16}$
  - A significant increase in reporter expression compared to the non-treated condition at a matching timepoint is observed at timepoints of 22 h and later (ligand-treated condition vs. no treatment condition at time 22 h and later, all $p < 0.05$).

Below are the outcomes from multi-factor ANOVAs and Tukey's HSD tests. Null hypotheses were that there existed no effects of signal sequence, TMD, co-expressed ligand, or their interaction on the measured reporter expression.

Functional signaling assay in **Supplementary Figure 7d**

- NTEVp signal sequence $p < 2\times10^{-16}$
  - All differences were significant (all $p < 0.05$).

- NTEVp TMD $p < 2\times10^{-16}$
  - All differences were significant (all $p < 0.05$).
- CTEVp signal sequence $p < 2\times10^{-16}$
  - All differences were significant (all $p < 0.05$).
- CTEVp TMD $p < 2\times10^{-16}$
  - All differences were significant (all $p < 0.05$).
- Co-expressed ligand $p < 2\times10^{-16}$
- Interaction between NTEVp signal sequence, NTEVp TMD, CTEVp signal sequence, CTEVp TMD, and co-expressed ligand $p = 0.1458$
  - Comparisons between co-expressed ligand and no ligand for all receptor pairs were significant (all $p < 0.05$).

Below are the outcomes from multi-factor ANOVAs and Tukey's HSD tests. Null hypotheses were that there existed no effects of ligand treatment, sorted octile of mNeonGreen expression, or their interaction on the measured reporter expression.

Functional signaling assay in **Supplementary Figure 7j**
- Treatment $p < 2\times10^{-16}$
- Octile $p < 2\times10^{-16}$
  - Differences between each octile were significant at each treatment condition (no treatment, 100 ng/mL IL-10, and 250 ng/mL IL-10) (all $p < 0.05$).
- Interaction between treatment and octile $p < 2\times10^{-16}$

Functional signaling assay in **Supplementary Figure 7n**
- Treatment $p < 2\times10^{-16}$
- Octile $p < 2\times10^{-16}$
  - All differences between octiles were not significant at each treatment condition (no treatment, 100 ng/mL IL-10, and 250 ng/mL IL-10) (all $p > 0.05$) except for the difference between octile 1 and octile 2 ($p < 0.05$).
- Interaction between treatment and octile $p < 2\times10^{-16}$

Below are the outcomes from multi-factor ANOVAs and Tukey's HSD tests. Null hypotheses were that there existed no effects of ligand treatment, sorted octile of mNeonGreen expression, or their interaction on the measured percentage of cells expressing reporter.

Functional signaling assay in **Supplementary Figure 7k**
- Treatment $p < 2\times10^{-16}$
- Octile $p < 2\times10^{-16}$
  - Differences between each octile were significant at each treatment condition (no treatment, 100 ng/mL IL-10, and 250 ng/mL IL-10) (all $p < 0.05$).
- Interaction between treatment and octile $p = 0.084$ (n.s.)

Functional signaling assay in **Supplementary Figure 7o**
- Treatment $p < 2\times10^{-16}$
- Octile $p < 2\times10^{-16}$
  - Differences between octiles 1 and 2, 3 and 4 were significant at each treatment condition (no treatment, 100 ng/mL IL-10, and 250 ng/mL IL-10) (all $p < 0.05$).
  - Difference between octiles 2 and 3 with 250 ng/mL IL-10 was significant ($p < 0.05$).
  - All other differences were not significant (all $p > 0.05$).
- Interaction between treatment and octile $p = 0.084$ (n.s.)

Below are the outcomes from multi-factor ANOVAs and Tukey's HSD tests. Null hypotheses were that there existed no effects of ligand treatment, natural TF, or their interaction on the measured reporter expression.

Functional signaling assay in **Figure 3g**

- Ligand treatment $p = 1.69\times10^{-8}$
  - No differences were significant for unmodified Jurkats (all $p > 0.05$).
  - No differences were significant for Jurkats engineered with Tbet-releasing receptors (all $p > 0.05$).
  - Both ligand treatments induced a significant increase for Jurkats engineered with cJun-releasing receptors and reporter 1 (all $p < 0.05$).
  - Treatment with 100 ng/mL ligand induced a significant increase for Jurkats engineered with cJun-releasing receptors and reporter 2 ($p < 0.05$) but treatment with 250 ng/mL ligand did not induce a significant increase ($p > 0.05$).
  - Both ligand treatments induced a significant increase for Jurkats engineered with BATF-releasing receptors (all $p < 0.05$).
- Natural TF $p < 2\times10^{-16}$
  - Background reporter expression with no ligand was significantly different between Jurkats engineered with BATF-releasing receptors and all other TFs, unmodified Jurkats and Jurkats engineered with cJun1- and BATF-releasing receptors, and Jurkats engineered with Tbet- and cJun1-releasing receptors (all $p < 0.05$). Background reporter expression with no ligand was not significantly different between all other comparisons (all $p > 0.05$).
  - Reporter expression with 100 ng/mL ligand treatment was significantly different between all comparisons (all $p < 0.05$) except for between unmodified Jurkats and Jurkats engineered with Tbet-releasing receptors ($p > 0.05$).
  - Reporter expression with 250 ng/mL ligand treatment was significantly different between all comparisons (all $p < 0.05$) except for between unmodified Jurkats and Jurkats engineered with Tbet-releasing receptors ($p > 0.05$).
- Interaction between treatment and natural TF $p = 4.04\times10^{-7}$

Functional signaling assay in **Supplementary Figure 8h**

- Ligand treatment $p = 0.0007$
  - No differences were significant for unmodified Jurkats (all $p > 0.05$).
  - No differences were significant for Jurkats engineered with Tbet-releasing receptors (all $p > 0.05$).
  - No differences were significant for Jurkats engineered with cJun-releasing receptors and reporter 1 (AP-1 binding sites) (all $p > 0.05$).
  - No differences were significant for Jurkats engineered with cJun-releasing receptors and reporter 2 (cJun dimer binding sites) (all $p > 0.05$).
  - Both ligand treatments induced a significant increase for Jurkats engineered with BATF-releasing receptors (all $p < 0.05$).
- Natural TF $p < 2\times10^{-16}$
  - Background reporter expression with no ligand was significantly different between Jurkats engineered with BATF-releasing receptors and all other TFs, and Jurkats engineered with BATF-releasing receptors and unmodified Jurkats (all $p < 0.05$). Background reporter expression with no ligand was not significantly different between all other comparisons (all $p > 0.05$).
  - Reporter expression with 100 ng/mL ligand treatment was significantly different between Jurkats engineered with BATF-releasing receptors and all other TFs, Jurkats engineered with BATF-releasing receptors and unmodified Jurkats, and Jurkats engineered with Tbet-releasing receptor and with cJun-releasing receptors with reporter 1 (all $p < 0.05$). All other comparisons were not significantly different (all $p > 0.05$).
  - Reporter expression with 250 ng/mL ligand treatment was significantly different between Jurkats engineered with BATF-releasing receptors and all other TFs, and Jurkats engineered with BATF-releasing receptors and unmodified Jurkats (all $p < 0.05$). All other comparisons were not significantly different (all $p > 0.05$).

- Interaction between treatment and natural TF $p = 5.73\times10^{-6}$

Functional signaling assay in **Supplementary Figure 8i**

- Ligand treatment $p = 0.1462$ (n.s.)
  - Ligand treatment with 100 ng/mL or 250 ng/mL IL-10 did not induce a significant increase in reporter expression for unmodified Jurkats and for all natural TF reporters (all $p > 0.05$).
- Natural TF $p < 2\times10^{-16}$
  - Background reporter expression with no ligand was not significantly different between Jurkats engineered with cJun reporter 2 (cJun dimer binding sites) and cJun reporter 1 (AP-1 binding sites), and between unmodified Jurkats and Jurkats engineered with a Tbet reporter (all $p > 0.05$). All other comparisons were statistically significant (all $p < 0.05$).
- Interaction between treatment and natural TF $p = 0.0392$ (n.s.)

Below are the outcomes from multi-factor ANOVAs and Tukey's HSD tests. Null hypotheses were that there existed no effects of natural TF identity, natural TF plasmid mass, or their interaction on the measured reporter expression.

Functional signaling assay in **Supplementary Figure 8g**

- Natural TF identity $p < 2\times10^{-16}$
  - Background reporter expression with no transfected natural TF plasmid was significantly different between all conditions (all $p < 0.05$) except for between the Tbet and cJun 2 conditions and between the BATF and cJun 1 conditions (both $p > 0.05$).
- Natural TF plasmid mass $p < 2\times10^{-16}$
  - cJun 1 (AP-1 binding sites)- The high natural TF plasmid mass induced a significant increase in reporter expression compared to the reporter only condition while the low and medium masses induced a significant decrease in reporter expression compared to the reporter only condition (all $p < 0.05$). Increasing the plasmid mass from low to medium and medium to high induced a significant increase in reporter expression (all $p < 0.05$).
  - cJun 2 (cJun dimer binding sites)- All natural TF plasmid masses induced a significant increase in reporter expression compared to the reporter only condition (all $p < 0.05$). The low and medium plasmid masses were significantly different from each other ($p < 0.05$) but the medium and high masses were not significantly different from each other ($p > 0.05$).
  - Tbet- All natural TF plasmid masses induced a significant increase in reporter expression compared to the reporter only condition (all $p < 0.05$). The low, medium, and high plasmid masses were not significant different from each other (all $p > 0.05$).
  - BATF- All natural TF plasmid masses induced a significant increase in reporter expression compared to the reporter only condition (all $p < 0.05$). The low and medium plasmid masses were significantly different from each other ($p < 0.05$) but the medium and high masses were not significantly different from each other ($p > 0.05$).
- Interaction between natural TF identity and plasmid dose $p = 3.04\times10^{-16}$

Below are the outcomes from multi-factor ANOVAs and Tukey's HSD tests. Null hypotheses were that there existed no effects of VEGFR pair, IL-10R pair, co-expressed ligand, or their interaction on the measured reporter expression.

Functional signaling assay in **Supplementary Figure 9b**

- VEGFR pair $p = 0.00852$
  - All differences were significant (all $p < 0.05$).
- IL-10R pair $p = 0.6201$ (not significant, n.s.)

- Ligand treatment $p < 2\times10^{-16}$
  - All differences were significant (all $p < 0.05$).
- All interaction terms were not significant, except for the interaction between the VEGFR pair and ligand treatment ($p = 6.96\times10^{-7}$)
  - Comparisons between conditions with the same ligand treatment but different VEGFR pair were not significant.
  - All other differences were significant (all $p < 0.05$).

Below are the outcomes from multi-factor ANOVAs and Tukey's HSD tests. Null hypotheses were that there existed no effects of TNF expression plasmid, TNF expression plasmid mass, or their interaction on the measured NF-kB reporter expression.

Functional signaling assay in **Supplementary Figure 10d**

- TNF expression plasmid $p = 4.26\times10^{-13}$
  - The mature TNF expression plasmids with mSS induced significantly more reporter expression than the proTNF-expressing plasmid at a high plasmid mass (all $p < 0.05$).
  - The mature TNF expression plasmids with hSS induced significantly more reporter expression than the proTNF-expressing plasmid at low, medium, and high plasmid masses (all $p < 0.05$).
- TNF expression plasmid mass $p = 0.0013$
  - All TNF plasmid masses induced a significant increase in reporter expression compared to a filler DNA transfection control (all $p < 0.05$).
- Interaction between TNF expression plasmid and plasmid mass $p = 0.0349$

Below are the outcomes from multi-factor ANOVAs and Tukey's HSD tests. Null hypotheses were that there existed no effects of CTEVp ECD-TMD, co-expressed TEV protease, or their interaction on the measured reporter expression.

Functional signaling assay in **Supplementary Figure 10f**

- CTEVp ECD-TMD $p = 5.97\times10^{-7}$
  - Comparisons between all CTEVp ECD-TMDs were significant (all $p < 0.05$) except for between R1-R1 and R2-R2 and between R2-R2 and R2-CD28 (all $p > 0.05$).
- Co-expressed TEVp $p < 2\times10^{-16}$
  - Co-expression of TEVp induced a significant increase in reporter expression compared to the no TEVp condition for all CTEVp chains (all $p < 0.05$).
- Interaction between CTEVp ECD-TMD and co-expressed TEVp $p = 5.82\times10^{-7}$

Below are the outcomes from multi-factor ANOVAs and Tukey's HSD tests. Null hypotheses were that there existed no effects of CTEVp ECD, NTEVp ECD, co-expressed ligand, or their interaction on the measured reporter expression.

Functional signaling assay in **Supplementary Figure 10h**

- NTEVp ECD $p = 2.62\times10^{-14}$
  - Replacement of R1 with R1d did not significantly impact background signal or induced signal with co-expressed ligand (all $p > 0.05$).
- CTEVp ECD $p < 2\times10^{-16}$
  - Replacement of R1 with R1d did not significantly impact background signal or induced signal with co-expressed ligand (all $p > 0.05$).
- Co-expressed ligand $p = 0.000162$
  - Co-expression of ligand with the R1d/R1d receptor pair induced a significant increase in reporter expression (all $p < 0.05$) while all others did not (all $p > 0.05$).
- Interaction between NTEVp ECD and CTEVp ECD $p = 0.1172$ (n.s.)

  - Replacement of R1 with R1d on both NTEVp and CTEVp did not significantly impact background signal ($p > 0.05$) but did significantly increase induced signal with co-expressed ligand ($p < 0.05$).
- Interaction between NTEVp ECD and co-expressed ligand $p = 0.0001$
- Interaction between CTEVp ECD and co-expressed ligand $p = 0.0001$

Below are the outcomes from multi-factor ANOVAs and Tukey's HSD tests. Null hypotheses were that there existed no effects of CTEVp ECD, CTEVp mutation, NTEVp ECD, NTEVp mutation, co-expressed ligand, or their interaction on the measured reporter expression.

Functional signaling assay in **Supplementary Figure 10j**

- NTEVp ECD $p = 2.13 \times 10^{-10}$
  - Comparison between TNFR1 and TNFR2 ECDs were significant (all $p < 0.05$).
- NTEVp mutation $p < 2 \times 10^{-16}$
  - Comparisons between all mutations were significant (all $p < 0.05$).
- CTEVp ECD $p = 2.33 \times 10^{-10}$
  - Comparison between TNFR1 and TNFR2 ECDs were significant (all $p < 0.05$).
- CTEVp mutation $p < 2 \times 10^{-16}$
  - Comparisons between all mutations were significant (all $p < 0.05$).
- Co-expressed ligand $p < 2 \times 10^{-16}$
  - Co-expressed ligand induced a significant increase in reporter expression for many receptor pairs with interfacial energies of 0 and 3.6 REUs (NTEVp ECD-mutation/CTEVp ECD-mutation: R1-WT/R2-WT, R2-WT/R1-WT, R2-WT/R2-WT, R1-75S/R1-WT, R2-75S/R1-WT, R2-75S/R2-WT) (all $p < 0.05$). Co-expressed ligand did not induce a significant increase for all other pairs (all $p > 0.05$).
- All 2nd order interactions are significant ($p < 0.05$) except for those between CTEVp ECD and co-expressed ligand and between CTEVp ECD and CTEVp mutation.

Below are the outcomes from multi-factor ANOVAs and Tukey's HSD tests. Null hypotheses were that there existed no effects of TEVp chain ECD, TF chain ECD, co-expressed ligand, or their interaction on the measured reporter expression.

Functional signaling assay in **Supplementary Figure 10k**

- TEVp chain ECD $p = 0.0100$
  - Comparison between TNFR1 and TNFR2 ECDs were significant (all $p < 0.05$).
- TF chain ECD $p = 0.0889$ (n.s.)
  - Comparisons between all mutations were not significant (all $p > 0.05$).
- Co-expressed ligand $p = 0.0005$
  - Co-expressed ligand induced a significant increase in reporter expression for receptors with a TNFR1 ECD on both chains (all $p < 0.05$). Co-expressed ligand did not induce a significant increase for all other pairs (all $p > 0.05$).
- Interaction between TEVp chain ECD and TF chain ECD $p = 0.3642$ (n.s.)
- Interaction between TEVp chain ECD and co-expressed ligand $p = 0.3151$ (n.s.)
- Interaction between TF chain ECD and co-expressed ligand $p = 0.0345$

Below are the outcomes from multi-factor ANOVAs and Tukey's HSD tests. Null hypotheses were that there existed no effects of NTEVp ECD, CTEVp ECD, co-expressed or recombinant ligand, or their interaction on the measured reporter expression.

Functional signaling assay in **Supplementary Figure 10l**

- NTEVp ECD $p < 2 \times 10^{-16}$
  - Comparison between TNFR1 and TNFR2 ECDs were significant (all $p < 0.05$).
- CTEVp ECD $p = 2.35 \times 10^{-15}$

    - Comparison between TNFR1 and TNFR2 ECDs were significant (all $p < 0.05$).
- Ligand treatment $p = 1.98\times10^{-7}$
    - Co-expressed ligand induced a significant increase in reporter expression for receptors with a TNFR2 ECD on the NTEVp chain and either a TNFR1 or TNFR2 ECD on the CTEVp chain (all $p < 0.05$). Co-expressed ligand did not induce a significant increase for all other pairs (all $p > 0.05$).
    - Treatment with 100 ng/mL recombinant TNF induced a significant increase in reporter expression for receptors with a TNFR2 ECD on both the NTEVp and CTEVp chains ($p < 0.05$). Treatment with 100 ng/mL recombinant TNF did not induce a significant increase for all other pairs (all $p > 0.05$).
- Interaction between NTEVp ECD and CTEVp ECD $p < 2\times10^{-16}$
- Interaction between NTEVp ECD and ligand treatment $p = 0.0576$ (n.s.)
- Interaction between CTEVp ECD and ligand treatment $p = 1.12\times10^{-6}$

Below are the outcomes from multi-factor ANOVAs and Tukey's HSD tests. Null hypotheses were that there existed no effects of NTEVp ECD-TMD, CTEVp ECD-TMD, co-expressed TGFb1, or their interaction on the measured reporter expression.

Functional signaling assay in **Supplementary Figure 11d**

- NTEVp ECD-TMD $p < 2\times10^{-16}$
    - Comparisons between all NTEVp ECD-TMDs were significant (all $p < 0.05$) except for between R1-R1 and R1-CD28 (all $p > 0.05$).
- CTEVp ECD-TMD $p < 2\times10^{-16}$
    - Comparisons between all CTEVp ECD-TMDs were significant (all $p < 0.05$).
- Co-expressed ligand $p = 2.59\times10^{-11}$
    - Co-expression of ligand induced a significant decrease in reporter expression compared to the no ligand condition for pairs NTEVp ECD-TMD/CTEVp ECD-TMD: R1-R1/R1-R1, R1-R1/R1-CD28, R1-CD28/R1-CD28 (all $p < 0.05$). All other receptor pairs did not induce a significant change in reporter expression (all $p > 0.05$).
- Interaction between NTEVp ECD-TMD and CTEVp ECD-TMD $p < 2\times10^{-16}$
- Interaction between NTEVp ECD-TMD and ligand treatment $p = 8.03\times10^{-7}$
- Interaction between CTEVp ECD-TMD and ligand treatment $p = 3.08\times10^{-6}$

Below are the outcomes from multi-factor ANOVAs and Tukey's HSD tests. Null hypotheses were that there existed no effects of NTEVp ECD-TMD, CTEVp ECD-TMD, co-expressed TGFb1, or their interaction on the measured reporter expression.

Functional signaling assay in **Supplementary Figure 11g**

- NTEVp ECD-TMD $p < 2\times10^{-16}$
    - Comparisons between all NTEVp ECD-TMDs were significant (all $p < 0.05$).
- CTEVp ECD-TMD $p < 2\times10^{-16}$
    - Comparisons between all CTEVp ECD-TMDs were significant (all $p < 0.05$).
- Co-expressed ligand $p < 2\times10^{-16}$
    - Co-expression of ligand induced a significant decrease in reporter expression compared to the no ligand condition for the NTEVp ECD-TMD/CTEVp ECD-TMD pair R2b-CD28/R1-CD28 (all $p < 0.05$). All other pairs did not induce significant change in reporter expression with co-expressed ligand (all $p > 0.05$).
- All second order interactions were statistically significant all $p < 2\times10^{-16}$.

Below are the outcomes from multi-factor ANOVAs and Tukey's HSD tests. Null hypotheses were that there existed no effects of NTEVp ECD-TMD-JMD, CTEVp ECD-TMD-JMD, co-expressed TGFb1, or their interaction on the measured reporter expression.

Functional signaling assay in **Supplementary Figure 11h**

- NTEVp ECD-TMD-JMD $p < 2\times10^{-16}$
  - Comparisons between all NTEVp ECD-TMDs were significant (all $p < 0.05$) except for between NTEVp ECD-TMD-JMD: R2b-R2-short and R1-R1-ext ($p > 0.05$).
- CTEVp ECD-TMD-JMD $p < 2\times10^{-16}$
  - Comparisons between all CTEVp ECD-TMDs were significant (all $p < 0.05$).
- Co-expressed ligand $p < 2\times10^{-16}$
  - Co-expression of ligand induced a significant decrease in reporter expression compared to the no ligand condition for the NTEVp ECD-TMD-JMD/CTEVp ECD-TMD-JMD pairs R1-R1-short/R1-R1-ext and R1-R1-ext/R1-R1-ext (all $p < 0.05$). All other pairs did not induce significant change in reporter expression with co-expressed ligand (all $p > 0.05$).
- All second order interactions were statistically significant all $p < 2\times10^{-16}$.

Below are the outcomes from multi-factor ANOVAs and Tukey's HSD tests. Null hypotheses were that there existed no effects of NTEVp ECD-TMD-JMD, CTEVp ECD-TMD-JMD, TGFb1 expression plasmid, or their interaction on the measured reporter expression.

Functional signaling assay in **Supplementary Figure 11i**

- NTEVp ECD-TMD-JMD $p < 2\times10^{-16}$
  - Comparisons between all NTEVp ECD-TMDs were significant (all $p < 0.05$) except for between NTEVp ECD-TMD-JMD: R2b-R2-short and R1-R1-ext ($p > 0.05$).
- CTEVp ECD-TMD-JMD $p < 2\times10^{-16}$
  - Comparisons between all CTEVp ECD-TMDs were significant (all $p < 0.05$) except for between CTEVp ECD-TMD-JMD: R2-R2-short and R2b-R2-short ($p > 0.05$).
- TGFb1 expression plasmid $p < 2\times10^{-16}$
  - Co-expression of TGFb1 induced a significant decrease in reporter expression compared to the no ligand condition for the NTEVp ECD-TMD-JMD/CTEVp ECD-TMD-JMD pairs R2-R2-short/R1-R1-short, R2b-R2-short/R1-R1-short, and R2b-R2-ext/R1-R1-ext with the proTGFb expression plasmid (all $p < 0.05$). All other pairs did not induce significant change in reporter expression with co-expressed ligand compared to the transfection control (all $p > 0.05$).
- All second order interactions were statistically significant all $p < 2\times10^{-16}$.

Below are the outcomes from multi-factor ANOVAs and Tukey's HSD tests. Null hypotheses were that there existed no effects of NTEVp SS-ECD-TMD, CTEVp SS-ECD-TMD, co-expressed TGFb1, or their interaction on the measured reporter expression.

Functional signaling assay in **Supplementary Figure 11k**

- NTEVp SS-ECD-TMD-JMD $p = 7.17\times10^{-6}$
- CTEVp SS-ECD-TMD-JMD $p = 6.02\times10^{-10}$
- Co-expressed TGFb1 $p = 0.521$
  - Co-expression of TGFb1 did not induce a significant change in reporter expression for all receptor pairs (all $p > 0.05$).
- Interaction between NTEVp SS-ECD-TMD and CTEVp SS-ECD-TMD $p = 1.41\times10^{-7}$.
- All other interactions were not statistically significant.

**Supplementary Note 3.** ***Detailed statistical results for Welch's t-tests***

Below are the outcomes of two-tailed Welch's *t*-tests followed by the Benjamini-Hochberg (BH) procedure. Null hypotheses were that there existed no effect of co-expressed ligand expression on the measured reporter expression.

Functional signaling assay in **Figure 1c**

(NTEVp ECD-TMD, CTEVp ECD-TMD: *p* value for comparison no VEGF vs. co-expressed VEGF165 or VEGF121)

- NTEVp R1-R1, CTEVp R1-R1, VEGF165: 0.004755
- NTEVp R1-R1, CTEVp R2-R2, VEGF165: 0.02098
- NTEVp R1-R1, CTEVp R1-CD28, VEGF165: $2.98x10^{-6}$
- NTEVp R1-R1, CTEVp R2-CD28, VEGF165: 0.001983
- NTEVp R2-R2, CTEVp R1-R1, VEGF165: 0.01493
- NTEVp R2-R2, CTEVp R2-R2, VEGF165: 0.1608 (not significant, n.s.)
- NTEVp R2-R2, CTEVp R1-CD28, VEGF165: $6.04x10^{-5}$
- NTEVp R2-R2, CTEVp R2-CD28, VEGF165: 0.03655
- NTEVp R1-CD28, CTEVp R1-R1, VEGF165: 0.000497
- NTEVp R1-CD28, CTEVp R2-R2, VEGF165: 0.003222
- NTEVp R1-CD28, CTEVp R1-CD28, VEGF165: 0.1583 (n.s.)
- NTEVp R1-CD28, CTEVp R2-CD28, VEGF165: 0.05485 (n.s.)
- NTEVp R2-CD28, CTEVp R1-R1, VEGF165: 0.00307
- NTEVp R2-CD28, CTEVp R2-R2, VEGF165: 0.3294 (n.s.)
- NTEVp R2-CD28, CTEVp R1-CD28, VEGF165: 0.03005
- NTEVp R2-CD28, CTEVp R2-CD28, VEGF165: 0.03635
- NTEVp R1-R1, CTEVp R1-R1, VEGF121: $1.21x10^{-5}$
- NTEVp R1-R1, CTEVp R2-R2, VEGF121: 0.03715
- NTEVp R1-R1, CTEVp R1-CD28, VEGF121: 0.000318
- NTEVp R1-R1, CTEVp R2-CD28, VEGF121: 0.001366
- NTEVp R2-R2, CTEVp R1-R1, VEGF121: 0.004607
- NTEVp R2-R2, CTEVp R2-R2, VEGF121: 0.01105
- NTEVp R2-R2, CTEVp R1-CD28, VEGF121: 0.000177
- NTEVp R2-R2, CTEVp R2-CD28, VEGF121: 0.213 (n.s.)
- NTEVp R1-CD28, CTEVp R1-R1, VEGF121: 0.000217
- NTEVp R1-CD28, CTEVp R2-R2, VEGF121: 0.01258
- NTEVp R1-CD28, CTEVp R1-CD28, VEGF121: 0.01879
- NTEVp R1-CD28, CTEVp R2-CD28, VEGF121: 0.01332
- NTEVp R2-CD28, CTEVp R1-R1, VEGF121: 0.007328
- NTEVp R2-CD28, CTEVp R2-R2, VEGF121: 0.009979
- NTEVp R2-CD28, CTEVp R1-CD28, VEGF121: 0.001163
- NTEVp R2-CD28, CTEVp R2-CD28, VEGF121: 0.2842 (n.s.)

Functional signaling assay in **Supplementary Figure 6c**
(NTEVp SS-ECD-TMD, CTEVp SS-ECD-TMD: *p* value for comparison no VEGF vs. co-expressed VEGF165)

- NTEVp R1-R1-R1, CTEVp R2-R2-CD28: 0.001157
- NTEVp R1-R1-R1, CTEVp IgGVH-R2-CD28: 0.009491
- NTEVp R1-R1-R1, CTEVp CD8a-R2-CD28: $7.378x10^{-5}$
- NTEVp IgGVH-R1-R1, CTEVp R2-R2-CD28: 0.002191
- NTEVp IgGVH-R1-R1, CTEVp IgGVH-R2-CD28: 0.002962
- NTEVp IgGVH-R1-R1, CTEVp CD8a-R2-CD28: $3.937x10^{-5}$
- NTEVp CD8a-R1-R1, CTEVp R2-R2-CD28: 0.007038
- NTEVp CD8a-R1-R1, CTEVp IgGVH-R2-CD28: 0.005922
- NTEVp CD8a-R1-R1, CTEVp CD8a-R2-CD28: 0.002485
- NTEVp R2-R2-R2, CTEVp R1-R1-R1: 0.006548
- NTEVp R2-R2-R2, CTEVp IgGVH-R1-R1: $5.539x10^{-4}$
- NTEVp R2-R2-R2, CTEVp CD8a-R1-R1: 0.001803
- NTEVp IgGVH-R2-R2, CTEVp R1-R1-R1: $8.274x10^{-4}$
- NTEVp IgGVH-R2-R2, CTEVp IgGVH-R1-R1: 0.001676
- NTEVp IgGVH-R2-R2, CTEVp CD8a-R1-R1: $1.803x10^{-5}$

- NTEVp CD8a-R2-R2, CTEVp R1-R1-R1: $5.368 \times 10^{-4}$
- NTEVp CD8a-R2-R2, CTEVp IgGVH-R1-R1: 0.004374
- NTEVp CD8a-R2-R2, CTEVp CD8a-R1-R1: $3.769 \times 10^{-5}$

Functional signaling assay in **Figure 1e**, and **Supplementary Figure 6f**
(NTEVp mutant, CTEVp mutant: *p* value for comparison no VEGF vs. co-expressed VEGF165)

- Full TEVp (protease chain), target chain: 0.1543 (not significant, n.s.)
- NTEVp WT, CTEVp WT: 0.7378 (n.s.)
- NTEVp 75S, CTEVp WT: 0.0854 (n.s.)
- NTEVp 75E, CTEVp WT: 0.0614 (n.s.)
- NTEVp WT, CTEVp 158P: 0.2431 (n.s.)
- NTEVp 75S, CTEVp 158P: 0.004421
- NTEVp WT, CTEVp 190K: 0.004541
- NTEVp 75E, CTEVp 158P: 0.1946 (n.s.)
- NTEVp 75S, CTEVp 190K: 0.003494
- NTEVp 75E, CTEVp 190K: 0.00026

Functional signaling assay in **Figure 2c**
(NTEVp ECD-TMD, CTEVp ECD-TMD: *p* value for comparison no IL-10 vs. co-expressed IL-10)

- NTEVp Ra-Ra, CTEVp Ra-Ra: 0.1039 (not significant, n.s.)
- NTEVp Ra-Ra, CTEVp Rb-Rb: 0.0536 (n.s.)
- NTEVp Ra-Ra, CTEVp Ra-CD28: 0.1060 (n.s.)
- NTEVp Ra-Ra, CTEVp Rb-CD28: 0.0025
- NTEVp Rb-Rb, CTEVp Ra-Ra: 0.0016
- NTEVp Rb-Rb, CTEVp Rb-Rb: 0.0182
- NTEVp Rb-Rb, CTEVp Ra-CD28: 0.0148
- NTEVp Rb-Rb, CTEVp Rb-CD28: 0.4330 (n.s.)
- NTEVp Ra-CD28, CTEVp Ra-Ra: 0.4071 (n.s.)
- NTEVp Ra-CD28, CTEVp Rb-Rb: 0.0952 (n.s.)
- NTEVp Ra-CD28, CTEVp Ra-CD28: 0.0016
- NTEVp Ra-CD28, CTEVp Rb-CD28: 0.0702 (n.s.)
- NTEVp Rb-CD28, CTEVp Ra-Ra: 0.0031
- NTEVp Rb-CD28, CTEVp Rb-Rb: 0.5485 (n.s.)
- NTEVp Rb-CD28, CTEVp Ra-CD28: 0.0011
- NTEVp Rb-CD28, CTEVp Rb-CD28: 0.0775 (n.s.)

Natural TF reporter assay in **Figure 3f**
(Natural TF: *p* value for comparison between no IL-10 and co-expression of IL-10)

- cJun 1: 0.0001
- cJun 2: 0.0002
- Tbet: 0.0001
- BATF: 0.0004

OR gate functional signaling assay in **Figure 4b**, and **Supplementary Figure 9b**
(VEGF NatE MESA receptor NTEVp mutant, IL-10 NatE MESA receptor NTEVp signal sequence, co-expressed ligand: *p* value for comparisons vs. no ligand condition)

- 75S, IgGVH, VEGF165 only: 0.0249
- 75S, IgGVH, IL-10 only: 0.00178
- 75S, IgGVH, VEGF165 + IL-10: 0.00182
- 75S, CD8a, VEGF165 only: 0.0237
- 75S, CD8a, IL-10 only: 0.000454
- 75S, CD8a, VEGF165 + IL-10: 0.0137
- WT, IgGVH, VEGF165 only: 0.00337

- WT, IgGVH, IL-10 only: $2.61X10^{-5}$
- WT, IgGVH, VEGF165 + IL-10: 0.00142
- WT, CD8a, VEGF165 only: 0.0143
- WT, CD8a, IL-10 only: $3.41x10^{-5}$
- WT, CD8a, VEGF165 + IL-10: $9.53x10^{-5}$

AND gate functional signaling assay in **Figure 4d**, and **Supplementary Figure 9e** (VEGFR NTEVp mutant and tethered ZF, IL-10R NTEVp signal sequence and tethered ZF, promoter architecture, co-expressed ligand: *p* value for comparisons vs. no ligand condition)

- 75S ZF6, IgGVH ZF1, P2, VEGF165 only: 0.00041
- 75S ZF6, IgGVH ZF1, P2, IL-10 only: 0.000293
- 75S ZF6, IgGVH ZF1, P2, VEGF165 + IL-10: 0.000176
- 75S ZF6, CD8a ZF1, P2, VEGF165 only: 0.00379
- 75S ZF6, CD8a ZF1, P2, IL-10 only: 0.0225
- 75S ZF6, CD8a ZF1, P2, VEGF165 + IL-10: 0.000102
- WT ZF6, IgGVH ZF1, P2, VEGF165 only: 0.00174
- WT ZF6, IgGVH ZF1, P2, IL-10 only: 0.00258
- WT ZF6, IgGVH ZF1, P2, VEGF165 + IL-10: 0.0029
- WT ZF6, CD8a ZF1, P2, VEGF165 only: 0.00046
- WT ZF6, CD8a ZF1, P2, IL-10 only: 0.0522 (not significant, n.s.)
- WT ZF6, CD8a ZF1, P2, VEGF165 + IL-10: $3.67x10^{-5}$
- 75S ZF1, IgGVH ZF6, P2, VEGF165 only: 0.0169
- 75S ZF1, IgGVH ZF6, P2, IL-10 only: 0.00144
- 75S ZF1, IgGVH ZF6, P2, VEGF165 + IL-10: 0.0025
- 75S ZF1, CD8a ZF6, P2, VEGF165 only: 0.0568 (n.s.)
- 75S ZF1, CD8a ZF6, P2, IL-10 only: 0.00116
- 75S ZF1, CD8a ZF6, P2, VEGF165 + IL-10: 0.000525
- WT ZF1, IgGVH ZF6, P2, VEGF165 only: 0.0199
- WT ZF1, IgGVH ZF6, P2, IL-10 only: 0.00183
- WT ZF1, IgGVH ZF6, P2, VEGF165 + IL-10: $1.4x10^{-5}$
- WT ZF1, CD8a ZF6, P2, VEGF165 only: 0.0216
- WT ZF1, CD8a ZF6, P2, IL-10 only: 0.011
- WT ZF1, CD8a ZF6, P2, VEGF165 + IL-10: 0.00161
- 75S ZF6, IgGVH ZF1, P4, VEGF165 only: 0.0112
- 75S ZF6, IgGVH ZF1, P4, IL-10 only: 0.0417
- 75S ZF6, IgGVH ZF1, P4, VEGF165 + IL-10: 0.0545 (n.s.)
- 75S ZF6, CD8a ZF1, P4, VEGF165 only: 0.00289
- 75S ZF6, CD8a ZF1, P4, IL-10 only: 0.0154
- 75S ZF6, CD8a ZF1, P4, VEGF165 + IL-10: 0.00671
- WT ZF6, IgGVH ZF1, P4, VEGF165 only: 0.00191
- WT ZF6, IgGVH ZF1, P4, IL-10 only: 0.0216
- WT ZF6, IgGVH ZF1, P4, VEGF165 + IL-10: 0.000304
- WT ZF6, CD8a ZF1, P4, VEGF165 only: 0.003137
- WT ZF6, CD8a ZF1, P4, IL-10 only: 0.4307 (n.s.)
- WT ZF6, CD8a ZF1, P4, VEGF165 + IL-10: 0.000171
- 75S ZF1, IgGVH ZF6, P4, VEGF165 only: 0.0159
- 75S ZF1, IgGVH ZF6, P4, IL-10 only: 0.00743
- 75S ZF1, IgGVH ZF6, P4, VEGF165 + IL-10: 0.00671
- 75S ZF1, CD8a ZF6, P4, VEGF165 only: 0.00225
- 75S ZF1, CD8a ZF6, P4, IL-10 only: 0.0251
- 75S ZF1, CD8a ZF6, P4, VEGF165 + IL-10: 0.00327
- WT ZF1, IgGVH ZF6, P4, VEGF165 only: 0.025
- WT ZF1, IgGVH ZF6, P4, IL-10 only: 0.000256

- WT ZF1, IgGVH ZF6, P4, VEGF165 + IL-10: 0.00543
- WT ZF1, CD8a ZF6, P4, VEGF165 only: 0.15 (n.s.)
- WT ZF1, CD8a ZF6, P4, IL-10 only: 0.000259
- WT ZF1, CD8a ZF6, P4, VEGF165 + IL-10: 0.00294

AND gate functional signaling assay in **Figure 4f**, and **Supplementary Figure 9g**
(VEGFR NTEVp mutant and tethered ZF, IL-10R NTEVp signal sequence and tethered ZF, promoter setup, co-expressed ligand: *p* value for comparisons vs. no ligand condition)

- 75S ZF6, IgGVH ZF1, Setup A, VEGF165 only: 0.037 (not significant, n.s.)
- 75S ZF6, IgGVH ZF1, Setup A, IL-10 only: 0.091 (n.s.)
- 75S ZF6, IgGVH ZF1, Setup A, VEGF165 + IL-10: 0.000363
- 75S ZF6, CD8a ZF1, Setup A, VEGF165 only: 0.345 (n.s.)
- 75S ZF6, CD8a ZF1, Setup A, IL-10 only: 0.481 (n.s.)
- 75S ZF6, CD8a ZF1, Setup A, VEGF165 + IL-10: 0.0813 (n.s.)
- WT ZF6, IgGVH ZF1, Setup A, VEGF165 only: 0.0671 (n.s.)
- WT ZF6, IgGVH ZF1, Setup A, IL-10 only: 0.182 (n.s.)
- WT ZF6, IgGVH ZF1, Setup A, VEGF165 + IL-10: 0.013
- WT ZF6, CD8a ZF1, Setup A, VEGF165 only: 0.00727
- WT ZF6, CD8a ZF1, Setup A, IL-10 only: 0.243 (n.s.)
- WT ZF6, CD8a ZF1, Setup A, VEGF165 + IL-10: 0.00168
- 75S ZF1, IgGVH ZF6, Setup A, VEGF165 only: 0.23 (n.s.)
- 75S ZF1, IgGVH ZF6, Setup A, IL-10 only: 0.89 (n.s.)
- 75S ZF1, IgGVH ZF6, Setup A, VEGF165 + IL-10: 0.0119
- 75S ZF1, CD8a ZF6, Setup A, VEGF165 only: 0.176 (n.s.)
- 75S ZF1, CD8a ZF6, Setup A, IL-10 only: 0.255 (n.s.)
- 75S ZF1, CD8a ZF6, Setup A, VEGF165 + IL-10: 0.00373
- WT ZF1, IgGVH ZF6, Setup A, VEGF165 only: 0.00215
- WT ZF1, IgGVH ZF6, Setup A, IL-10 only: 0.00744
- WT ZF1, IgGVH ZF6, Setup A, VEGF165 + IL-10: 0.00136
- WT ZF1, CD8a ZF6, Setup A, VEGF165 only: 0.00497
- WT ZF1, CD8a ZF6, Setup A, IL-10 only: 0.161 (n.s.)
- WT ZF1, CD8a ZF6, Setup A, VEGF165 + IL-10: 0.0154
- 75S ZF6, IgGVH ZF1, Setup B, VEGF165 only: 0.0455 (n.s.)
- 75S ZF6, IgGVH ZF1, Setup B, IL-10 only: 0.194 (n.s.)
- 75S ZF6, IgGVH ZF1, Setup B, VEGF165 + IL-10: 0.00218
- 75S ZF6, CD8a ZF1, Setup B, VEGF165 only: 0.152 (n.s.)
- 75S ZF6, CD8a ZF1, Setup B, IL-10 only: 0.466 (n.s.)
- 75S ZF6, CD8a ZF1, Setup B, VEGF165 + IL-10: 0.059 (n.s.)
- WT ZF6, IgGVH ZF1, Setup B, VEGF165 only: 0.0186
- WT ZF6, IgGVH ZF1, Setup B, IL-10 only: 0.465 (n.s.)
- WT ZF6, IgGVH ZF1, Setup B, VEGF165 + IL-10: 0.00291
- WT ZF6, CD8a ZF1, Setup B, VEGF165 only: 0.00639
- WT ZF6, CD8a ZF1, Setup B, IL-10 only: 0.941 (n.s.)
- WT ZF6, CD8a ZF1, Setup B, VEGF165 + IL-10: 0.0386 (n.s.)
- 75S ZF1, IgGVH ZF6, Setup B, VEGF165 only: 0.11 (n.s.)
- 75S ZF1, IgGVH ZF6, Setup B, IL-10 only: 0.775 (n.s.)
- 75S ZF1, IgGVH ZF6, Setup B, VEGF165 + IL-10: 0.00439
- 75S ZF1, CD8a ZF6, Setup B, VEGF165 only: 0.00477
- 75S ZF1, CD8a ZF6, Setup B, IL-10 only: 0.0939 (n.s.)
- 75S ZF1, CD8a ZF6, Setup B, VEGF165 + IL-10: 0.00915
- WT ZF1, IgGVH ZF6, Setup B, VEGF165 only: 0.542 (n.s.)
- WT ZF1, IgGVH ZF6, Setup B, IL-10 only: 0.442 (n.s.)
- WT ZF1, IgGVH ZF6, Setup B, VEGF165 + IL-10: 0.208 (n.s.)

- WT ZF1, CD8a ZF6, Setup B, VEGF165 only: 0.274 (n.s.)
- WT ZF1, CD8a ZF6, Setup B, IL-10 only: 0.773 (n.s.)
- WT ZF1, CD8a ZF6, Setup B, VEGF165 + IL-10: 0.0698 (n.s.)

Functional signaling assay in **Supplementary Figure 9a**
(NTEVp, CTEVp: *p* value for comparisons of no ligand vs. co-expressed ligand conditions below)

- VEGF 75S NTEVp mutant, IL-10 CTEVp, VEGF165 only: 0.000114
- VEGF 75S NTEVp mutant, IL-10 CTEVp, IL-10 only: 0.01068
- VEGF 75S NTEVp mutant, IL-10 CTEVp, VEGF165 + IL-10: 0.0224
- VEGF WT NTEVp, IL-10 CTEVp, VEGF165 only: 0.00751
- VEGF WT NTEVp, IL-10 CTEVp, IL-10 only: 0.0159
- VEGF WT NTEVp, IL-10 CTEVp, VEGF165 + IL-10: $3.06 \times 10^{-5}$
- IL-10 IgGVH signal sequence NTEVp, VEGF CTEVp, VEGF165 only: 0.988 (not significant, n.s.)
- IL-10 IgGVH signal sequence NTEVp, VEGF CTEVp, IL-10 only: 0.0137
- IL-10 IgGVH signal sequence NTEVp, VEGF CTEVp, VEGF165 + IL-10: 0.312 (n.s.)
- IL-10 CD8a signal sequence NTEVp, VEGF CTEVp, VEGF165 only: 0.308 (n.s.)
- IL-10 CD8a signal sequence NTEVp, VEGF CTEVp, IL-10 only: 0.291 (n.s.)
- IL-10 CD8a signal sequence NTEVp, VEGF CTEVp, VEGF165 + IL-10: 0.833 (n.s.)

Below are the outcomes of two-tailed Welch's *t*-tests followed by BH procedure. Null hypotheses were that there existed no effect of exogenous ligand treatment on the measured reporter expression.

Functional signaling assay in **Figure 1e**
(NTEVp mutant, CTEVp mutant: *p* value for comparison no VEGF vs. exogenous VEGF165)

- NTEVp WT, CTEVp 190K: 0.00172
- NTEVp 75S, CTEVp 190K: 0.00551
- NTEVp 75E, CTEVp 190K: 0.9914 (not significant, n.s.)

Functional signaling assay in **Figure 1f**

- All-in-one 75S NTEVp mutant, 0 vs. 100 ng/mL: 0.7439 (not significant, n.s.)
- All-in-one WT NTEVp, 0 vs. 100 ng/mL: 0.02101
- Receptors only 75S NTEVp mutant, 0 vs. 100 ng/mL: 0.5191 (n.s.)
- Receptors only WT NTEVp, 0 vs. 100 ng/mL: 0.8802 (n.s.)

Functional signaling assay in **Supplementary Figure 6d**
(NTEVp ECD-TMD, CTEVp ECD-TMD: *p* value for comparison no VEGF vs. exogenous VEGF165)

- NTEVp R1-R1, CTEVp R1-R1: 0.1081 (not significant, n.s.)
- NTEVp R1-R1, CTEVp R1-CD28: 0.2062 (n.s.)
- NTEVp R1-R1, CTEVp R2-CD28: 0.6378 (n.s.)
- NTEVp R2-R2, CTEVp R1-R1: 0.1427 (n.s.)
- NTEVp R2-R2, CTEVp R1-CD28: 0.8239 (n.s.)
- NTEVp R1-CD28, CTEVp R1-R1: 0.2582 (n.s.)
- NTEVp R1-CD28, CTEVp R2-R2: 0.9203 (n.s.)
- NTEVp R2-CD28, CTEVp R1-R1: 0.3769 (n.s.)

Functional signaling assay in **Supplementary Figure 1h**
(experiment number: *p* value for comparison no VEGF vs. co-expressed VEGF165)
Reporter expression plot:

- 1: 0.5244 (not significant, n.s.)
- 2: 0.5065 (n.s.)
- 3: 0.6429 (n.s.)

Percent of cells with active reporter plot:

- 1: 0.5574 (n.s.)

- 2: 0.13 (n.s.)
- 3: 0.7434 (n.s.)

Functional signaling assay in **Supplementary Figure 6i**
(timing of NaB addition, NaB dose: *p* value for comparison no VEGF vs. exogenous VEGF165)

- 24h, 0mM: 0.02711 (not significant, n.s.)
- 24h, 0.1mM: 0.008135
- 24h, 0.25mM: 0.05903 (n.s.)
- 24h, 0.5mM: 0.02991 (n.s.)
- 24h, 1mM: 0.02376 (n.s.)
- 24h, 2mM: 0.06088 (n.s.)
- 48h, 0mM: 0.1137 (n.s.)
- 48h, 0.1mM: 0.4599 (n.s.)
- 48h, 0.25mM: 0.01261
- 48h, 0.5mM: 0.01039
- 48h, 1mM: 0.1391 (n.s.)
- 48h, 2mM: 0.003519
- 72h, 0mM: 0.2704 (n.s.)
- 72h, 0.1mM: 0.0369 (n.s.)
- 72h, 0.25mM: 0.001372
- 72h, 0.5mM: 0.01095
- 72h, 1mM: 0.01178
- 72h, 2mM: 0.8232 (n.s.)

Functional signaling assay in **Figure 2d**

- No ligand vs. early ligand treatment only: 0.0043
- No ligand vs. late ligand treatment only: 0.0019
- No ligand vs. early and late ligand treatment: 0.0016
- Early ligand treatment only vs. late ligand treatment only: 0.0035
- Early ligand treatment only vs. early and late ligand treatment: 0.0024
- Late ligand treatment only vs. early and late ligand treatment: 0.0075

Functional signaling assay in **Figure 2e**

- All-in-one hIgG VH signal sequence, 0 vs. 100 ng/mL: 0.0130
- All-in-one hIgG VH signal sequence, 0 vs. 250 ng/mL: 0.0054
- All-in-one hIgG VH signal sequence, 100 vs. 250 ng/mL: 0.0142
- All-in-one CD8a signal sequence, 0 vs. 100 ng/mL: 0.0030
- All-in-one CD8a signal sequence, 0 vs. 250 ng/mL: 0.0018
- All-in-one CD8a signal sequence, 100 vs. 250 ng/mL: 0.0392 (not significant, n.s.)
- All-in-one inert receptors, 0 vs. 100 ng/mL: 0.257 (n.s.)
- All-in-one inert receptors, 0 vs. 250 ng/mL: 0.0435 (n.s.)
- All-in-one inert receptors, 100 vs. 250 ng/mL: 0.0308
- Receptors only hIgG VH signal sequence, 0 vs. 100 ng/mL: 0.0419 (n.s.)
- Receptors only hIgG VH signal sequence, 0 vs. 250 ng/mL: 0.0119
- Receptors only hIgG VH signal sequence, 100 vs. 250 ng/mL: 0.0276
- Receptors only CD8a signal sequence, 0 vs. 100 ng/mL: 0.0215
- Receptors only CD8a signal sequence, 0 vs. 250 ng/mL: 0.0314
- Receptors only CD8a signal sequence, 100 vs. 250 ng/mL: 0.9646 (n.s.)

Functional signaling assay in **Supplementary Figure 7g**
(Transposon type, signal sequence or receptor type: *p* value for comparison between untreated cells and 250 ng/mL IL-10 treatment)

- Sleeping Beauty, hIgG VH: 0.0040

- Sleeping Beauty, CD8a: 0.0006
- Sleeping Beauty, inert receptors: 0.2020 (n.s.)
- PiggyBac, hIgG VH: 0.0063
- PiggyBac, CD8a: 0.0003
- PiggyBac, inert receptors: 0.2797 (n.s.)

Functional signaling assay in **Supplementary Figure 7h**
(Cell line- PiggyBac engineered with receptors with varying signal sequences or non-modified: *p* value for comparison between untreated cells and 250 ng/mL IL-10 treatment.
- Unmodified HEK293FTs: 0.3086 (n.s.)
- hIgG VH: 0.0011
- CD8a: 0.0065

(Cell line- PiggyBac engineered with receptors with varying signal sequences or non-modified: *p* value for comparison between untreated cells and treatment with conditioned media from IL-10-secreting HEK293FT cells.
- Unmodified HEK293FTs: 0.1491 (n.s.)
- hIgG VH: $3.07 \times 10^{-5}$
- CD8a: 0.0007

(Cell line- PiggyBac engineered with receptors with varying signal sequences or non-modified: *p* value for comparison between treatment with 250 ng/mL IL-10 and treatment with conditioned media from IL-10-secreting HEK293FT cells.
- Unmodified HEK293FTs: 0.3231 (n.s.)
- hIgG VH: 0.0018
- CD8a: 0.0123

Below are the outcomes of two-tailed Welch's *t*-tests followed by BH procedure. Null hypotheses were that there existed no effect of exogenous ligand treatment or co-culture condition on the measured NFAT reporter expression.

NFAT reporter assay in **Figure 3e**
(Jurkat cell line: *p* value for comparison between untreated cells and 250 ng/mL IL-10 treatment)
- Unmodified Jurkats: 0.8276 (n.s.)
- IL-10-driven CAR circuit: 0.3669 (n.s.)
- Constitutive CAR expression: 0.8575 (n.s.)
- NFAT reporter only: 0.8705 (n.s.)

(Jurkat cell line: *p* value for comparison between untreated cells and co-culture with SKOV3 cells)
- Unmodified Jurkats: 0.3935 (n.s.)
- IL-10-driven CAR circuit: 0.0097
- Constitutive CAR expression: 0.0011
- NFAT reporter only: 0.3822 (n.s.)

(Jurkat cell line: *p* value for comparison between co-culture with SKOV3 cells and co-culture with SKOV3 cells and treatment with treatment with 250 ng/mL IL-10)
- Unmodified Jurkats: 0.7541 (n.s.)
- IL-10-driven CAR circuit: 0.0004
- Constitutive CAR expression: 0.7858 (n.s.)
- NFAT reporter only: 0.7953 (n.s.)

(Jurkat cell line: *p* value for comparison between co-culture with SKOV3 cells and IL-10-secreting SKOV3 cells)
- Unmodified Jurkats: 0.0682 (n.s.)
- IL-10-driven CAR circuit: 0.0115
- Constitutive CAR expression: 0.5669 (n.s.)
- NFAT reporter only: 0.0136

Below are the outcomes of two-tailed Welch's *t*-tests followed by BH procedure. Null hypotheses were that there existed no effect of PMA or antibody treatment for T cell activation on the measured NFAT reporter expression.

NFAT reporter assay in **Supplementary Figure 8c**
(Cell line, treatment: *p* value for comparison between untreated cells and each treatment)

- Unmodified Jurkats, 25 ng/mL PMA: 0.0824 (n.s.)
- Unmodified Jurkats, 100 ng/mL PMA: 0.4789 (n.s.)
- Unmodified Jurkats, Soluble CD3 and CD28 antibodies: 0.0688 (n.s.)
- Unmodified Jurkats, Immobilized CD3 and soluble CD28 antibodies: 0.4227 (n.s.)
- Jurkats with engineered NFAT reporter, 25 ng/mL PMA: 0.3739 (n.s.)
- Jurkats with engineered NFAT reporter, 100 ng/mL PMA: 0.0582 (n.s.)
- Jurkats with engineered NFAT reporter, Soluble CD3 and CD28 antibodies: 0.0056
- Unm Jurkats with engineered NFAT reporter Jurkats, Immobilized CD3 and soluble CD28 antibodies: 0.0016

Below are the outcomes of two-tailed Welch's *t*-tests followed by BH procedure. Null hypotheses were that there existed no effect of exogenous ligand treatment on the measured reporter expression or percentage of cells expressing reporter.

Functional signaling assay in **Supplementary Figure 7q**
(Cell line, treatment with or without IL-10: *p* value for comparison of reporter expression with and without NaB)

- Cells with reporter only, no IL-10: 0.5832 (n.s.)
- Cells with reporter only, IL-10: 0.3070 (n.s.)
- Cells with reporter and receptors, no IL-10: 0.0142
- Cells with reporter and receptors, IL-10: 0.0005

Functional signaling assay in **Supplementary Figure 7q**
(Cell line, treatment with or without IL-10: *p* value for comparison of percentage of cells expressing reporter with and without NaB)

- Cells with reporter only, no IL-10: n/a (both conditions in this comparison had 0 cells expressing reporter across all replicates (i.e. with 0 error)).
- Cells with reporter only, IL-10: 0.4226 (n.s.)
- Cells with reporter and receptors, no IL-10: 0.0034
- Cells with reporter and receptors, IL-10: 0.0004

Below are the outcomes of two-tailed Welch's *t*-tests followed by BH procedure. Null hypotheses were that there existed no effect of transfected soluble transcription factor plasmid on the measured expression level of different hybrid promoter architectures.

Functional signaling assay in **Supplementary Figure 9c**
(Promoter architecture, transfected transcription factor plasmid: *p* value for comparison vs. transfected empty vector)

- P1, ZF1: 0.000392
- P1, ZF6: 0.00016
- P1, ZF1+ZF6: 0.00209
- P2, ZF1: 0.000833
- P2, ZF6: 0.00255
- P2, ZF1+ZF6: 0.00149
- P3, ZF1: 0.0107
- P3, ZF6: 0.00128
- P3, ZF1+ZF6: 0.000182

- P4, ZF1: 0.0143
- P4, ZF6: 0.00523
- P4, ZF1+ZF6: 0.000161

Below are the outcomes of two-tailed Welch's *t*-tests followed by BH procedure. Null hypotheses were that there existed no effect of exogenous IL-10 treatment, co-transfected IL-10, co-transfected synTF, or co-transfected empty vector on the measured expression level of mbIL-15 by western blot analysis.

Functional signaling assay in **Supplementary Figure 8b**
(Transfected or genomically integrated circuit, ligand treatment or transfection: *p* value for comparison of expression level between untreated/treated conditions or between transfected with empty vector and transfected with IL-10 or synTF conditions)

- Transfected circuit, co-transfected IL-10 vs. co-transfected empty vector: 0.0219
- Transfected circuit, co-transfected synTF vs. co-transfected empty vector: 0.0150
- Genomically-integrated circuit, untreated cells vs. treatment with 250 ng/mL IL-10: 0.0343
- Genomically-integrated circuit, co-transfected IL-10 vs. co-transfected empty vector: 0.0203
- Genomically-integrated circuit, co-transfected synTF vs. co-transfected empty vector: 0.0077

Below are the outcomes of two-tailed Welch's *t*-tests followed by the Benjamini-Hochberg (BH) procedure. Null hypotheses were that there existed no effect of co-expressed ligand expression on the measured reporter expression.

Functional signaling assay in **Supplementary Figure 10e**
(NTEVp ECD-TMD, CTEVp ECD-TMD: *p* value for comparison no TNF vs. co-expressed TNF)

- NTEVp R1-R1, CTEVp R1-R1: 0.0078
- NTEVp R1-R1, CTEVp R2-R2: 0.2951 (n.s.)
- NTEVp R1-R1, CTEVp CD28-R1: 0.0362
- NTEVp R1-R1, CTEVp CD28-R2: 0.2550 (n.s.)
- NTEVp R2-R2, CTEVp R1-R1: 0.0561 (n.s.)
- NTEVp R2-R2, CTEVp R2-R2: 0.0192
- NTEVp R2-R2, CTEVp CD28-R1: 0.3708 (n.s.)
- NTEVp R2-R2, CTEVp CD28-R2: 0.4972 (n.s.)
- NTEVp CD28-R1, CTEVp R1-R1: 0.1415 (n.s.)
- NTEVp CD28-R1, CTEVp R2-R2: 0.7466 (n.s.)
- NTEVp CD28-R1, CTEVp CD28-R1: 0.4761 (n.s.)
- NTEVp CD28-R1, CTEVp CD28-R2: 0.0512 (n.s.)
- NTEVp CD28-R2, CTEVp R1-R1: 0.9175 (n.s.)
- NTEVp CD28-R2, CTEVp R2-R2: 0.9184 (n.s.)
- NTEVp CD28-R2, CTEVp CD28-R1: 0.0107
- NTEVp CD28-R2, CTEVp CD28-R2: 0.0127

**Supplementary Note 4.** ***Conversion of TNFR***

Tumor necrosis factor (TNF) is an immunostimulatory cytokine that is involved in inflammatory responses and is upregulated locally and systemically in most chronic inflammatory diseases. Targeting TNF with a cell-based therapy could permit sensing and responding to local sites of inflammation with immunosuppression, which could help manage chronic inflammatory disease without systemically immunosuppressing the patient and increasing their risk of infection and malignancy. TNF is a homotrimeric ligand that signals through TNFR1 or TNFR2, which are part of the TNF cytokine receptor class, and each activates different signaling pathways[8,9]. TNF can be found in the human body in membrane-tethered form (mTNF) or soluble form (sTNF)[9]. TNFR1 has higher affinity for TNF and can sense either mTNF or STNF, while TNFR2 has lower affinity and mainly senses mTNF with only weak interactions with sTNF[10]. Both receptors have similar mechanisms of signaling–TNFRs pre-associate in the absence of ligand with each other in a homo-associative manner as these associations are mediated by various domains in the ECD itself and the stalk regions[11-13]. Ligand binding induces a conformational change in these complexes as well as higher order homo-associations with other receptor complexes to transduce signaling (**Supplementary Figure 10a**)[14-17]. We hypothesized that this high order clustering mechanism might be amenable to split TEVp reconstitution when receptors with matching ECDs are paired. We also hypothesized that we could take advantage of smaller clusters of pre-associated receptors with the same ECD to sequester the MESA signaling components separately and drive heteroassociation upon ligand addition.

To characterize TNFR NatE MESA receptors, we began by evaluating expression. We observed strong surface expression of all variants tested, and a few design choices impacted surface expression (**Supplementary Figure 10b**). First, variants with a TNFR2 ECD were generally more highly expressed on the surface and on a whole cell basis than variants with a TNFR1 ECD (**Supplementary Figure 10c**). Secondly, variants with the respective native TMD were more highly expressed than variants with a CD28 TMD. Because signal sequence had no effect on surface expression, we proceeded to functionally evaluate only variants with the respective native signal sequence for each ECD. To use a setup in which ligand is co-expressed with receptors, we first validated bioactivity of three different ligand expression systems by expressing them in an engineered HEK293FT cell line that contains a previously validated genomically integrated NF-κB inducible reporter to ensure that trimers could form and initiate signaling through endogenous receptors (including TNFR1) expressed by HEK293FTs (**Supplementary Figure 10d**). We selected the TNF expression plasmid that drove the highest amount of reporter expression and co-transfected it along with plasmids encoding pairwise combinations of each ECD-TMD-signaling domain variant (**Supplementary Figure 10e**). We observed minimal signaling by all receptor pairs that included at least one receptor with a native TMD, and we observed higher amounts of signaling by receptors pairs in which both receptors contained a CD28 TMD. In particular, we did observe that a number of the TNFR2/TNFR2 pairings produced a ligand-dependent increase in reporter expression that was most pronounced when both receptors contain the CD28 TMD (**Supplementary Figure 10e,l**). These results suggested that both the ligand-unbound and ligand-bound states of receptors with native TNFR TMDs did not permit split TEVp reconstitution and signaling, while the CD28 TMD does permit signaling, though with high levels of ligand-independent background. We verified that this lack of signaling was not due to something about the native receptor TMD/JMD structure that caused an inability for reconstituted TEVp to cleave its recognition sequence (**Supplementary Figure 10f**). We also verified that proteolytic cleavage of TNFR1 by matrix metalloproteinases was not responsible for minimal signaling (**Supplementary Figure 10g-h**). We hypothesized that changing the split TEVp mutants to be variants that reconstitute more easily might permit more signaling by native TMD-bearing receptors. We indeed found that making it easier for split TEVp components to reconstitute conferred high levels of reporter expression (both background and induced signal increased) and produced some moderately-inducible receptors (**Supplementary Figure 10i-j**). This inducibility was independent of ECD pairing and was also observed in extended culture receptor transfection assays with recombinant, exogenous ligand (**Supplementary Figure 10l**). Similarly, we found that removing the need for reconstitution entirely by employing the MESA *trans*-cleavage mechanism also conferred inducible signaling with high background for some ECD pairings (**Supplementary Figure 10k**).

Overall, conversion of TNFRs to NatE MESA receptors yielded inducible, though low-performing receptors. Employing native transmembrane domains without split protease tuning only yielded inducible pairs with the CD28 TMD, so background signaling was high. Protease tuning yielded more inducible receptors, although background was also high. Together, these results suggest that the state change by which TNFRs signal is much less conducive to a split TEVp reconstitution and cleavage mechanism than other NatE MESA receptors explored here. It is possible that TNFRs associate in a way that does not promote split TEVp reconstitution, and signal observed with split TEVp mutants and *trans*-cleavage receptors is mainly determined by transient receptor encounters that change as a function of receptor cluster size. This conclusion could be supported by findings that constitutively active TNFR mutants have C-terminal domains that are held further apart in the ligand-bound state[18]. While conversion of TNFR into a TNF-inducible synthetic biosensor was successful, the conversion process did not yield a high performing biosensor.

**Supplementary Note 5.** ***Conversion of TGF-βR***

Transforming growth factor β (TGF-β) is a multifunctional cytokine that has multiple (sometimes opposing) functions depending on its context. TGF-β generally plays an immunosuppressive role, downregulating effector functions and expansion of many immune cell types[19]. These properties also make TGF-β a tumor suppressor, halting growth of pre-malignant cells. Throughout cancer development, however, TGF-β supports metastasis by suppressing immune surveillance and cytotoxic activity in the tumor microenvironment[20]. As a result, targeting TGF-β with a cell-based therapy that drives customized transcriptional output could enable tumor targeting of cytotoxic functions. TGF-β, in its mature form, is a homodimeric ligand that signals through TGF-βR1 and TGF-βR2, which are serine/threonine kinase receptors that drive SMAD signaling pathways[20]. A TGF-βR2 pair first binds to TGF-β, and then TGF-βR1 is recruited to form a heterotetrameric complex (**Supplementary Figure 11a**). Both receptor types are generally found as monomers in the absence of ligand, but TGF-βR1 can be found partially pre-dimerized when overexpressed[21,22]. The monomeric nature of these receptors in the absence of ligand and the ligand-mediated heteroassociation are both attractive features for conversion into a ligand-induced split TEVp-reconstitution-based mechanism.

We first evaluated surface and whole cell expression of TGF-β NatE MESA receptor variants. We found that compared to the other natural receptors explored, TGF-βR-based receptor chains were not as well expressed on the cell surface across all tested variants (**Supplementary Figure 11b**). TGF-βR1-containing receptor chains were more highly expressed than TGF-βR2-containing receptor chains when native signal sequences were employed, but this flipped when alternative signal sequences were employed, especially the CD8a signal sequence (**Supplementary Figure 11c**). We proceeded to functionally evaluate the variants with the CD8a signal sequence because they were best expressed on the surface. We evaluated receptor performance when co-expressed with TGF-β1 (the mature protein sequence, aa279-390, with a human IgK light chain secretion sequence). We observed that most receptor pairs showed no inducibility (background and induced reporter output were similar), while some were de-inducible (presence of ligand resulted in significantly less reporter output) (**Supplementary Figure 11d**). Of note, we found that receptor pairs with matching TGF-βR1 ECDs produced higher magnitudes of signal with and without ligand, which agrees with reports that this receptor can homodimerize when highly expressed, as is the expected expression regime in this experiment[21,22]. We also found that some pairings with both receptor chains containing CD28 TMDs also produced increased levels of background and induced signal, though much lower than what was observed with other NatE MESA receptors. We were surprised to see that none of the mixed pairs of ECDs yielded ligand-induced signaling and that none of the matched TGF-βR2 ECDs yielded ligand-induced signaling. Because no receptor pairings showed a ligand-induced increase in signaling and overall low levels of signaling, we verified that this was not due to something about the native receptor TMD/JMD structure that caused an inability for reconstituted TEVp to cleave its recognition sequence (**Supplementary Figure 11e**).

Based on these results and because expression level and particularly surface expression of these receptors was low compared to other NatE MESA receptors characterized in this study, we decided to evaluate if our designs had removed critical parts of the JMD that are required for expression. This hypothesis was motivated by the availability of a commercial cell line for detection of TGF-β using chimeric receptors with TGF-βR ECDs (PathHunter express TGFβR1/TGFβR2 dimerization assay, Eurofins DiscoverX), which used a slightly longer JMD for both ECD types to reconstitute an enzyme that can produce a chemiluminescent signal. We also hypothesized that alternative isoforms of the TGF-βR ECDs could improve performance. In particular, we noted that the isoforms we originally selected *should* bind TGF-β1 without additional cofactors, but that the TGF-βR2 isoform 2 is a better binder of the other TGF-β isoforms and may have advantageous properties in our system[23]. For TGF-βR1, extension of the JMD from 5 to 11 amino acids had no impact on surface expression (**Supplementary Figure 11f**). For TGF-βR2, we found that the switch from isoform 1 to 2 alone had minimal effect on surface expression, but when combined with JMD extension from 5 to 11 amino acids or a CD28 TMD, receptors bearing TGF-βR2 isoform 2 were the best surface-expressed receptors across NTEVp and CTEVp chains (**Supplementary Figure 11f**). When functionally evaluated with co-expressed TGF-β, no pairs demonstrated a ligand-inducible increase in reporter expression

(**Supplementary Figure 11g-h**). We also tested a variety of TGF-β1 ligand expression systems including the mature TGF-β1 sequence with a human IgK light chain secretion sequence, the mature TGF-β1 sequence with a mouse IgK light chain secretion sequence, and the pre-processed proTGF-β1 sequence with its native signal sequence and latency associated peptide (LAP). No ligand expression system yielded a ligand-induced increase in reporter expression, though the proTGF-β1 ligand induced some significant decreases in reporter expression (**Supplementary Figure 11i**). Overall, we conclude that the native TGF-βR1/2 binding geometry is not conducive to split protease reconstitution and cleavage.

Finally, we next chose to investigate how employing various CD28 TMD lengths and JMDs might overcome geometric constraints with native TGF-βR1/2 TMDs to yield inducible receptor pairs. We evaluated how switching from a truncated mouse to a truncated human CD28 TMD, extending the human CD28 TMD to be full-length, and including a human CD28 JMD impact expression and resulting signaling by CD28-bearing receptors. Overall, we found that these extensions minimally impacted surface expression, (**Supplementary Figure 11j**). Functional performance was not improved because no receptor pairs were inducible and ectodomain choice was still the most important determinant in magnitude of signaling (**Supplementary Figure 11k**). These results suggested that ligand-mediated assembly of receptor heterotetramers and TGF-βR1 or TGF-βR2 dimers is not conducive to split TEVp reconstitution and biosensor conversion. Overall, conversion of TGF-βR to a synthetic biosensor was unsuccessful in the chosen design space.

**Supplementary Note 6.** ***Evaluation of strategies for generating stable biosensor cells and impacts of genomic context on performance.***

To better understand which cells are signaling within the engineered cell population and guide future implementations of these NatE MESA receptors, we sorted cells using two different strategies. First, we sorted cells based on the magnitude of constitutive fluorescent protein mNeonGreen expressed as a proxy for how many copies of the transposon were stably integrated (**Supplementary Figure 7i**). We broke the top half of the population, which is where all of the reporter positive cells exist, into four sections (referred to as octiles of the full population). Cells with higher mNeonGreen expression exhibited higher background and induced reporter expression, and fold induction was relatively constant across these populations (**Supplementary Figure 7j,k**). Interestingly, in general, the addition of ligand only moderately increased the percent of cells exhibiting signaling (reporter-positive), suggesting that addition of ligand does not make reporter-negative cells become reporter-positive. Instead, addition of ligand increases reporter output of already reporter positive cells (**Supplementary Figure 7l**). We hypothesized that cells with nonzero background (reporter positive before ligand addition) are least likely to silence the reporter gene and are responsible for most of the signaling within the population[24]. The second sort strategy we employed was designed to evaluate whether cells with no background signal were capable of signaling, and if so, if a low background sort strategy could result in a higher performing population based on fold induction. We again looked at the top four octiles based on mNeonGreen expression but also restricted gating to include only reporter negative cells (**Supplementary Figure 7m**). We found that cells in all four octiles were still capable of signaling, and again, the magnitude of signal decreased as mNeonGreen expression decreased (**Supplementary Figure 7n,o**). We again observed only moderate increases in percent of cells signaling upon ligand addition and the overall percentages were much lower suggesting that there are a higher proportion of non-signaling cells in the reporter negative cell population (**Supplementary Figure 7p**). To evaluate if silencing was limiting the proportion of reporter-positive cells, we employed sodium butyrate on the parent (pre-sort population) and found that percent of signaling cells increased, along with the magnitude of both background and induced signal (**Supplementary Figure 7q**). Interestingly, across all of these strategies that impact genetic context (sorting based on transposon copy number via mNeonGreen expression, sorting based on background reporter, treatment with a chromatin-modifying drug, implementation via two different transposon designs), we found that fold induction was relatively conserved, suggesting that our functional assays are capturing performance characteristics of the receptors themselves, independent of context (**Supplementary Figure 7r**). This suggests that the IL-10 biosensor is robust to genetic context and resulting expression level.

**Supplementary Note 7. *Definition of synergy throughout this study.***

To evaluate logical AND function performance in this study, we quantified synergy of each design. An AND gate is defined by exhibiting synergy if the output when both inputs are present is larger than the sum of the outputs in response to each individual input. Explicit definitions of synergy metrics for relevant figure panels are listed below. If a synergy metric is greater than 1, the AND function is synergistic.

In **Figure 3e**, synergy is used to evaluate T cell activation via NFAT reporter output with inputs of IL-10 (soluble; administered exogenously with recombinant IL-10 or secreted by engineered, co-cultured SKOV3 cells) and HER2 (surface-bound; present on wildtype, co-cultured SKOV3 cells). Synergy is defined in two ways represented by **Equations 1,2**:

$$Synergy = \frac{(NFAT\ output\ with\ wt\ SKOV3\ cells\ and\ rIL10) - (NFAT\ output\ with\ no\ treatment)}{(NFAT\ output\ with\ wt\ SKOV3\ cells) + (NFAT\ output\ with\ rIL10)}$$

**Equation 1**

$$Synergy = \frac{(NFAT\ output\ with\ IL10{-}secreting\ SKOV3\ cells) - (NFAT\ output\ with\ no\ treatment)}{(NFAT\ output\ with\ wt\ SKOV3\ cells) + (NFAT\ output\ with\ rIL10)}$$

**Equation 2**

In **Figure 4d, Figure 4f,** and associated supplementary figures, synergy is used to evaluate reporter output with inputs of co-expressed IL-10 and co-expressed VEGF. Synergy is defined by **Equation 3**:

$$Synergy = \frac{(Background\ subtracted\ reporter\ output\ with\ co{-}expressed\ VEGF\ and\ IL10)}{(Background\ subtracted\ reporter\ output\ with\ co{-}expressed\ VEGF) + (Background\ subtracted\ reporter\ output\ with\ co{-}expressed\ IL10)}$$

**Equation 3**

**Supplementary Note 8. *Generalized design rules and workflow for conversion of natural receptors to NatE MESA receptors***

To aid in future conversions of natural receptors into NatE MESA receptors, we propose a set of characteristics and design rules to follow. This information is also depicted in **Supplementary Figure 12**.

First, characterizing surface and whole cell expression by immunohistochemistry and western blotting is essential to confirm proper receptor folding, maturation and trafficking, though for a high throughput conversion, this might be characterized only for functional hits. Next, testing receptor functionality with co-expressed ligand in the transient context can help identify receptor combinations that do not yield favorable interactions. Especially for cases in which surface expression characterizations are skipped, using co-expressed ligand removed the requirement for surface expression and still enables functional testing. Finally, testing the performance of successful receptors in the stable expression context is crucial to characterize response to exogenous ligand in a translationally relevant context. Transposon integration was suitable for this application given it allows for high expression levels of large DNA cargo. Transfection-based assays to detect exogenous ligand require careful tweaking of timing to allow for receptor buildup and reporter buildup, which is challenging to balance with the sensitive state of cells after transfection. This may be helped by the use of reagents to halt growth and boost protein production, such as valproic acid. Further characterization of receptors such as ligand dose response and microscopy studies are informative but not essential. Additionally, the strategies explored here to mitigate gene silencing with sodium butyrate treatment and cell sorting can also be helpful from an application standpoint, but do not change overall conclusions about receptor performance in the genomic context. As for the receptor design space, it is more important to test multiple TMD choices rather than signal sequences when first trying to identify a functional receptor pair because favorable geometry of the signaling domains in the ligand-unbound and ligand-bound states is required. Signal sequence can be used to improve surface expression or general expression but will not change inducibility (i.e. break or rescue receptor function in response to ligand).

**Supplementary Note 9.** ***List of acronyms used in this study***

| | |
|---|---|
| ANOVA | Analysis of variance |
| AP-1 | Activator protein-1 |
| APC | Allophycocyanin |
| AU | Arbitrary units |
| BATF | Basic leucine zipper transcription factor ATF-like |
| BCA | Bicinchoninic acid assay |
| BlastR | Blasticidin resistance gene |
| BSA | Bovine serum albumin |
| CAR | Chimeric antigen receptor |
| CD28 | Cluster of differentiation 28 |
| CD8a | Cluster of differentiation 8 |
| cHS4 | 5'-HS4 chicken β-globin insulator |
| CMV | Cytomegalovirus (promoter) |
| CTEVp | C-terminal protein component split TEVp |
| DMEM | Dulbecco's Modified Eagle Medium |
| DMSO | Dimethyl sulfoxide |
| DNA | Deoxyribonucleic acid |
| ECD | Extracellular domain, ectodomain |
| ECL | Enhanced chemiluminescence |
| EDTA | Ethylenediaminetetraacetic acid |
| FACS | Fluorescence activated cell sorting |
| FBS | Fetal bovine serum |
| FI | Fold induction or fluorescence intensity |
| FSC-A | Area in the Forward Scatter channel |
| FSC-H | Height in the Forward Scatter channel |
| GMFI | Geometric mean of fluorescence intensity |
| hEF1a | Human elongation factor-1 alpha (promoter) |
| HEPES | 4-(2-hydroxyethyl)-1-piperazineethanesulfonic acid |
| hIgG VH | Human immunoglobulin heavy chain variable domain |
| HSD | Tukey's honest significance test |
| hSS | Human signal sequence, specifically derived from the immunoglobulin kappa light chain |
| HygroR | Hygromycin resistance gene |
| hyPBase | Hyperactive Piggy Bac transposase |
| ICD | Intracellular domain |
| IgG | Immunoglobulin G |
| IL-10 | Interleukin 10 |
| IL-10Ra/b | Interleukin 10 receptor alpha/beta |
| Int | Intein |
| IRES | Internal ribosome entry site |
| ITR | Inverted terminal repeat |
| JMD | Juxtamembrane domain |
| LB | Luria-Bertani |
| LP | Landing pad |
| LTR | Long terminal repeat |
| MEA750s | Molecules of Equivalent Alexa-750 |
| MEBFPs | Molecules of Equivalent blue fluorescent protein |
| MEFLs | Molecules of Equivalent Fluorescein |

| | |
|---|---|
| MEPTRs | Molecules of Equivalent PE-Texas Red |
| MESA | Modular extracellular sensor architecture |
| MFI | Mean fluorescence intensity (in arbitrary units) |
| MGEV | Multiple gene expression vector |
| miRFP720 | Monomeric infrared fluorescent protein (with 720 nm emission) |
| mSS | Mouse signal sequence, specifically derived from the immunoglobulin kappa light chain |
| mNG | Monomeric neon green fluorescent protein |
| mTB2 | Monomeric tag blue fluorescent protein |
| Mut | Mutant |
| NaB | Sodium butyrate |
| NatE MESA | Natural ectodomain modular extracellular sensor architecture |
| natTF | Natural transcription factor |
| NEB | New England Biolabs |
| NES | Nuclear export signal |
| NFAT | Nuclear factor of activated T cells |
| NF-kB | Nuclear factor kappa light chain enhancer of activated B cells |
| NLS | Nuclear localization signal |
| NTEVp | N-terminal protein component of split TEVp |
| PBS | Phosphate-buffered saline |
| PCR | Polymerase chain reaction |
| PDB | Protein Data Bank |
| PRS | Protease recognition sequence (for TEVp) |
| PuroR | Puromycin resistance gene |
| PVDF | Polyvinylidene fluoride |
| Rapalog, Rapa | Rapamycin analog, here specifically Takara AP21967 |
| RCP | Rainbow calibration particles |
| Rep | Reporter |
| REU | Rosetta Energy Units |
| RIPA | Radioimmunoprecipitation assay buffer |
| RNA | Ribonucleic acid |
| RNase | Ribonuclease that degrades RNA |
| RTK | Receptor tyrosine kinase |
| S.E.M. | Standard error of the mean |
| SB100X | Hyperactive Sleeping Beauty transposase |
| scFv | Single chain variable fragment |
| SDS | Sodium dodecyl sulfate |
| SS | Signal sequence |
| SSC-A | Area in the Side Scatter channel |
| synTF | Synthetic transcription factor |
| Tbet | T-box expressed in T cells |
| TBS | Tris-buffered saline |
| TBST | Tris-buffered saline with Tween |
| TEVp | Tobacco etch virus protease |
| TF | Transcription factor |
| TGF-β | Transforming growth factor beta |
| TGF-βR1/2 | Transforming growth factor beta receptor 1/2 |
| TMD | Transmembrane domain |
| TNF | Tumor necrosis factor |
| TNFR1/2 | Tumor necrosis factor receptor 1/2 |

| | |
|---|---|
| TUPV | Transcriptional unit positioning vector |
| URCP | Ultra-rainbow calibration particles |
| VEGF | Vascular endothelial growth factor |
| VEGFR1/2 | Vascular endothelial growth factor 1/2 |
| WT | Wildtype |
| ZF | Zinc finger |
